# Increasing Cancer Stemness Drives Prostate Cancer Progression, Plasticity, Therapy Resistance and Poor Patient Survival

**DOI:** 10.1101/2025.04.27.650697

**Authors:** Xiaozhuo Liu, Anmbreen Jamroze, Eduardo Cortes, Yibing Ji, Kathy Zhao, Faming Zhao, Julian Ho, Yang Liu, Elai Davicioni, Shan Wu, Wen (Jess) Li, Felix Y. Feng, Joshi J. Alumkal, Daniel E. Spratt, Christopher J. Sweeney, Han Yu, Qiang Hu, Cheng Zou, Dingxiao Zhang, Kevin Lin, Yue Lu, Gurkamal Chatta, Kent L. Nastiuk, David W. Goodrich, Kiera Rycaj, Jason S. Kirk, Igor Puzanov, Zheng Xia, Song Liu, Jianmin Wang, Dean G. Tang

**Affiliations:** Department of Pharmacology and Therapeutics, Roswell Park Comprehensive Cancer Center, Buffalo, NY 14263, USA; Department of Biostatistics and Bioinformatics, Roswell Park Comprehensive Cancer Center, Buffalo, NY 14263, USA; Department of Biostatistics, University at Buffalo, Buffalo, NY 14263, USA; Veracyte Inc, San Diego, CA 92121, USA; Department of Biomedical Engineering, The University of Texas at Austin, Austin, TX 78712, USA; Department of Radiation Oncology, University of California, San Francisco, CA 94143, USA; Department of Internal Medicine, Department of Hematology and Oncology, Rogel Cancer Center, University of Michigan, Ann Arbor, MI 48109, USA; University Hospitals Seidman Cancer Center and Case Western Reserve University, Cleveland, Ohio, USA; South Australian Immunogenomics Cancer Institute, University of Adelaide, Adelaide, SA 5000, Australia; School of Biomedical Sciences, Hunan University, Changsha, China; Department of Epigenetics and Molecular Carcinogenesis, the University of Texas M.D Anderson Cancer Center, 1515 Holcombe Blvd., Houston, TX 77030, USA; Department of Medicine, Roswell Park Comprehensive Cancer Center, Buffalo, NY 14263, USA; Department of Cancer Genetics, Roswell Park Comprehensive Cancer Center, Buffalo, NY 14263, USA; Experimental Therapeutics Graduate Program, University at Buffalo and Roswell Park Comprehensive Cancer Center, Buffalo, NY 14263, USA

**Author notes:** Corresponding authors: Xiaozhuo Liu, PhD, Department of Pharmacology and Therapeutics, Roswell Park Comprehensive Cancer Center, Buffalo, NY 14263., Song Liu, PhD, Department of Biostatistics and Bioinformatics, Roswell Park Comprehensive Cancer Center, Buffalo, NY 14263., Jianmin Wang, PhD, Department of Biostatistics and Bioinformatics, Roswell Park Comprehensive Cancer Center, Buffalo, NY 14263., Dean Tang, MD, PhD, Department of Pharmacology and Therapeutics, Roswell Park Comprehensive Cancer Center, Buffalo, NY 14263. These authors contributed equally to this work (Xiaozhuo Liu, Anmbreen Jamroze, Eduardo Cortes). We dedicate this work to the memory of Dr. Felix Y. Feng, whose contributions to this project and the field of prostate cancer research are immeasurable.

**Keywords:** Prostate cancer, stemness, AR signaling, heterogeneity, plasticity, aggressiveness, *RB1* loss, MYC

## Abstract

**Background:** Cancer progression is often accompanied by dedifferentiation and acquisition of stem cell-like properties (stemness). In prostate cancer (PCa), lineage plasticity and therapy resistance remain major clinical challenges, yet a unified quantitative transcriptomic framework connecting stemness, androgen receptor (AR) signaling, castration resistance, and disease progression across the PCa continuum is lacking.

**Methods:** We integrated 87,339 transcriptomic profiles from 33 preclinical and clinical datasets spanning the PCa continuum from normal prostate and treatment-naïve primary PCa (Pri-PCa) to PCa treated with neoadjuvant ADT (nADT) and metastatic castration-resistant PCa (mCRPC), with single-cell RNA-seq analyses encompassing 115,197 cells. Cancer stemness was quantified using a transcriptome-derived mRNA-based Stemness Index (mRNAsi; hereafter Stemness), and a 12-gene PCa-Stem signature was developed to capture PCa-specific stemness. Stemness, PCa-Stem, canonical AR activity (c_AR-A), castration-reprogrammed AR activity (cr_AR-A), *RB1*-loss, *PTEN*-loss, and MYC activity signatures were quantified across cohorts. Functional validation included MYC inhibition and representative PCa-Stem signature gene depletion in PCa models. Clinical prognostic significance was evaluated in independent patient cohorts.

**Results:** The Stemness score and c_AR-A increased concordantly during early prostate tumorigenesis but diverged with PCa progression: as Gleason grade increased, c_AR-A declined while Stemness continually increased. mCRPC exhibited the highest Stemness and lowest c_AR-A, a pattern recapitulated in *Pten*/*Rb1*/*Trp53*-deficient mouse models. Global Stemness increased progressively across the PCa continuum, was enriched in aggressive PAM50-LumB and PCS1 subtypes, associated with proliferative and lineage plasticity programs, and predicted poor patient survival. The newly derived 12-gene PCa-Stem signature provided a PCa-specific molecular representation of Stemness and tracked disease progression and poor patient survival. Network analyses identified a coordinated mitotic regulatory program linking MYC activity, *RB1*-loss, cr_AR-A, and the PCa-Stem signature. Spatial and single-cell transcriptomic analyses localized the PCa-Stem program to lineage plasticity-related epithelial cells and demonstrated progressive expansion of PCa-Stem⁺ epithelial cells during PCa progression. Functional perturbation of representative PCa-Stem signature genes, as well as genetic and pharmacological MYC inhibition, consistently suppressed Stemness-associated phenotypes in diverse PCa models.

**Conclusions:** Cancer Stemness quantitatively captures PCa aggressiveness, lineage plasticity, treatment resistance, disease progression, and poor patient survival. cr_AR-A, *RB1* loss, and MYC activation cooperate to reinforce the high-Stemness state and therapy resistance in mCRPC. Collectively, our work establishes a trajectory-integrated transcriptomic framework defining cancer Stemness as a quantifiable molecular and clinical determinant of PCa aggressiveness, lineage plasticity, disease progression, therapy resistance and patient survival.

## Background

Malignant transformation and cancer progression are characterized by progressive loss of differentiation (or oncogenic dedifferentiation) and gain of stem cell-like traits (i.e., stemness). Poorly differentiated tumors are enriched in gene expression profiles of normal embryonic and adult stem cells and in cancer stem cells [1–5]. However, the oncogenic dedifferentiation-based stemness concept [6] that prostate cancer (PCa) represents abnormal organogenesis [7, 8] has not been systematically addressed and there lack quantitative measurements of the overall stemness-associated aggressiveness in PCa.

PCa is projected to claim the lives of >36,000 American men in 2026 [9]. PCa incidence rates increased 3% per year from 2014 - 2019 driven by advanced PCa diagnosis and the proportion of men diagnosed with metastases has doubled from 2011 to 2019 when the next generation of androgen receptor (AR) pathway inhibitors (ARPIs) such as abiraterone (Zytiga; FDA approval in 2011) and enzalutamide (Xtandi; FDA approval in 2012) were introduced to the clinic. Significant increases in diagnosis of aggressive and advanced disease and continued high PCa mortality have been linked to therapy resistance driven by intrinsic PCa cell heterogeneity (e.g., in AR expression) and treatment-induced plasticity [8].

The level of malignancy of primary PCa (Pri-PCa) has been traditionally assessed by pathologists using a combined Gleason Score (GS), which gauges histopathological differentiation pattern of prostatic glands. The higher the GS, the more undifferentiated and more malignant the Pri-PCa is. The GS system, although clinically the ‘gold’ standard in pinpointing PCa diagnosis, is only semi-quantitative, can be subjective (to individual pathologist’s assessment), and is generally not applicable to treatment-failed tumors and distant metastases (which have mostly lost differentiated glandular structures). Notably, whether more advanced Pri-PCa (GS8-10), treatment failure and metastatic dissemination are quantitatively associated with increased cancer stemness remains unknown.

AR is the master regulator of normal prostatic epithelium differentiation and represents the therapeutic target of PCa treatments including androgen-deprivation therapy (ADT) and ARPIs. Inevitably, tumors develop resistance to ADT/ARPIs leading to the state of castration-resistant PCa (CRPC), which involves many mechanisms including genetic alterations of *AR* and key genes in AR pathway [10], lineage plasticity [8, 11, 12], and increased stemness [8, 13]. Under therapeutic pressure, AR binding shifts toward noncanonical targets (i.e., reprogramming of AR cistrome), resulting in extensive reconfiguration of the AR transcriptional program [14–17]. These observations suggest a progressive divergence of AR signaling from its canonical, pro-differentiation role during tumor evolution. Thus, quantifying canonical AR activity across disease stages should provide a tractable means to capture this adaptive trajectory. However, how such changes in AR signaling relate to global cancer stemness and tumor aggressiveness remains unclear.

These discussions raise the need for a quantitative framework that simultaneously captures AR signaling and stemness to reveal how their dynamic interplay contributes to PCa aggressiveness, plasticity, and therapy resistance. Here, we adapted and developed transcriptome-based gene signature scores to quantitatively assess the canonical AR activity (c_AR-A) and oncogenic stemness levels (Stemness score; Stemness for short) and their inter-relationship across the PCa spectrum. Specifically, we integrated 87,339 transcriptomic profiles from 24 clinical cohorts and 9 preclinical datasets, including bulk RNA-seq, single-cell RNA-seq, and spatial transcriptomic data, encompassing the PCa continuum from normal/benign prostate through early-and advanced-stage PCa, pre-and post-therapy PCa, and, finally, metastatic CRPC (mCRPC), to systematically track the dynamic evolution of AR activity and Stemness. We further developed a 12-gene PCa-Stem signature to capture PCa-specific stemness and investigated its biological underpinnings and clinical relevance. Our findings demonstrate that Stemness quantitatively tracks PCa progression and adverse clinical outcomes, establishing a trajectory-integrated, pan-cohort framework that defines pervasively increasing cancer Stemness as a quantitative hallmark of PCa aggressiveness, lineage plasticity, therapy resistance, and poor patient prognosis.

## Materials and Methods

### I. PCa datasets and data collection

In this study, we utilized a total of 33 published and in-house datasets, comprising 24 clinical cohorts and 9 preclinical datasets and encompassing 87,339 transcriptomic profiles across the evolutionary spectrum of PCa (Fig. S1A-B). Transcriptomic analyses included bulk RNA-seq, single-cell RNA-seq (scRNA-seq; 104 samples and 115,197 cells), and spatial transcriptomics, together with genomic and clinicopathologic data where available. The datasets encompassed normal prostate, tumor-adjacent benign tissue, treatment-naïve primary PCa (Pri-PCa) with increasing GS, PCa treated with neo-adjuvant ADT (nADT) or long-term ADT, mCRPC, and PCa cell lines, PCa patient-derived xenografts (PDX) and xenografts. Clinical information, genomic data and gene expression profiling data were downloaded and detailed information on each dataset is listed in Table S1. Dataset descriptions include dataset source references, PubMed IDs (PMIDs; RRID:SCR_004846), DOI links, sample type and source, sample sizes, data types used in the current study (clinicopathologic data, RNA-seq, microarray, whole exome sequencing [WES]), and public data access portals.

#### Data Access and Repository Information

Data for this study were retrieved from the following public repositories and databases:

1. GTEx Portal (RRID:SCR_013042; https://gtexportal.org/)
2. Gene Expression Omnibus (GEO; RRID:SCR_005012; https://www.ncbi.nlm.nih.gov/geo/)
3. cBioPortal for Cancer Genomics (RRID:SCR_014555; https://www.cbioportal.org/)
4. Zenodo Repository (RRID:SCR_004129; https://zenodo.org/)
5. Decipher GRID Database (RRID:SCR_006552; ClinicalTrials.gov identifier: NCT02609269)
6. European Genome-Phenome Archive (EGA; RRID:SCR_004944; https://ega-archive.org/)
7. European Nucleotide Archive (ENA; RRID:SCR_006515; https://www.ebi.ac.uk/ena/browser/home)
8. UCSC Xena Functional Genomics Portal (RRID:SCR_018938; https://xenabrowser.net/)
9. Single Cell Portal (RRID:SCR_014816; https://singlecell.broadinstitute.org/single_cell) Accession numbers and portal links for the 33 datasets described below are summarized in Table S1.

#### Brief Descriptions of the 33 Datasets

1. **GTEx 2013** (PMID: 23715323; DOI: 10.1038/ng.2653) [18]: The Genotype-Tissue Expression (GTEx) Project repository collected high-throughput and clinical data of normal tissue from many organs. RNA-seq data of 245 normal prostate tissue samples was downloaded from the GTEx portal (RRID:SCR_013042; https://www.gtexportal.org/, version: v8).
2. **Smith 2015** (PMID: 26460041; DOI: 10.1073/pnas.1518007112) [19]: RNA-seq data of 5 prostate basal cell (Trop2^+^CD49f^hi^) and corresponding 5 luminal cell (Trop2^+^CD49f^lo^) samples purified by fluorescence-activated cell sorting (FACS) from benign human prostate tissue in 5 PCa patients (n=5) was retrieved from the GEO database GSE82071.
3. **Liu 2016** (PMID: 27926864; DOI: 10.1016/j.celrep.2016.11.010) [20]: RNA-seq data of 3 FACS-purified human prostate basal cell (CD45^-^EpCAM^+^CD49f^hi^CD38^lo^) and corresponding luminal cell (CD45^-^EpCAM^+^CD49f^lo^CD38^hi^) and luminal progenitor cell (CD45^-^EpCAM^+^CD49f^lo^CD38^lo^) populations from the benign prostate tissues of 3 PCa patients (n=3) was retrieved from the supplementary data of the indicated reference. The microarray-based transcriptomic profiles of the same samples were retrieved from the GEO database GSE89050.
4. **Zhang 2016** (PMID: 26924072; DOI: 10.1038/ncomms10798) [21]: RNA-seq data of 3 human prostate basal cell (Trop2^+^CD49f^hi^) with the corresponding 3 luminal cell (Trop2^+^CD49f^lo^) samples FACS-purified from benign prostate tissues of 3 PCa patients (n=3) was retrieved from the GEO database GSE67070.
5. **TCGA PRAD 2015** [22] and **Pan-Cancer Atlas 2018**: The Cancer Genome Atlas (TCGA) (https://www.cancer.gov/tcga) repository collected high-throughput molecular and clinical data from primary cancer and matched normal/benign samples spanning 33 human cancer types, including prostate adenocarcinoma (PRAD). For each cancer type, TCGA published an initial “marker paper” summarizing analyses performed. Supplemental and associated data files supporting these “marker papers” are available through the NCI Genomic Data Commons (GDC) website (https://gdc.cancer.gov/about-data/publications). For PCa, **TCGA PRAD 2015** [22] (PMID: 26544944; DOI: 10.1016/j.cell.2015.10.025) is the marker paper on PCa published in 2015, which included the dataset of 333 primary prostate adenocarcinomas (281 treatment-naïve Pri-PCa, 49 PCa with post-surgery adjuvant ADT, and 52 matched normal/benign prostate tissue). Genomic data were obtained from **cBioPortal for Cancer Genomics** (RRID:SCR_014555; https://www.cbioportal.org), specifically the TCGA-PRAD Firehose Legacy study (https://www.cbioportal.org/study/summary?id=prad_tcga). RNA-seq expression data was obtained from Zhang *et al*. [23] (PMID: 32350277; DOI: 10.1038/s41467-020-15815-7). The expanded **TCGA PRAD 2018 (Pan-Cancer Atlas)** cohort included 494 patients with Pri-PCa (422 treatment-naïve Pri-PCa, 70 PCa with post-surgery adjuvant hormone treatment, 2 PCa with chemotherapy) and 52 matched adjacent normal/benign prostate tissue. Clinicopathologic features (age, Gleason scores, PSA levels, tumor stages) and genomic alterations data (mutation frequency, copy number alterations [CNAs], structural variants [SVs], tumor mutation burden [TMB], fraction of genome altered [FGA]) were retrieved from cBioPortal for Cancer Genomics PRAD PanCancer Atlas study (https://www.cbioportal.org/study/summary?id=prad_tcga_pan_can_atlas_2018) and from the Pan-Cancer Atlas companion site (https://www.cell.com/pb-assets/consortium/pancanceratlas/pancani3/index.html). Survival data were obtained from Liu *et al*. [24] (PMID: 29625055; DOI: 10.1016/j.cell.2018.02.052). Normalized RNA-seq data were downloaded via NCI GDC hub at UCSC Xena Functional Genomics Portal [25] (https://gdc.xenahubs.net, version 2019-07-20; PMID: 32444850; DOI: 10.1038/s41587-020-0546-8).
6. **Rajan 2014** (PMID: 24054872; DOI: 10.1016/j.eururo.2013.08.011) [26]: Clinicopathologic (Gleason scores) and RNA-seq data of 7 matched pre-and post-nADT PCa samples were obtained from Table 1 of the indicated reference and the GEO database GSE48403.
7. **Sharma 2018** (PMID: 30314329; DOI: 10.3390/cancers10100379) [27]: Clinicopathologic (Gleason scores) and RNA-seq data of 7 matched pre-and post-nADT PCa samples from the responder group (also defined as “Low Impact Group” in the source reference) were obtained from supplementary Table S1 of the indicated reference and the GEO database GSE111177.
8. **Long 2020** (PMID: 32951005; DOI: 10.1016/j.eururo.2013.08.011) [28]: RNA-seq data of 6 matched pre-and post-nADT PCa samples were obtained from the GEO database GSE82071. Additionally, RNA-seq data from 10 locally advanced PCa samples along with their paired adjacent benign prostate tissues were obtained from the GEO database GSE82071.
9. **Nastiuk (RPCI) 2020** (PMID: 41646678; DOI: 10.64898/2026.01.10.26343859) [29]: Clinicopathologic data (Gleason scores) and RNA-seq data of 43 Pri-PCa samples from post-nADT PCa patients and 43 tumor samples from age-, stage-, clinically-matched PCa patients without nADT were obtained from Drs. K. Nastiuk and G. Chatta at the Roswell Park Comprehensive Cancer Center.
10. **Gerhauser 2018** (PMID: 30537516; DOI: 10.1016/j.ccell.2018.10.016) [30]: RNA-seq data of 118 primary tumor samples and 9 adjacent benign prostate tissue from early-onset PCa patients were obtained from the European Genome-Phenome Archive (EGA) (https://ega-archive.org), accession number EGAS00001002923.
11. **Wyatt 2014** (PMID: 25155515; DOI: 10.1186/s13059-014-0426-y) [31]: Clinicopathologic (Gleason scores) and RNA-seq data of 12 primary PCa samples from treatment-naïve PCa patients and 12 adjacent benign prostate tissue were obtained from supplementary Table S1 of the indicated reference and the European Nucleotide Archive (ENA) (https://www.ebi.ac.uk/ena/browser/home), accession number PRJEB6530.
12. **Spratt 2017** (PMID: 28358655; DOI: 10.1200/JCO.2016.70.2811) [32]: Transcriptomic profiles (Microarray) and clinicopathologic data of 855 radical prostatectomy specimens from a multi-institutional study of intermediate-and high-risk localized PCa patients were obtained from Veracyte GRID (https://decipherbio.com/grid/).
13. **Tosoian 2020** (PMID: 32231245; DOI: 10.1038/s41391-020-0226-2) [33]: Transcriptomic profiles (Microarray) and clinicopathologic data of 405 radical prostatectomy and biopsy specimens from high-risk localized PCa patients were obtained from Veracyte GRID (https://decipherbio.com/grid/).
14. **CHAARTED correlatives 2021** (PMID: 34129855; DOI: 10.1016/j.annonc.2021.06.003) [34]: Transcriptomic profiles (Microarray) and clinicopathologic data of 160 biopsy specimens from a Phase 3 trial of metastatic hormone-sensitive PCa patients were obtained from the NCTN Data Archive (https://nctn-data-archive.nci.nih.gov), accession number NCT00309985-D14.
15. **Decipher GRID Biopsy** (PMID: 37060201; DOI: 10.1002/cncr.34790) [35]: Transcriptomic profiles from 82,470 prospectively collected prostate biopsy samples were retrieved from the Decipher GRID database (RRID: SCR_006552; https://decipherbio.com/grid; ClinicalTrials.gov identifier: NCT02609269; https://clinicaltrials.gov/ct2/show/NCT02609269). Patient data were de-identified from clinical use of the Decipher prostate genomic classifier in accordance with the Safe Harbor method described in the HIPAA Privacy Rule 45 CFR 164.514(b) and (c) (Veracyte, San Diego, CA). The samples, comprising a large cohort of localized PCa, were utilized to compare the distribution of Stemness Index values across National Comprehensive Cancer Network (NCCN) risk groups.

#### Clinicopathologic Breakdown and Risk Group Analysis

Detailed clinicopathologic review revealed that the majority of the cohort presents with low-grade localized PCa. The TNM staging breakdown includes:

- **T1 stage:** Approximately 50% of samples, indicating early-stage cancer localized within the prostate.
- **T2 stage:** Approximately 8%, representing cancer confined within the prostate but more extensive than T1.
- **T3 and T4 stages:** Less than 1%, suggesting advanced PCa with local spread.
- **N0:** Approximately 98%, indicating no regional lymph node involvement.
- **N1:** Approximately 2%, suggesting regional lymph node involvement.

The Decipher GRID database (RRID:SCR_006552) categorizes patients into NCCN risk groups based on TNM stage, PSA level, and Gleason score:

- **Very Low - Low Risk:** T1–T2a, N0, M0, PSA <10 ng/mL, and Gleason score ≤6; often managed with active surveillance or less aggressive treatment.
- **Intermediate Risk – Favorable:** Typically T2b–T2c, N0, M0, PSA 10–20 ng/mL, and Gleason score 7; managed with less aggressive treatment but closer monitoring.
- **Intermediate Risk – Unfavorable:** Similar clinical features to favorable but with higher disease burden or adverse factors requiring careful monitoring.
- **High Risk:** Usually involves T3a, N0, M0, PSA >20 ng/mL, or Gleason score 8–10; often managed aggressively with surgery, radiation, and/or hormonal therapy.
- **Very High Risk:** Typically includes T3b–T4, any N, M0, and high Gleason scores or PSA levels (PSA >20 ng/mL); patients are at significant risk for metastatic progression and are treated aggressively, often with multimodal therapies.

#### Observations on Disease Severity and Treatment Naïvety

Analysis of the cohort indicates that approximately 13% of cases fall into the High and Very High-Risk categories (8,376 + 2,399 = 10,775 samples), suggesting that the majority (approximately 87%) are below Intermediate Risk (≤GS7).

This distribution implies that most patients were treatment-naïve and likely ADT-free due to less aggressive disease status.

16 **Bolis 2021** (PMID: 34857732; DOI: 10.1038/s41467-021-26840-5) [36]: RNA-seq and clinicopathologic data of 1,223 clinical samples from an integrated cohort consisting of normal prostate specimens (n = 174), Pri-PCa (n = 714) and mCRPC (n = 335) was obtained from the Zenodo repository (Record ID: 5546618; https://zenodo.org/records/5546618). This resource of PCa Transcriptome Atlas (https://prostatecanceratlas.org) was built and integrated from the following studies/datasets: (1) Genotype-Tissue Expression Database [18] (GTEx; PMID: 23715323); (2) The Cancer Genome Atlas [22] (TCGA, TCGA-PRAD; PMID: 26544944); (3) Atlas of RNA-sequencing profiles of normal human tissues [37] (GSE120795; PMID: 31015567); (4) Integrative epigenetic taxonomy of PNPCa [38] (GSE120741; PMID: 30464211); (5) Prognostic markers in locally advanced lymph node-negative PCa (PRJNA477449); (6) The long noncoding RNA landscape of NEPC and its clinical implications [39] (PRJEB21092; PMID: 29757368); (7) Integrative clinical sequencing analysis of metastatic CRPC [10] (PRJNA283922; dbGaP: phs000915; PMID: 26000489); (8) CSER—exploring precision cancer medicine for sarcoma and rare cancers [40] (PRJNA223419; dbGaP: phs000673; PMID: 28783718); (9) Molecular basis of NEPC [41] (Beltran 2016; PRJNA282856; dbGaP: phs000909; PMID: 26855148); (10) Heterogeneity of androgen receptor splice variant-7 (AR-V7) protein expression and response to therapy in CRPC [42] (GSE118435; PMID: 30334814); (11) RNA-seq of human PCa cell lines and mCRPC [43–45] (GSE14750; PMIDs: 32460015; 33658518; 34244513 | GSE171729; PMID: 34244513); and (12) Molecular profiling stratifies diverse phenotypes of treatment-refractory metastatic CRPC [46] (PRJNA520923; GEO: GSE126078; PMID: 31361600).

17 **Taylor 2010** (PMID: 20579941; DOI: 10.1016/j.ccr.2010.05.026) [47]: Transcriptomic profiles (Microarray) and clinicopathologic (recurrence-free survival) data of 29 adjacent benign prostate tissue, 131 Pri-PCa, and 19 mPCa were obtained from the GEO database GSE21034 and the cBioPortal for Cancer Genomics PRAD MSKCC 2010 study (cBioPortal: MSKCC Prostate Adenocarcinoma cohort).

18 **SU2C 2015** (PMID: 26000489; DOI: 10.1016/j.cell.2015.05.001) [10]: Clinical and genomic data of 150 mCRPC samples were obtained from the cBioPortal for Cancer Genomics PRAD SU2C 2015 study (https://www.cbioportal.org/study/summary?id=prad_su2c_2015). RNA-seq data of 98 mCRPC samples was retrieved from Zhang *et al*. [23] (PMID: 32350277; DOI: 10.1038/s41467-020-15815-7).

19 **SU2C 2019** (PMID: 31061129; DOI: 10.1073/pnas.1902651116) [48]: This dataset includes clinicopathologic and genomic data for 444 mCRPC samples. RNA-seq data of 266 mCRPC samples (prepared by using poly(A) enrichment for RNA-seq library construction) were obtained from the cBioPortal for Cancer Genomics PRAD SU2C 2019 study (https://www.cbioportal.org/study/summary?id=prad_su2c_2019). In parallel with TCGA-PRAD, clinicopathologic features (age, Gleason scores, PSA levels), genomic alterations (mutation frequency, CNAs, SVs, TMB, fraction of genome altered), and patient outcome data (overall survival) were also retrieved from cBioPortal (RRID:SCR_014555). These datasets were used for comparative analysis between Stemness-high and Stemness-low groups, and to explore genomic drivers and clinical associations as described in the Methods.

20 **Labrecque 2019** (PMID: 31361600; DOI: 10.1172/JCI128212) [46]: RNA-seq data of 98 mCRPC samples and 39 PDX-LuCaP was obtained from the GEO database GSE126078.

21 **Alumkal 2020** (PMID: 32424106; DOI: 10.1073/pnas.1922207117) [13]: RNA-seq data of 25 mCRPC samples before treatment with enzalutamide was obtained from Supplementary dataset S01 of the indicated reference.

22 **Westbrook 2022** (PMID: 36109521; DOI: 10.1038/s41467-022-32701-6) [12]: RNA-seq data of 42 mCRPC samples consisting of 21 matched pre/post Enzalutamide treatment samples was retrieved from the indicated reference.

23 **Beltran 2016** (PMID: 26855148; DOI: 10.1038/nm.4045) [41]: RNA-seq data of mCRPC samples, including 34 castration-resistant adenocarcinoma (mCRPC-Adeno) and 15 mCRPC with neuroendocrine histologies (mCRPC-NEPC), was obtained from cBioPortal for the Cancer Genomics (https://www.cbioportal.org/study/summary?id=nepc_wcm_2016).

24 **Goodrich 2017** (PMID: 28059767; DOI: 10.1126/science.aah4199) [49]: RNA-seq data of murine prostate tumors from genetically engineered mouse models (GEMMs) of PCa of different genotypes, including 4 SKO (single knockout: PBCre4;*Pten^f/f^*), 13 DKO (double knockout PBCre4;*Pten^f/f^:Rb1^f/f^*), and 6 TKO (PBCre4;*Pten^f/f^:Rb1^f/f^:Trp53^f/f^*) was obtained from the GEO database GSE90891 (*f* = a floxed allele of the indicated genes).

25 **CCLE 2018** (PMID: 22460905; DOI: 10.1038/nature11003) [50]: RNA-seq data of 1,019 human cell lines were obtained from the Xena Functional Genomics Portal [25] (https://xenabrowser.net, version: 2018-05-30). Among them, eight PCa cell lines — DU145 (RRID:CVCL_0105), LNCaP clone FGC (RRID:CVCL_1379), MDA PCa 2b (RRID:CVCL_4745), NCI-H660 (RRID:CVCL_0459), PC-3 (RRID:CVCL_0035), VCaP (RRID:CVCL_2235), 22Rv1 (RRID:CVCL_1045), and PRECLH (PrEC LH; RRID:CVCL_V626) — were included for analysis.

26 **XG-LNCaP** and **XG-LAPC9** (PMID: 30190514; DOI: 10.1038/s41467-018-06067-7) [51]: RNA-seq data of 12 LNCaP xenografts (XG-LNCaP) and 10 LAPC9 xenografts (XG-LAPC9) was obtained from the GEO database GSE88752.

27 **Whitlock 2020** (LNCaP MYC knockdown) (PMID: 32681068; DOI: 10.1038/s41388-020-01389-7) [52]: RNA-seq data for the LNCaP MYC-knockdown system were obtained from the GEO database (GSE135942). This dataset included non-targeting control samples (n=3) and MYC-knockdown samples (n=6) generated using two shMYC constructs (shMYC-512 and shMYC-637) targeting the MYC open reading frame (ORF), each performed in triplicate. The shMYC-627 construct, which targets the 3′UTR, was excluded from analysis.

28 **Holmes 2022** (22Rv1 MYC inhibition) (PMID: 35476451; DOI: 10.1126/sciadv.abh3635) [53]: RNA-seq data for MYC inhibition in the AR-positive CRPC cell line 22Rv1 were obtained from the GEO database GSE178869. The dataset included 22Rv1 cells treated with the small-molecule MYC inhibitor MYCi975 at 10 μM for 48 h or with 0.2% DMSO as the vehicle control (n = 8). These data were used to evaluate changes in MYC activity and Stemness following direct pharmacological inhibition of MYC.

29 **Han 2019** (PC3 MYC inhibition) (PMID: 31679823; DOI: 10.1016/j.ccell.2019.10.001) [54]: RNA-seq data for pharmacological MYC inhibition in the AR-negative cell line PC3 were obtained from the GEO database. GSE135800 included PC3 cells treated with MYCi975 at 8 μM for 24 h or with DMSO as the vehicle control (n = 6), whereas GSE135876 included PC3 cells treated with MYCi361 at 6 μM for 24 h or with DMSO as the vehicle control (n = 6). These independent MYC-inhibitor datasets were analyzed to assess the effects of direct MYC inhibition on MYC activity and Stemness in an AR- negative ARPI-refractory PCa cell model.

30 **Coleman 2019** (PCa BET inhibition) (PMID: 30846826; DOI: 10.1038/s41598-019-40518-5) [55]: RNA-seq data from a panel of 11 human PCa cell lines treated with the BET bromodomain inhibitor JQ1 were obtained from the GEO database GSE98069. The dataset included paired vehicle-and JQ1-treated samples from LNCaP, abl, C4-2B, LN95, 22Rv1, V16D, MR42D, MR49F, M12, PC3, and NCI-H660 cells (total n = 22). Cells were treated with 500 nM JQ1 or 0.05% DMSO vehicle for 24 h. These cell lines collectively represented androgen-dependent PCa, AR-positive CRPC, enzalutamide-resistant CRPC, AR-negative CRPC, and NEPC. These data were used to determine whether BET bromodomain inhibition, which suppresses MYC-associated transcriptional activity while also affecting additional BET-dependent programs, was accompanied by changes in MYC activity and Stemness across diverse PCa states.

31 **Watanabe 2023** (PCa spatial transcriptomics) (PMID: 37240308; DOI: 10.3390/ijms24108955) [56]: Spatial transcriptomic data from a treatment-naïve Pri-PCa specimen containing spatially coexisting de novo NEPC (pT4N1M0) and AR pathway-positive prostate adenocarcinoma (ARPC; Gleason score 4 + 4, pT2aN0M0) were obtained from the GEO database GSE230282 (patient n = 1; specimen n = 1). Formalin-fixed, paraffin-embedded tissue was profiled using the 10x Genomics Visium CytAssist Spatial Gene Expression platform, generating 4,397 tissue-covered spatial spots. These data were used to compare the spatial distributions of Stemness and PCa-Stem enrichment between the pathologically annotated NEPC and ARPC regions.

32 **Zhao 2024** (Integrated PCa scRNA-seq meta-atlas) (PMID: 39418984; DOI: 10.1016/j.ebiom.2024.105398) [57]: Processed scRNA-seq data from an integrated human PCa scRNA-seq meta-atlas were obtained from the Zenodo repository (Record ID: 12605534; https://zenodo.org/records/12605534). The meta-atlas integrated 10 previously published human PCa scRNA-seq datasets comprising 93 samples and 222,529 high-quality cells, including: (1) GSE117403 (PMID: 30566875; n = 6); (2) GSE150692 (PMID: 32915138; n = 3); (3) GSE137829 (PMID: 33328604; n = 6); (4) GSE157703 (PMID: 33032611; n = 2); (5) SCP1244/phs001988.v1.p1 (PMID: 33664492; n = 15); (6) GSE143791 (PMID: 34719426; n = 16); (7) EGAS00001005787 (PMID: 34936871; n = 20); and (8–10) three batches from Chen et al. (PMID: 33420488), including GSE141445 (n = 13) and two additional datasets from the PCa Cell Atlas (www.pradcellatlas.com; n = 3 and 9). Collectively, these datasets span normal prostate, adjacent benign prostate tissue, Pri-PCa, and mCRPC. Following epithelial-cell re-clustering, 89,055 epithelial cells classified as basal, hillock, club, luminal, lineage plasticity-related cells (LPCs), and NE cell populations were retained for the present analyses. These epithelial cells were used to quantify global Stemness, PCa-Stem activity, and expression of the individual PCa-Stem signature genes across prostate epithelial cell states.

33 **Cheng 2022** (FACS-enriched prostate epithelial scRNA-seq) (PMID: 35058087; DOI: 10.1016/j.eururo.2021.12.039) [58]: scRNA-seq data from FACS-isolated human prostate epithelial cells were obtained from the NCBI Sequence Read Archive (PRJNA699369). The original study included specimens from six patients, comprising three treatment-naïve Pri-PCa (including matched adjacent benign tissues for two patients), two locally recurrent CRPC cases (including one small-cell neuroendocrine carcinoma; SCNC), and one mCRPC case. Epithelial populations were enriched by fluorescence-activated cell sorting using Trop2, CD45, CXCR2, and CD49f markers. The remapped expression matrix used in the present study contained 26,142 cells from 11 FACS-derived epithelial sample-level datasets. which were used to evaluate PCa-Stem activity and expression of the 12 individual PCa-Stem signature genes across the annotated cell populations.

#### Sex as a biological variable

All clinical datasets analyzed in this study consisted exclusively of male patients, consistent with the male-specific nature of PCa. Preclinical models, including xenografts, patient-derived xenografts, and GEMMs were also male-derived. All PCa cell lines analyzed in the study were of male origin, with the exception of Figure S5A, where a broader cohort of 1,019 cell lines from the CCLE, comprising both male-and female-derived samples across multiple cancer types, was analyzed. Analyses of CCLE data in Figure S5A were performed without stratification by sex.

### II. Bulk transcriptomic data normalization and transcriptome-based signature quantification

To ensure consistency across diverse transcriptomic datasets, all data acquisition and normalization, and signature quantification (scoring) were performed using methods tailored to data type and analytical scope. All analyses were conducted within each dataset, thereby avoiding inter-study batch effects. For comparisons across pathological stages (e.g., normal/benign → Pri-PCa → mCRPC), we used only harmonized, pre-integrated transcriptomic datasets, including the PCa Transcriptome Atlas (Bolis 2021) [36] and Taylor 2010 [47], which provide variance-stabilized, library size–adjusted expression matrices normalized across disease stages.

In this project, we adopted, derived, and utilized multiple transcriptomic gene signatures representing PCa differentiation, stemness, oncogenic signaling, and tumor-suppressor loss. Signatures analyzed included: (1) the Stemness Index, (2) Canonical AR activity (c_AR-A; 10 genes), (3) MYC signaling activity (MYC-sig; 58 genes), (4) the PCa-Stem Signature (12 genes), (5) castration-reprogrammed AR activity (cr_AR-A; 63 genes), (6) cr_AR-Growth_Wang (8 genes), (7) *RB1*-loss signature (158 genes), and (8) *PTEN*-loss signature (45 genes).

#### Transcriptome-based quantitative measurement of oncogenic stemness (the Stemness Index)

We implemented mRNA-based **Stemness Index** (mRNAsi) [6] to determine the degree of oncogenic dedifferentiation and to quantify the aggressiveness in PCa. Briefly, we computed Spearman correlations between the stemness model’s weight vector and the transcriptome expression profile of the queried samples. Spearman correlation calculations were performed using the ‘stats’ package in R (version 4.3.3; RRID:SCR_001905). The resulting correlation coefficients were then linearly transformed to scale the mRNAsi scores between 0 and 1 using the following formula: Stemness Index = (Spearman correlation + 0.2619) /0.4151. Spearman correlation was adopted for calculating the Stemness Index because of its robustness to non-normal expression distributions and its reduced susceptibility to cross-dataset batch effects inherent in large-scale integrative analyses.

#### Canonical AR activity and MYC activity

Canonical AR activity (**c_AR-A**) was determined by RNA-seq based measurements of AR-regulated transcripts under intact androgen/AR signaling conditions. Similarly, the MYC signaling (**MYC-sig**) activity was determined by MYC-regulated transcripts. We employed Z-score normalization [59] to quantify c_AR-A and MYC-sig. Briefly, the numeric AR or MYC activity score was calculated based on a linear combination of expression values (Z-scores) of experimentally validated AR target genes (10 genes) [60] or MYC target genes (58 genes; GSEA Molecular Signature Database 2022.1 version, MSigDB) [61]. The target gene lists were detailed in Table S2. The expression value for each AR or MYC target gene was converted to Z-score by Z = (x − μ)/σ, where x is the expression value of the target gene in a specific sample, μ is the mean and σ is the standard deviation across all samples of a gene. Finally, the combined Z-scores were summed across all genes to represent the expression score of canonical AR or MYC target genes. All Z-score analyses were confined to individual datasets to avoid cross-study normalization artifacts.

For cross-stage analyses using harmonized datasets (e.g., the PCa Transcriptome Atlas; ref. [36]), c_AR-A and MYC-sig were additionally quantified using single-sample gene set enrichment analysis (ssGSEA) to enable direct comparison with other gene set–based signatures.

#### PCa-Stem Signature (the 12-gene PCa-specific stemness signature)

A PCa–specific stemness signature was constructed by identifying genes consistently enriched in high-stemness tumors across distinct disease states. Differential expression analysis compared the top 33% (Stemness-high) versus bottom 33% (Stemness-low) samples in treatment-naïve Pri-PCa (TCGA-PRAD) and mCRPC (SU2C 2019) cohorts. Genes upregulated in both cohorts (fold change ≥ 2, FDR < 0.05; Table S3) were intersected to generate the 12-gene **PCa-Stem signature** (Table S2). Per-sample enrichment scores were quantified by ssGSEA.

#### Castration reprogrammed AR activity (cr_AR-A) signatures

The 63-gene **cr_AR-A** (Table S2) signature was derived by integrating differential chromatin analyses of AR and H3K27ac ChIP-seq performed in CRPC versus Pri-PCa samples (GSE130408) [16] with differential transcriptional analysis comparing CRPC and Pri-PCa samples (GSE21032) [47], thereby identifying genes exhibiting concordant CRPC-enriched AR binding, enhancer activation, and transcriptional up-regulation.

The **cr_AR-Growth_Wang** (8 genes; Table S2) signature was adopted from Wang *et al*. [14] and consists of AR-bound, enhancer-validated growth regulators identified in androgen-independent LNCaP-abl cells, representing a proliferation-oriented, castration-reprogrammed AR transcriptional program. Both AR-reprogrammed signatures were quantified by ssGSEA.

#### Tumor-suppressor-loss signatures

Transcriptional signatures reflecting functional loss of key tumor suppressors were analyzed using previously validated gene sets. The ***RB1*-loss signature** (158 genes; Table S2), derived from Ertel *et al.* [62], captures E2F-driven transcriptional deregulation associated with RB1 pathway inactivation. The ***PTEN*-loss signature** (45 genes; Table S2), derived from Liu *et al.* [63], represents transcriptional consequences of *PTEN* loss. Both signatures were quantified using ssGSEA. *<u>ssGSEA implementation</u>:* ssGSEA was performed using the Gene Set Variation Analysis (GSVA) [64] package (version 1.50.5; RRID:SCR_021058) in R (version 4.3.3; RRID:SCR_001905) to quantify per-sample enrichment scores. All ssGSEA analyses were conducted within individual datasets, using rank-based enrichment to minimize sensitivity to expression scale differences. For analyses spanning disease stages, ssGSEA was applied to the harmonized transcriptomic datasets to minimize batch effects. All gene sets are provided in Table S2.

### III. Network and pathway enrichment analyses

Protein–protein interaction (PPI) networks of the 12-gene PCa-Stem signature were generated using the STRING database (version 12.0; RRID:SCR_005223) [65]. An extended stemness-associated PPI network was additionally generated by integrating the PCa-Stem signature genes with representative stemness-and lineage-plasticity-associated regulators.

Functional enrichment analyses of the PCa-Stem signature were performed using Metascape (version 3.5; RRID:SCR_016620) [66] and ToppGene Suite [67], following established principles for pathway enrichment analysis [68]. Enrichment was assessed for Gene Ontology biological process, molecular function, and cellular component terms, together with curated pathway collections.

#### Gene set enrichment analysis (GSEA)

GSEA was carried out using the GSEAPreranked module of GSEA software (version 4.2.3; RRID:SCR_003199). For each comparison, the entire list of detected genes was ranked by sign(log2 fold change) × −log10(FDR), where FDR represents the multiple-testing-adjusted significance from the corresponding differential-expression analysis. The resulting ranked gene list was used for pre-ranked GSEA. Hallmark and curated gene sets (C2) from the Molecular Signatures Database (MSigDB, version v2023.1.Hs), together with additional gene signatures curated from the literature, were interrogated; the gene sets used in the present study and their original sources are summarized in Table S5. Analyses were performed following the standard GSEAPreranked procedure described in the GSEA User Guide, using the weighted enrichment statistic, 1,000 gene-set permutations, and gene-set sizes of 15–500 genes. Enrichment was assessed by normalized enrichment score (NES), nominal P value, and GSEA FDR q value, with FDR q < 0.05 considered statistically significant [69].

#### Ingenuity Pathway Analysis (IPA)

QIAGEN Ingenuity Pathway Analysis (IPA; Spring 2026 release; RRID:SCR_008653) was used to identify predicted upstream regulators and literature-supported molecular relationships associated with the PCa-Stem program. The 12-gene PCa-Stem signature was uploaded into IPA and analyzed using Core Analysis according to the manufacturer’s recommended settings. Upstream-regulator associations were evaluated using right-tailed Fisher’s exact test/overlap P values [70], and regulatory networks were constructed based on direct and indirect relationships curated in the IPA Knowledge Base. For the upstream-regulator network, IPA-derived regulator–target relationships were exported as an edge list and imported into Cytoscape Web (version 1.0.8; RRID:SCR_003032) for visualization and manually organized into three functional layers: upstream transcription-factor regulators, plasticity-associated signal relayers, and downstream stemness executors. Separately, a stemness-anchored extended regulatory network was generated within IPA by integrating the 12 PCa-Stem signature genes with established stemness-associated regulators and signaling molecules.

### IV. Spatial transcriptomic analysis

Publicly available Visium spatial transcriptomic data (GSE230282) were obtained from the GEO database. The dataset had been previously annotated by pathologists as containing coexisting *de novo* neuroendocrine PCa (NEPC) and AR pathway-positive prostate adenocarcinoma (ARPC) within the same tissue section [56]. Spatial transcriptomic data were processed and analyzed following the analytical framework described by Zhao *et al.* (2024) [57]. The published processed dataset and pathological annotations were retained, and additional PCa-Stem-and stemness-related analyses were performed as described below. PCa-Stem signature scores for individual spatial spots were calculated using the same AUCell-based approach applied to the scRNA-seq datasets. Global Stemness Index (mRNAsi) was calculated using the transcriptome-based stemness algorithm [6] described earlier. Spearman correlation analysis was performed to evaluate the concordance between PCa-Stem and mRNAsi across spatial transcriptomic spots. Finally, both stemness metrics were mapped onto the tissue section and compared between the pathologist-defined NEPC and ARPC regions to evaluate their spatial distribution and heterogeneity.

### V. Single-cell RNA-seq analysis

To investigate cancer stemness during PCa progression at single-cell resolution, two independent human PCa scRNA-seq resources were analyzed. Discovery analyses were performed using integrated PCa epithelial scRNA-seq meta-atlas established by Zhao *et al.* (2024)[57], which comprises 89,055 epithelial cells from 10 published PCa scRNA-seq datasets spanning normal/benign prostate, Pri-PCa, and mCRPC. The published Seurat object and epithelial cell annotations were retained for downstream analyses. New AUCell-based signature scoring was performed to quantify PCa-Stem signature across the annotated epithelial cell populations without modifying the original clustering or cell-type annotations [57]. Expression of the individual PCa-Stem signature genes was also evaluated across epithelial cell states.

Independent validation was performed using Cheng *et al.* (2022) PCa scRNA-seq dataset, comprising 26,142 cells from 11 FACS-enriched sample-level datasets [58]. Although epithelial cells constituted the predominant cell population, small numbers of leukocytes, myeloid cells, endothelial cells, and fibroblasts remained in the dataset, consistent with the original publication [58]. Cell-type annotations followed the classification framework established by Zhao *et al.* (2024) [57]. The remapped expression matrix was analyzed using a consistent downstream signature-scoring framework to evaluate PCa-Stem activity and expression of the individual PCa-Stem signature genes across the annotated cell populations. *<u>PCa-Stem signature scoring using AUCell</u>:* Stemness activity at single-cell resolution was quantified using the AUCell algorithm implemented in the AUCell R package. The previously established PCa-Stem signature, consisting of 12 stemness-associated genes (*AURKB, BIRC5, CENPA, DEPDC1B, DLGAP5, HJURP, HMMR, KLK12, MELK, NEK2, PBK, and UBE2C*), was used as the input gene set. AUCell calculates the Area Under the Curve (AUC) representing the enrichment of a gene signature within the highest-ranked expressed genes of each individual cell. AUCell scores were calculated for individual epithelial cells based on rank-based gene signature enrichment, enabling robust evaluation of PCa-Stem activity independent of sequencing depth. The resulting AUCell scores were visualized on Uniform Manifold Approximation and Projection (UMAP) embeddings and compared among epithelial cell subtypes and disease stages to characterize the distribution of stemness during PCa progression.

#### Identification of PCa-Stem⁺ epithelial cells using AUCell

To further classify individual epithelial cells according to PCa-Stem status, an AUC threshold was determined using the automatic threshold-selection procedure implemented in the AUCell package. Cells with AUC scores above the selected threshold were classified as PCa-Stem⁺ epithelial cells whereas those with scores at or below the threshold were designated PCa-Stem⁻. The proportion of PCa-Stem⁺ epithelial cells was subsequently calculated for each sample and compared across different clinical disease states to evaluate dynamic changes in Stemness during PCa progression.

#### Independent validation in the Cheng et al. 2022 dataset

The robustness of the PCa-Stem signature was independently evaluated using the FACS-enriched prostate epithelial scRNA-seq dataset generated by Cheng *et al.* [58]. AUCell-based PCa-Stem signature scoring was applied to evaluate the distribution of PCa-Stem activity across the annotated cell populations, and expression of the individual PCa-Stem signature genes was examined to assess the reproducibility of PCa-Stem-associated cellular distributions in this independent patient cohort.

#### Pseudobulk analysis and comparison of PCa-Stem with mRNAsi

To facilitate a direct comparison between the PCa-Stem signature and the global Stemness Index (mRNAsi) at the sample level, a pseudobulk strategy was employed. Specifically, epithelial cell expression profiles from individual patient samples were aggregated to generate pseudobulk expression matrices, followed by normalization using Seurat’s pseudobulk workflow. This pseudobulk strategy was used to reduce the impact of sparsity and dropout inherent to scRNA-seq and to provide transcriptome-level expression profiles suitable for calculation of mRNAsi. mRNAsi scores were computed for each pseudobulk sample using the previously described one-class logistic regression (OCLR)-based method [6]. In parallel, PCa-Stem activity was quantified at the pseudobulk level using ssGSEA [64] based on the previously established 12-gene PCa-Stem signature. The resulting ssGSEA enrichment scores were used as sample-level PCa-Stem scores. Spearman correlation analysis was performed to assess the concordance between PCa-Stem ssGSEA scores and mRNAsi across pseudobulk samples. The two stemness measures were further compared across clinical disease stages to evaluate their respective associations with PCa progression.

### VI. Integrative analysis of genomic alterations in Pri-PCa (TCGA) and mCRPC (SU2C)

To explore molecular and clinical features associated with the PCa stemness, we performed integrative analysis of genomic, transcriptomic, clinicopathologic, and outcome data using cBioPortal for Cancer Genomics (RRID:SCR_014555), which provides interactive access to multidimensional cancer genomics datasets and integrated visualization tools [71]. Specifically, we analyzed data from Pri-PCa (TCGA-PRAD 2018, Pan-Cancer Atlas) and mCRPC (SU2C 2019) cohorts. Clinicopathologic variables (e.g., age, Gleason scores, PSA levels, tumor stage), genomic alterations (e.g., mutation frequency, TMB, fraction of genome altered, CNAs, and SVs), and patient outcome data (e.g., overall survival, progression-free survival) were retrieved from cBioPortal (RRID:SCR_014555) as described in the “PCa clinical datasets and data collection” section. Transcriptomic data used for the Stemness Score calculation were obtained as described in the same section. For comparative analysis, mCRPC samples were stratified into top and bottom 33% by the Stemness Scores. In TCGA, only treatment-naïve Pri-PCa samples were used for a similar 33% stratification, excluding 70 patients who received adjuvant (post-surgery) hormone therapy to reflect spontaneous disease progression. Differential expression analysis was conducted using DESeq2 [72] (version 1.42.1; RRID:SCR_015687), and gene-level and genome-wide alteration frequencies were assessed across stratified groups and disease stages (e.g., Pri-PCa vs. mCRPC) using OncoPrint visualizations, bar plots, and correlation analysis. Associations with clinical outcomes were evaluated as described in the “Statistical analysis” section.

### VII. Functional characterization of representative PCa-Stem signature genes in castration-resistant LAPC4-AI and androgen-sensitive LNCaP cells

#### Cell culture

The androgen-independent (AI) LAPC4 (LAPC4-AI) cells were derived as described [73] and cultured at 37°C with 5% CO₂ in IMDM (Thermo Fisher, Cat# 21056023) supplemented with charcoal-stripped serum and 1× penicillin/streptomycin. LNCaP prostate cancer cells were cultured in RPMI-1640 medium (Gibco, Cat# 11875093) supplemented with 10% fetal bovine serum (FBS) and 1× penicillin/streptomycin. Both cell lines were maintained under standard tissue culture conditions at 37°C in a humidified incubator with 5% CO₂. Cells were routinely passaged to maintain exponential growth and confirmed to be mycoplasma-free prior to experimentation.

#### siRNA transfection

Cells at ∼70% confluence were transfected with 1 µM Accell SMARTpool siRNAs targeting *HMMR*, *AURKB*, or *PBK*, or a non-targeting control (NTC) siRNA (Horizon Discovery). The following reagents were used: Accell Human *HMMR* siRNA SMARTpool (E-010409-00-0005); Accell Human *PBK* siRNA SMARTpool (E-005390-00-0005); Accell Human *AURKB* siRNA SMARTpool (E-003326-00-0005). siRNA complexes were prepared in serum-free medium and added to cells at 1 µM final concentration per manufacturer’s protocol. Media were replaced with complete medium after incubation for 24 h. Knockdown efficiency and functional assays were performed at the indicated time points.

#### RNA isolation and qRT-PCR

Total RNA was isolated 48 h post-transfection using the PicoPure RNA Isolation Kit (Applied Biosystems, Cat# KIT0204). RNA (0.5 μg) was reverse-transcribed using SuperScript™ VILO™ Master Mix (Invitrogen, Cat# 11755050). Quantitative PCR was carried out utilizing TaqMan™ Fast Advanced Master Mix (Applied Biosystems, Cat# 4444557) and pre-designed, gene-specific TaqMan probes for AURKB (Assay ID: Hs00945858_g1), PBK (Assay ID: Hs00902990_m1), and HMMR (Assay ID: Hs00234864_m1). β-Actin (ACTB; Assay ID: Hs01060665) was utilized as an endogenous control. Amplification and fluorescence detection were performed on a QuantStudio™ 6 Flex Real-Time PCR System (Applied Biosystems). Relative gene expression was calculated using the comparative CT (2^(-ΔΔC_T)) method, normalized to ACTB, and plotted relative to the non-targeting control (NTC). Data were presented as mean ± SD from technical triplicates.

#### Trans-well Matrigel invasion assays

At 48 h post-transfection, cells were harvested and equal numbers were seeded into Matrigel-coated trans-well inserts (Corning, Cat# 354480) in serum-free medium. Medium containing serum was placed in the lower chamber as a chemoattractant. After 24 h, non-invading cells were removed, and invaded cells on the underside of the membrane were fixed and stained with DAPI. Images were acquired using a Keyence microscope. Invaded cells were quantified by counting nuclei in multiple non-overlapping fields per insert. Data were presented as mean ± SD from technical replicates.

#### Colony formation assays

At 24 h post-transfection, cells were replated at clonal density (500 cells per well in 6-well plates) and cultured for 14 days. Colonies were fixed and stained with crystal violet with representative images shown. Where indicated, colonies were quantified using a consistent size threshold across conditions.

#### 3D spheroid formation and real-time cytotoxicity assays

At 24 h after siRNA transfection, LAPC4-AI cells were seeded for 3D culture using live-cell compatible Corning 96-well clear-bottom plates (Corning, Cat# 3595) and organoid medium mixed with 5% Matrigel (Corning, Cat# 356255) [73]. Real-time cytotoxicity assays were initiated 24 h later and monitored for 72 h using a cell-impermeant dead-cell dye (Thermo Fisher, Cat# L7013). Fluorescence was recorded at defined intervals using an IncuCyte live-cell imaging system (Sartorius) to assess cell death kinetics. Fluorescent signals were background-corrected using cell-free wells. Data were presented as mean ± SD from technical triplicates.

#### Prostasphere-Forming Efficiency and Stemness Assay

To evaluate intrinsic stemness following gene knockdown, a prostasphere-forming efficiency assay was performed using AggreWell™400 24-well plates (Catalog #34411). The engineered geometry of these plates, containing approximately 1,200 microwells per well, prevents random clumping. Plates were pre-treated by adding 500 µL of Anti-Adherence Rinsing Solution (Catalog #07010) per well and centrifuging at 1300 x g for 5 minutes in a swinging bucket rotor to remove air bubbles from the microwells. The wells were then aspirated, rinsed with 2 mL of warm medium, and pre-filled with 1 mL of warm, serum-free MammoCult™ Medium (Catalog #05620).

Following filtration through a 37 µm Reversible Strainer (Catalog #27215), strict single-cell suspensions of siRNA-transfected cells were seeded at exactly 12,000 cells per well to achieve a calibrated density of precisely 10 cells per microwell. Additional MammoCult™ Medium was added to achieve a final volume of 2 mL per well. Plates were then centrifuged at 100 x g for 3 minutes to capture the cells in the microwells and cultured at 37°C with 5% CO₂ and 95% humidity.

To track self-renewal kinetics longitudinally, plates were immediately transferred to the IncuCyte System (37°C, 5% CO₂). Whole-well phase-contrast images were captured every 24 hours using 10x objective. Sphere growth dynamics (sphere Area) were calculated using Image J.

#### Double Immunofluorescence (IF) Staining

Cells were fixed with 4% paraformaldehyde for 10–15 minutes and permeabilized using 0.25% Triton X-100. To prevent non-specific binding, samples were blocked with 3% BSA. To visualize mitotic chromosomes and spindle networks, cells were incubated overnight at 4°C with mouse anti-Phospho-Histone H3 (Ser10) (CST #9706, 1:200) and rabbit anti-alpha/beta-Tubulin (CST #2148, 1:100). Following washing, samples were incubated with Alexa Fluor 488-conjugated anti-mouse and Alexa Fluor 594-conjugated anti-rabbit secondary antibodies (1 hour, dark). Coverslips were mounted using ProLong™ Gold Antifade Mountant with DAPI (Invitrogen #P36931) for simultaneous nuclear counterstaining and photobleaching prevention.

#### Western Blotting

Cells were lysed in RIPA buffer (Thermo Scientific, 89900) with Halt™ Protease/Phosphatase Inhibitors (Thermo Scientific, 78440), sonicated on ice (10–15s), and centrifuged (14,000 × g, 15 min, 4°C). Protein concentrations were quantified via BCA assay (Thermo Scientific, 23225). Lysates (15–30 µg) were denatured in 4X NuPAGE™ LDS Sample Buffer (Invitrogen, NP0007) with Reducing Agent (Invitrogen, NP0004) at 90°C for 10 min. Proteins were resolved on NuPAGE™ 4– 12% Bis-Tris gels (Invitrogen, NP0321BOX) in MES SDS buffer (Invitrogen, NP0002) and transferred to 0.45 µm PVDF membranes using NuPAGE™ Transfer Buffer (Invitrogen, NP0006). Membranes were blocked in 5% BSA/TBST (1 h, RT) and incubated overnight at 4°C with primary antibodies against PBK (Novus, NBP1-84342), RHAMM/CD168 (Novus, NBP1-76538), Aurora B (Abcam, ab2254), γ-H2AX (Millipore, 05-636), and β-Actin (Santa Cruz, sc-47778 HRP). Following TBST washes, membranes were incubated with HRP-conjugated secondary antibodies (1 h, RT). Protein bands were visualized using Bright Star developing solution (SKU: MMZR-XR9) and imaged on a Bio-Rad ChemiDoc Go Imaging System.

### VIII. Statistical analyses

Statistical analyses were performed using GraphPad Prism software (version 10.4.1 (532); RRID:SCR_002798) and R (version 4.3.3; RRID:SCR_001095). For comparisons between two groups, independent sample or paired *t*-tests were used to compare group means, assuming normality and equal variance. For non-parametric comparisons between two groups, the Wilcoxon rank-sum test or Wilcoxon matched-pairs signed-rank test was used as appropriate. For multi-group comparisons, Tukey’s range test was employed following one-way ANOVA (adjusted p-values listed in Table S4). For non-parametric comparisons involving more than two groups, the Kruskal–Wallis test followed by appropriate post hoc multiple-comparison correction was used. The Wilcoxon rank-sum test was applied for comparisons of altered genome fraction, TMB, or age between two groups. For differential expression analyses and comparisons involving multiple hypotheses (e.g., RNA-seq-based gene comparisons), p-values were further adjusted using the false discovery rate (FDR) method to control for multiple testing.

Pearson’s correlation coefficient was used to evaluate linear associations between two continuous variables, assuming normally distributed paired observations. Linear regression modeling was applied to examine the relationship between two continuous variables, with the slope of the best-fit regression line reported where applicable. Spearman’s rank correlation was employed to assess monotonic relationships between continuous or ordinal variables.

The Jonckheere-Terpstra (J-T) trend test was used to assess monotonic association between an ordinal variable and a continuous variable. For analyses involving proportions across ordered disease categories, the Cochran–Armitage trend test was used to test for a linear trend in alteration proportions across GS6, GS9–10, Pri-PCa/ADT, and mCRPC. Chi-square test or Fisher’s exact test was performed to evaluate the associations between categorical variables.

For the analysis of patient progression-free survival (PFS), recurrence-free survival (RFS) and overall survival (OS), the standard Kaplan-Meier method was used for the estimation of survival fractions and log-rank tests were used for group comparisons. Cox proportional hazards models were used for multivariable analysis and for assessing the prognostic significance of gene signatures (the Stemness Index, PCa-Stem signature, etc). For the mRNA-based gene signatures (the Stemness Index, PCa-Stem signature, c_AR-A), patients were stratified into high and low signature groups based on median split in cohorts including Spratt 2017 (Fig. 4A), Tosoian 2020 (Fig. 4B), SU2C 2019 (Fig. S6A), and Taylor 2010 (Fig. S6D-E). For some analyses, including CHAARTED 2021 (Fig. 4C) and SU2C 2019 (Fig. S6B), patients were alternatively stratified by comparing the top 25% versus the bottom 75% of signature scores to assess whether patients with very high Stemness levels exhibited a higher risk of disease progression.

**Figure 1.**
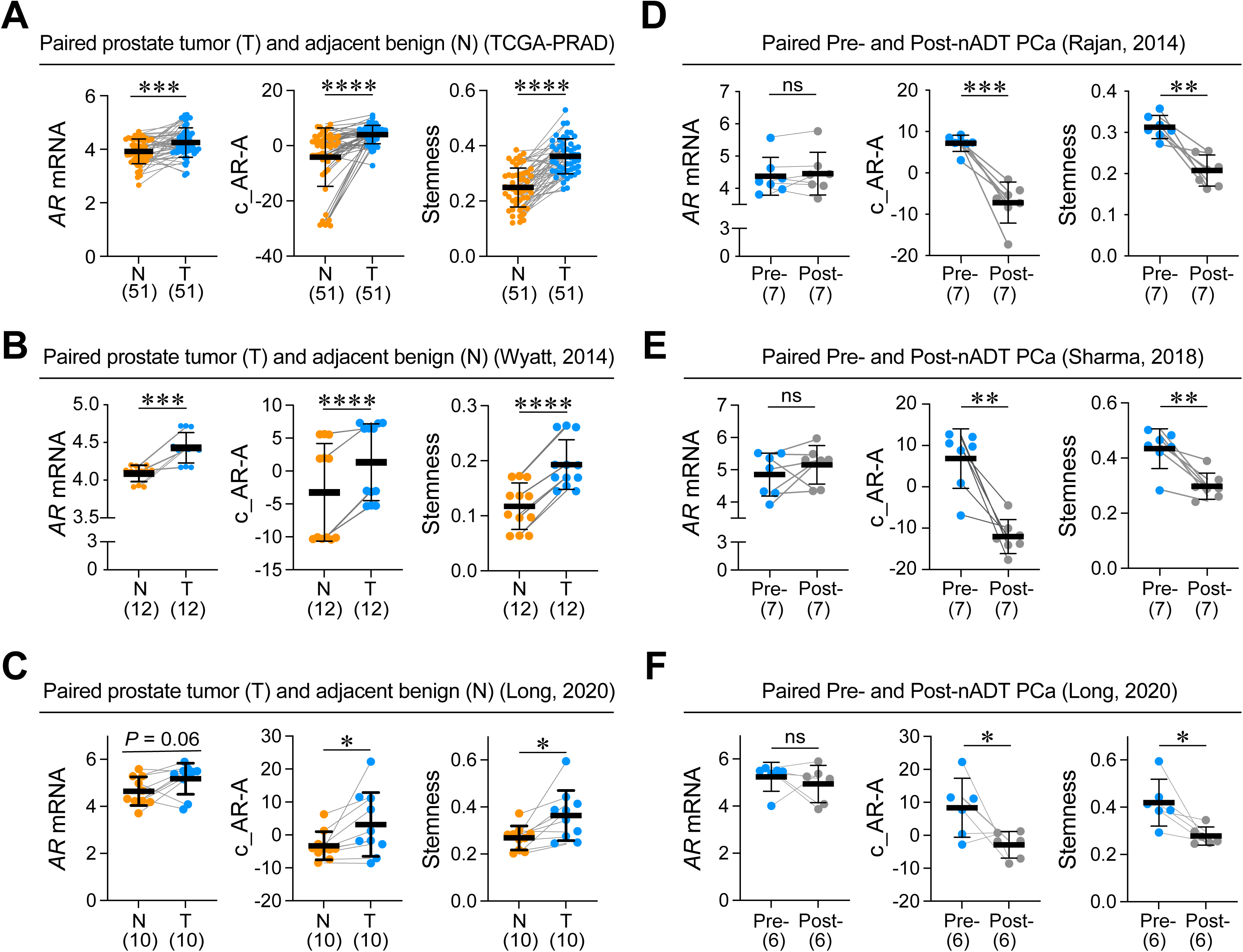
Pri-PCa shows concordant increases of c_AR-A and Stemness and both are decreased by nADT. **A-C,** PCa has increased c_AR-A as well as the Stemness compared to matched benign tissues. Pairwise comparisons of *AR* mRNA levels, canonical c_AR-A, and Stemness scores showing significant increases in primary tumors (T) as compared to the matched adjacent benign prostate tissues (N) from the same individual in TCGA-PRAD cohort (n=51; **A**), and Wyatt 2014 cohort (n=12; **B**). In the Long 2020 cohort (n=10; **C**), increases in c_AR-A and Stemness reached statistical significance whereas the increase in *AR* mRNA levels approached statistical significance (*P* = 0.06). **D-F,** nADT decreases both c_AR-A and Stemness. Pairwise comparison of *AR* mRNA levels, c_AR-A, and Stemness showing significant decrease of both c_AR-A and Stemness after nADT (Post-nADT) as compared to the matched samples before nADT (Pre-nADT) from the same individual (**D**, Rajan 2014 cohort; **E**, Sharma 2018 cohort; **F**, Long 2020 cohort). In all plots, sample sizes are indicated (see also Table S1 for cohort and RNA-seq data access information). Within the plots, results are shown as the mean ± SD and pairs are linked with a grey line. Significance was calculated by two-tailed paired Student’s *t*-test (ns, not significant; *, *P* < 0.05; **, *P* < 0.01; ***, *P* < 0.001; ****, *P* < 0.0001).

**Figure 2.**
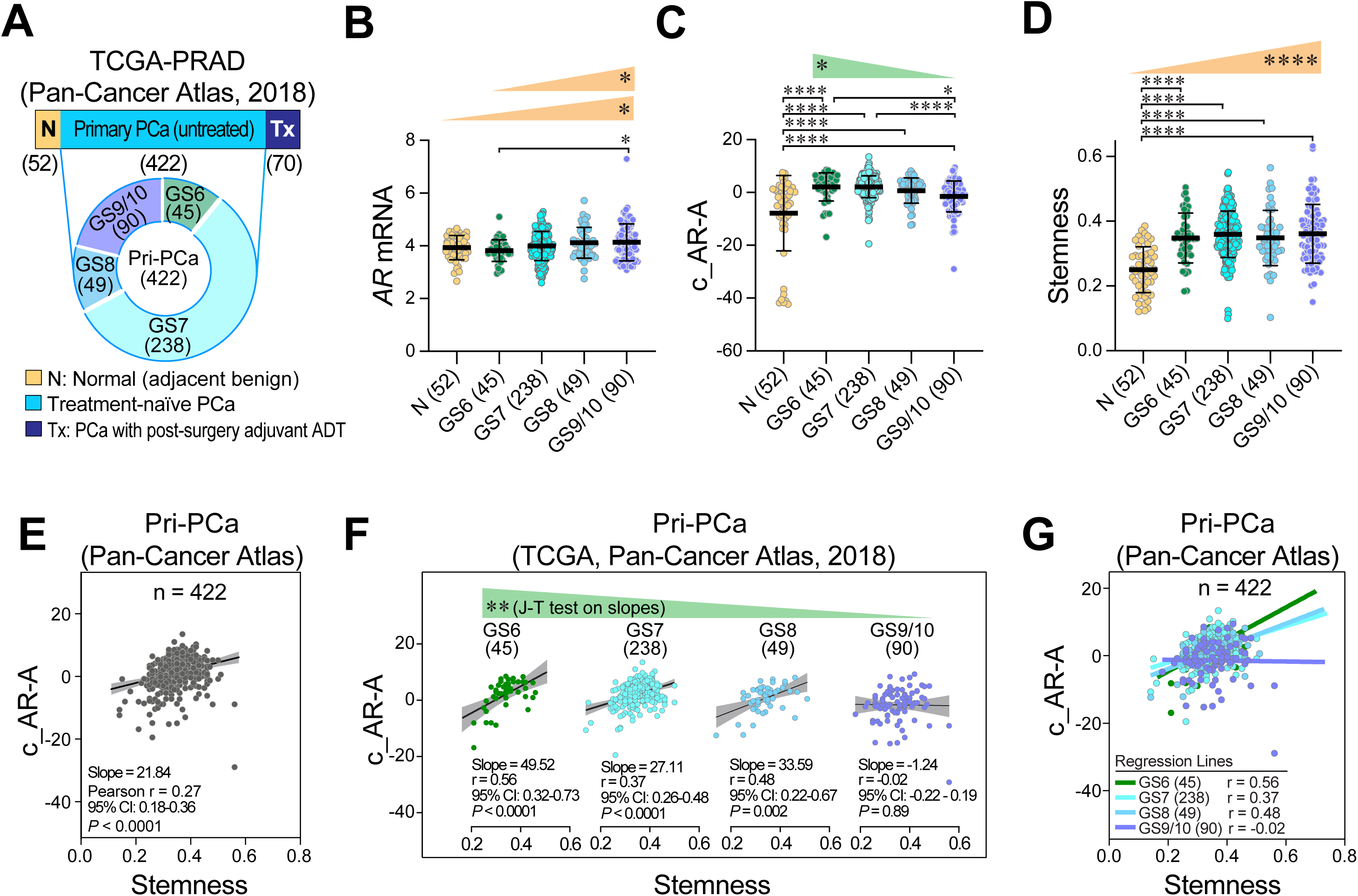
*Continual increases in Stemness with declining c_AR-A during spontaneous PCa progression*. **A,** The pie chart showing sample sizes in different Gleason grades in the TCGA untreated Pri-PCa cohort. **B-D,** Progression of treatment-naïve Pri-PCa is accompanied by decreasing c_AR-A but increasing Stemness. Data is presented as mean ± SD. Shown on top are the Jonckheere-Terpstra’s trend (J-T) test results (wedges). **E-F,** Progression in Pri-PCa is accompanied by a gradual loss of the positive correlation between canonical c_AR-A and Stemness. Scatter plots illustrating the correlation between c_AR-A and Stemness in (**E**) combined Pri-PCa cases in TCGA with an overall linear regression model and (**F**) individual GS grades with corresponding linear regressions showing decreasing slopes from low to high GS grades by J-T test. The slopes of best-fit regression line are also shown. **G,** Merged scatter plot showing the correlation between c_AR-A and Stemness across the 4 GS PCa with each GS represented by a distinct color. The correlation becomes progressively weaker from GS6 to GS9/10, transitioning from positive to nearly zero. The correlation coefficients (r) and p-values for each GS are as follows: GS6 (r = 0.56, *P* < 0.0001), GS7 (r = 0.37, *P* < 0.0001), GS8 (r = 0.48, *P* = 0.022), and GS9/10 (r = -0.02, *P* = 0.89). Linear regression lines are overlaid for each GS, demonstrating the decline in the correlation co-efficiency as the GS increases. All statistical analyses were performed using R and statistical significance was assessed using one-way analysis of variance followed by Tukey’s multiple-comparison test (B-D). J-T trend test in B-D and F was used to calculate the statistical significance of the trend across different groups. *, *P* < 0.05; **, *P* < 0.01; ***, *P* < 0.001; ****, *P* < 0.0001; ns, not significant.

**Figure 3.**
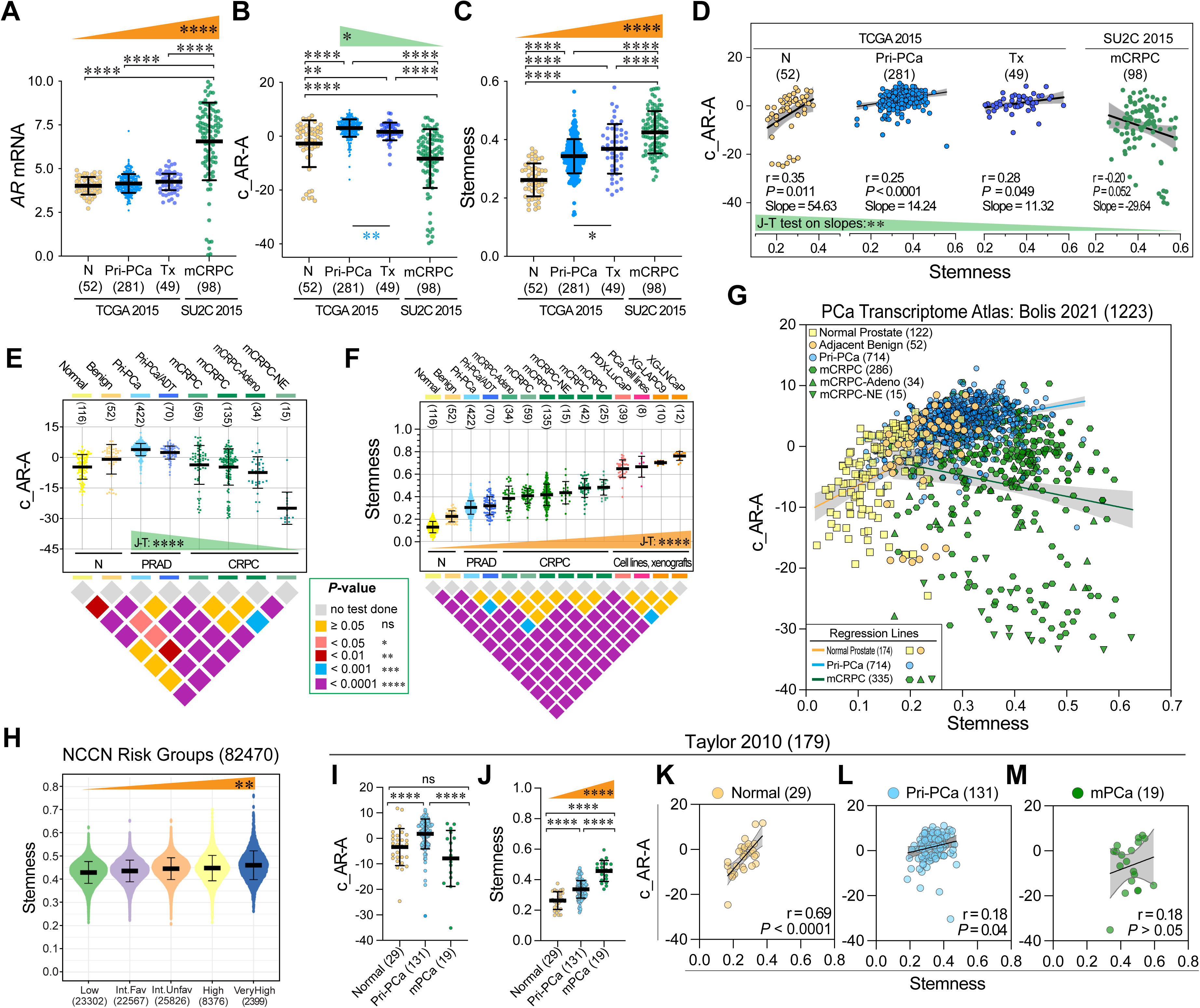
Therapies and metastatic progression drive further divergence between Stemness and canonical c_AR-A with mCRPC showing the highest Stemness but the lowest c_AR-A. **A-D,** Hormonal treatment further drives down c_AR-A but significantly upregulates Stemness. **A-C**, mCRPC displays the lowest pro-differentiation c_AR-A but highest Stemness. Shown are *AR* mRNA levels (**A**), c_AR-A (**B**) and Stemness (**C**) in tumor-adjacent benign tissues (N), treatment-naïve Pri-PCa and advanced Pri-PCa with post-surgery adjuvant ADT (Tx) in TCGA, and mCRPC (SU2C 2015). Sample sizes are indicated in parentheses. Shown on top are the Jonckheere-Terpstra’s (J-T) trend test results (wedges). **D,** Stemness inversely correlates with c_AR-A in mCRPC. Shown are scatter plots with best-fit linear regression lines illustrating the correlation between c_AR-A and Stemness with Pearson correlation coefficient (r), the slopes of lines and J-T test results of slopes indicated below. **E-F,** PCa progression is accompanied by loss of c_AR-A (**E**) but gain of Stemness (**F**) across PCa states. Shown are samples representing the spectrum of PCa development and progression, from normal (organ donors in GTEx) to (tumor-adjacent) benign tissues, to (treatment-naïve) Pri-PCa, aggressive Pri-PCa treated with adjuvant ADT (Pri-PCa/ADT), mCRPC, and, finally, to (clonally-derived) PCa models including PDX, xenografts (XG) and cells lines. From Left to right: Normal (n=116): Normal prostates from GTEx; Benign (n=52): tumor-adjacent benign tissue from TCGA-PRAD; Pri-PCa (n=422): treatment-naïve localized PCa from TCGA-PRAD; Pri-PCa/ADT (n=70): advanced Pri-PCa treated with adjuvant hormone therapy from TCGA-PRAD; mCRPC (n=59): mCRPC from GSE126078; mCRPC (n=135): mCRPC from SU2C 2015 cohort; mCRPC-Adeno (n=34): adenocarcinoma mCRPC from Beltran 2016 cohort; mCRPC-NE (n=15): neuroendocrine mCRPC from Beltran 2016 cohort; mCRPC (n=42): mCRPC from Westbrook 2022; mCRPC (n=25): mCRPC from Alumkal 2020; PDX-LuCaP (n=39): PCa PDXs from GSE126078; PCa cell lines (n=8): PCa cell lines from CCLE; XG-LAPC9 (n=10) and XG-LNCaP (n=12): PCa xenografts from GSE88752. Statistical significances between groups were shown in inverted pyramid grid plot as indicated by different colors. Student’s *t*-test was used to compare each group and adjusted p-value for multiple comparisons was done by Tukey’s range test. See Table S4 for the summary of statistical comparisons. J-T trend test was used to assess the statistical significance of the trend across groups. See also Table S1 for cohort and RNA-seq data access information. **G,** Scatter plots in the Bolis 2021 integrated cohort (normal n=174, Pri-PCa n=714, mCRPC n=335) showing the changing correlation between c_AR-A and Stemness across disease stages: positive in normal to weaker correlations in Pri-PCa to negative correlations in mCRPC. Regression lines are color-coded, and the slope of the best-fit regression line and Pearson’s r are provided: yellow for normal (Pearson’s r=0.42, *P* <0.0001; slope: 43.87, 95%CI, 29.57 to 58.18); blue for Pri-PCa (r=0.32, *P* <0.0001; slope: 15.04, 95%CI, 11.76 to 18.32); and green for mCRPC (r=-0.15, *P* = 0.005; slope: -17.25, 95%CI,-29.24 to -5.26). **H**, Violin plot showing increasing Stemness among the NCCN Risk Groups containing 82470 microarray samples with localized disease in GRID. J-T trend test was used to assess the statistical significance of the trend across groups. **I-M,** Increasing Stemness during PCa progression in the Taylor 2010 cohort consisting of tumor-adjacent benign tissue (Normal; n=29), Pri-PCa (n=131) and metastatic mPCa (n=19). The c_AR-A significantly increased in Pri-PCa vs. Normal and significantly decreased in mPCa compared to Pri-PCa (**I**) whereas the Stemness continued to increase from Benign to Pri-PCa to mPCa (**J**). **K-M**, Scatter plots illustrating the correlations between c_AR-A and Stemness in different contexts with corresponding linear regression lines: **K,** Positive correlation in normal (Pearson’s r=0.69, *P* <0.0001); **L,** a gradual loss of positive correlation between Stemness and c_AR-A during PCa progression (Pearson’s r=0.18, *P* = 0.043); **M,** a loss of positive correlation between the Stemness and c_AR-A at more advanced mPCa stage (Pearson’s r=0.18, *P* =0.47). All data in dot plots and violin plots are shown as mean ± SD. Statistical significance was determined using one-way analysis of variance (ANOVA) followed by Tukey’s multiple-comparison test. Jonckheere-Terpstra’s trend (J-T) test was used to calculate the statistical significance of the trend across different groups. Pearson correlation coefficients (Pearson’s r) were used to calculate the correlations between the Stemness and c_AR-A. Note: ns, not significant; *, *P* < 0.05; **, *P* < 0.01; ***, *P* < 0.001; ****, *P* < 0.0001.

**Figure 4.**
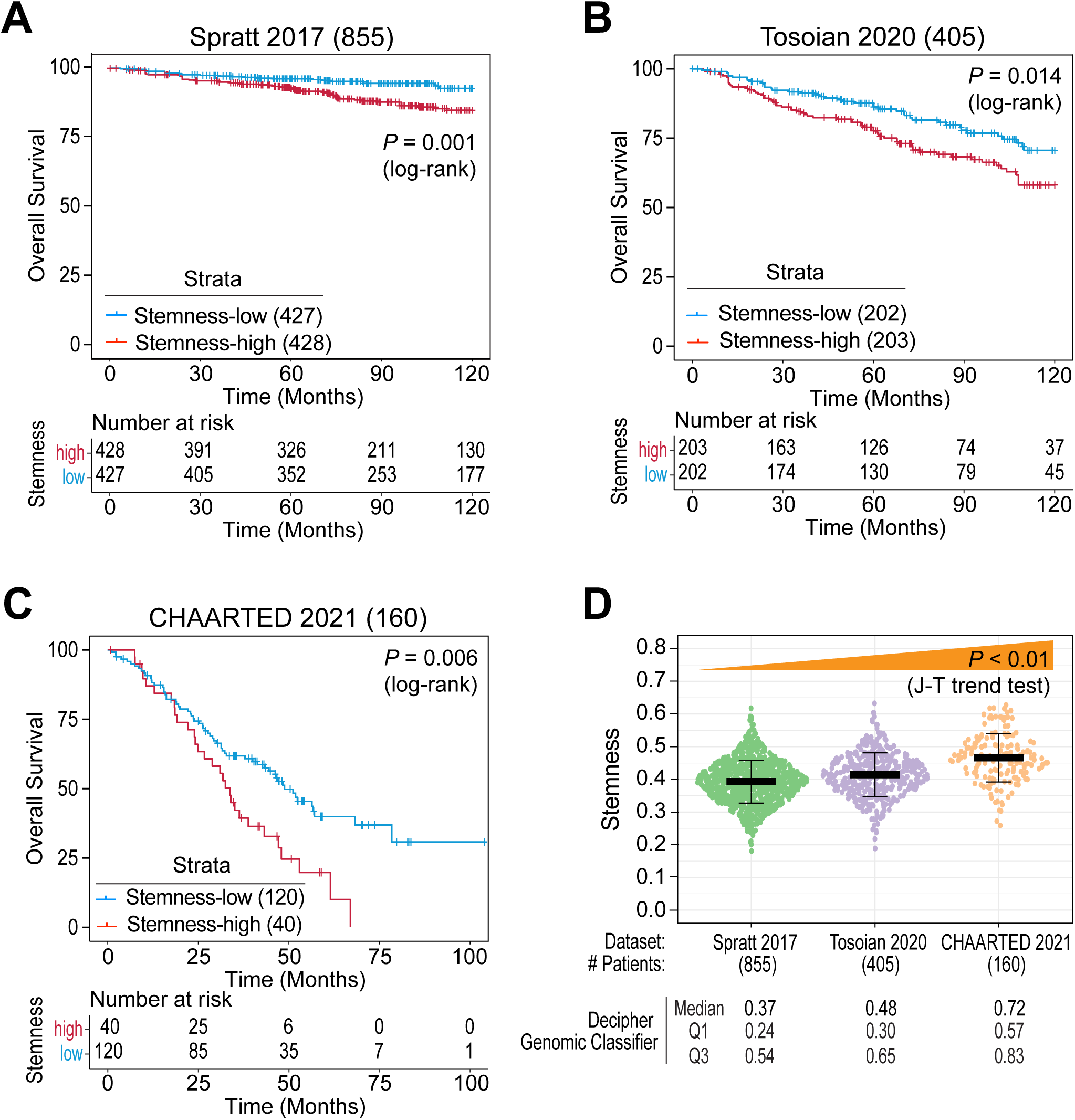
High Stemness is associated with poor patient overall survival. **A-C,** Kaplan-Meier plots showing that high Stemness associates with worse overall survival in PCa patients from Spratt 2017 (**A**), Tosoian 2020 (**B**), and CHAARTED 2021 (**C**) cohorts. *P* value was determined using the log-rank test. **D,** Stemness increases across patient groups with higher Decipher genomic classifier scores. The three cohorts were arranged by their median Decipher genomic classifier scores, and the J-T trend test was used to assess statistical significance of this trend.

For the receiver operating characteristic (ROC) analysis, ROC curves and corresponding area under the curve (AUC) values were computed to evaluate the discriminative performance of the 12-gene PCa-Stem signature and the general Stemness Index. ROC analyses were performed using the pROC package in R (version 4.3.3), with AUC estimated by the non-parametric DeLong method. Classifications included (i) high-grade (GS > 7) versus low-grade (GS ≤ 7) Pri-PCa, and (ii) mCRPC versus treatment-naïve Pri-PCa. All ROC analyses were performed within the PCa Transcriptome Atlas to avoid inter-dataset batch effects.

Unless otherwise specified in the corresponding figure legends, all statistical tests were two-sided, and *P* < 0.05 was considered statistically significant.

## Results

### Adapting and developing the stemness and c_AR-A index scores for PCa analysis

To determine the global and dynamic changes of and inter-relationship between the degree of malignancy (oncogenic dedifferentiation) and level of (luminal) differentiation during PCa development and progression (Fig. S1A-B), we employed transcriptome-based algorithms to annotate 87,339 transcriptomic profiles from 33 datasets, including 3,153 bulk RNA-seq and 84,081 microarray samples, together with single-cell and spatial transcriptomic datasets, encompassing the PCa continuum (Fig. S1B; Table S1). In our studies, we focused on and compared three progressive stages: from normal prostate to localized Pri-PCa, from low-grade to high-grade Pri-PCa, and from treatment-naïve Pri-PCa to CRPC/mCRPC (Fig. S1A). We defined “progression” as stages beyond Pri-PCa and as disease entities that are more aggressive in a comparative manner. Specifically, we adapted the Stemness Index [6] (mRNAsi; **Stemness** for short) (Fig. S1C) to gauge the degree of malignancy (or aggressiveness) and employed the Z-score normalized canonical AR activity or **c_AR-A** (Fig. S1D) to measure the level of differentiation. Canonical c_AR-A can be determined by quantifying AR-regulated transcripts under intact androgen/AR signaling conditions, and there are several such AR-regulated gene lists [22, 23, 60, 74, 75] (Fig. S2A-B). We validated the c_AR-A list based on their enrichment in prostatic epithelium differentiation, correlation of individual gene expression with c_AR-A signatures in clinical datasets, and overall consistency in correlation among the AR-A scores (Fig. S2B-D). Given that Bluemn’s c_AR-A score [60] correlated well with other scores (Fig. S2E-G) and the 10 genes in this signature consistently represented AR targets in multiple clinical datasets (Fig. S2D; Table S2), we decided to use this AR-regulated gene list for our c_AR-A quantification.

### Normal prostate epithelium shows a negative correlation between Stemness and c_AR-A

The normal human prostate (NHP) comprises 2 major epithelial types: basal and luminal cells (Fig. S3A). We annotated two studies that profiled FACS-purified NHP luminal and basal cell populations [19, 21] and observed that the luminal cell population (Trop2^+^CD49f^lo^) possessed higher *AR* mRNA levels and c_AR-A but lower Stemness compared to basal cells (Trop2^+^CD49f^hi^) and, in both datasets, c_AR-A negatively correlated with Stemness (Fig. S3B-C). These results support that the NHP basal cell compartment harbors primitive stem cells [7, 19, 21, 76, 77] and that c_AR-A measures the level of prostate epithelial differentiation.

### Pri-PCa exhibits concordantly increased Stemness and c_AR-A, and neoadjuvant ADT (nADT) decreases both Stemness and c_AR-A

In analyzing c_AR-A and the Stemness in matched pairs of prostate tumor (T) and normal (N; tumor-adjacent benign) tissues in the TCGA-PRAD [22] (n=51 pairs), Wyatt 2014 [31] (n=12 pairs), and Long 2020 [28] (n=10 pairs) cohorts, we observed concordantly increased c_AR-A and Stemness (Fig. 1A-C). Accompanying elevated c_AR-A, *AR* mRNA levels were also increased in Pri-PCa compared to benign tissue (Fig. 1A-C; note that increase in *AR* mRNA levels in the Long 2020 cohort approached statistical significance). The paired T/N comparisons (Fig. 1A-C) suggest that early PCa growth is characterized by concordantly increased Stemness and c_AR-A. To test whether c_AR-A might causally drive increased Stemness during early prostate tumorigenesis, we investigated 4 PCa nADT (i.e., ∼2-6 months of ADT before prostatectomy or radiation) cohorts [26–28]. In the first 3 cohorts with matched pairs of pre-and post-nADT samples (from same patients), nADT concordantly downregulated c_AR-A (without decreasing *AR* mRNA levels) as well as the Stemness (Fig. 1D-F). In the Nastiuk cohort with non-paired post-nADT and no-nADT samples (i.e., from different patients but matched for ages and tumor grades and stages), nADT increased *AR* mRNA levels (Fig. S4A) but reduced both c_AR-A and Stemness (Fig. S4B-C).

### Pri-PCa progression is accompanied by reduced c_AR-A but continually increasing Stemness

We next assessed the dynamic changes of Stemness and c_AR-A during *spontaneous* (i.e., therapy-free) Pri-PCa progression (Fig. 2). To this end, we sorted out treatment-naïve tumors in TCGA-PRAD by excluding samples from patients who received adjuvant (post-surgery) hormone therapy (i.e., Pri-PCa/ADT or Tx; n=70) (Fig. 2A) and grouped the Pri-PCa samples according to increasing GS (Fig. 2B-D). When compared with the 52 benign (N) samples, all the four GS groups from Pri-PCa (GS6, GS7, GS8, and GS9/10 combined) showed higher c_AR-A and Stemness (Fig. 2C-D), consistent with the results in pairwise T/N comparisons (Fig. 1A-C). However, when compared to GS6 PCa, the three higher GS groups showed increasing *AR* mRNA levels (Fig. 2B) but decreasing c_AR-A (Fig. 2C) and increasing Stemness (Fig. 2D). Jonckheere-Terpstra’s (J-T) trend test validated a gradual reduction in canonical AR signaling but an increasing trend of Stemness with Pri-PCa progression (Fig. 2C-D). Notably, although we observed a modest positive correlation between c_AR-A and Stemness in pooled Pri-PCa samples (Pearson’s r=0.27; Fig. 2E), this positive correlation gradually diminished with PCa progression when we categorized the samples by tumor grade (Fig. 2F). The linear regression analysis confirmed that the interaction between Stemness and c_AR-A significantly depended on tumor grade (*P*=0.0003), with a linear decrease in the slope of the correlation as GS increased (Fig. 2G).

Collectively, these results indicate that, in contrast to early tumorigenesis, *spontaneous PCa progression (i.e., with increasing GS and tumor grade) is marked by a divergence between oncogenic Stemness and pro-differentiation c_AR-A*.

### Therapies and metastatic progression drive further divergence between attenuated c_AR-A and increased Stemness

Next, we asked how clinical treatment and metastatic progression may impact the Stemness and c_AR-A by querying untreated Pri-PCa and Pri-PCa/ADT from TCGA and mCRPC in SU2C [10] (Fig. 3A-C). c_AR-A was elevated in Pri-PCa compared to adjacent benign tissue (N) but reduced in more aggressive Pri-PCa/ADT samples, while the Stemness, in contrast, continually and persistently increased from N → Pri-PCa → Pri-PCa/ADT (Fig. 3B-C). Strikingly, mCRPC, despite markedly upregulating *AR* mRNA (Fig. 3A), showed the lowest c_AR-A but highest Stemness (Fig. 3B-C), *indicating a shift from a positive correlation between* c_*AR-A and Stemness in Pri-PCa to a negative association in mCRPC* (Fig. 3D).

By employing normalized RNA-seq data from the Bolis 2021 PCa transcriptome atlas [36], which included normal prostate specimens (n = 174), Pri-PCa (n = 714) and mCRPC (n = 335), we tracked the Stemness and c_AR-A across the PCa spectrum (Fig. 3E). c_AR-A increased from benign to Pri-PCa but declined in Pri-PCa/ADT and continued to decrease in mCRPC, reaching its lowest in mCRPC-NE [41] as supported by J-T test (Fig. 3E). In contrast, the Stemness exhibited a continually increasing trajectory across the PCa spectrum encompassing Normal/Benign → treatment-naïve Pri-PCa → Pri-PCa/ADT → (m)CRPC (*P*<0.0001, J-T Trend test; Fig. 3F; Fig. S5A). Notably, the 3 subtypes of mCRPC, i.e., AR-positive (ARPC), CRPC-NEPC and AR^-^/NE^-^double-negative PCa (DNPC) all showed higher Stemness than Pri-PCa and displayed increasing Stemness among the 3 subtypes (Fig. S5B). Unexpectedly, tumor-adjacent benign prostate tissues showed much higher Stemness than normal prostate in the Genotype-Tissue Expression Project (GTEx) [18] (Fig. 3F; Fig. S5A), which were all obtained from cancer-free organ donors. This surprising finding suggests that the transcriptomes of adjacent benign tissues have apparently skewed towards those of cancer samples, consistent with previous observations that adjacent benign tissues represent an intermediate state between healthy and tumor tissues [78]. Also of interest, the Cancer Cell Line Encyclopedia (CCLE) [50, 79] cell lines including PCa cell lines, and PCa-derived PDX [46] and xenografts (XG) [51] exhibited the highest Stemness (Fig. 3F; Fig. S5A), supporting that cancer cell lines and xenografts are clonally derived from and highly enriched in cancer stem cells (CSCs).

As in our early dissection of the relationship between c_AR-A and Stemness during spontaneous Pri-PCa progression (Fig. 2F-G), analysis of the Bolis 2021 PCa transcriptome atlas [36] revealed a similar change from a positive correlation between c_AR-A and Stemness in Pri-PCa (blue line; Pearson’s r=0.32, *P*<0.0001; Fig. 3G) to a negative correlation between the two in mCRPC (green line; Pearson’s r=-0.15, *P*=0.005; Fig. 3G). We further extended our Stemness and c_AR-A interrogation to include microarray-derived transcriptomic datasets [35, 47], allowing us to access data from PCa cohorts generated before the next-generation sequencing (NGS) era. Analyzing the GRID database of over 82,000 prospectively collected biopsy samples of localized PCa from clinical use of the Veracyte Decipher test [35], we systematically quantified the Stemness across the NCCN Risk Groups and observed increasing Stemness along the NCCN Risk trajectory (Fig. 3H). Likewise, in the Taylor dataset [47], we observed that c_AR-A was the highest in Pri-PCa but significantly decreased in metastatic PCa (mPCa) (Fig. 3I) while the Stemness increased progressively from benign to Pri-PCa and reached the highest levels in mPCa (Fig. 3J). Linear regression analysis confirmed a gradual loss of positive correlation between c_AR-A and Stemness during PCa progression, as illustrated by distinct regression lines for each stage (Fig. 3K-M).

Finally, we analyzed the changes of Stemness and c_AR-A in genetically engineered mouse models (GEMMs) of PCa with increasing aggressiveness due to individual or combined deletion of 3 tumor suppressor genes, *Pten*, *Rb1* and *Trp53* [49] (Fig. S5C-H). Briefly, *Pten*^-/-^single knockout (SKO) prostate tumors develop around 9-12 weeks, and mice rarely develop metastasis with a median lifespan of 48 weeks. In contrast, the *Pten*^-/-^;*Rb1*^-/-^double knockout (DKO) mice develop highly metastatic PCa that shortens median survival to ∼38 weeks whereas the triple KO (TKO; *Pten*^-/-^*;Rb1*^-/-^*;Trp53*^-/-^) tumors are exclusively AR^−^ NEPC and castration-resistant *de novo* with high metastatic rate and lifespan of ∼16 weeks [49]. We found that the c_AR-A was the highest in SKO tumors but much reduced in aggressive DKO and TKO tumors (Fig. S5C). In contrast, Stemness was the lowest in SKO but significantly increased in DKO and TKO (Fig. S5D). Correlation analyses revealed a positive correlation between c_AR-A and Stemness in SKO that turned negative in both DKO and TKO tumors (Fig. S5E-H).

The above results, collectively, indicate that *hormone therapy and metastatic progression further drive marked increases in Stemness with concomitant decreases in* c_*AR-A*.

### The general Stemness score is prognostic of poor survival in PCa patients

We investigated global Stemness and its prognostic significance, and observed that high Stemness consistently correlated with worse patient survival across various cohorts including the Spratt 2017 [32] (Fig. 4A), Tosoian 2020 [33] (Fig. 4B), CHAARTED 2021 [34] (Fig. 4C), and the mCRPC (SU2C 2019 [48]; Fig. S6A-B) as well as Taylor 2010 [47] (Fig. S6C-D). Notably, the CHAARTED cohorts displayed the highest mean Stemness compared to Tosoian and Spratt cohorts, and the highest median Decipher Prostate Genomic Classifier [32–34] among the three cohorts (Fig. 4D). Interestingly, like high Stemness, low c_AR-A was correlated with poor patient survival in the Taylor dataset (Fig. S6E).

### Stemness-high PCa are associated with aggressive molecular subtypes, proliferative programs, and lineage-plastic states

To identify key determinants of global Stemness and explore differences between tumors with high versus low Stemness scores, we stratified Pri-PCa (TCGA-PRAD) and mCRPC (SU2C 2019) samples separately based on Stemness scores (Fig. 5A). Differential gene expression (DEG) analyses revealed significant differences between the top 33% (Stemness-high) and the bottom 33% (Stemness-low) PCa samples in both treatment-naïve Pri-PCa and treatment-failed mCRPC (Fig. 5B-C; Table S3).

**Figure 5.**
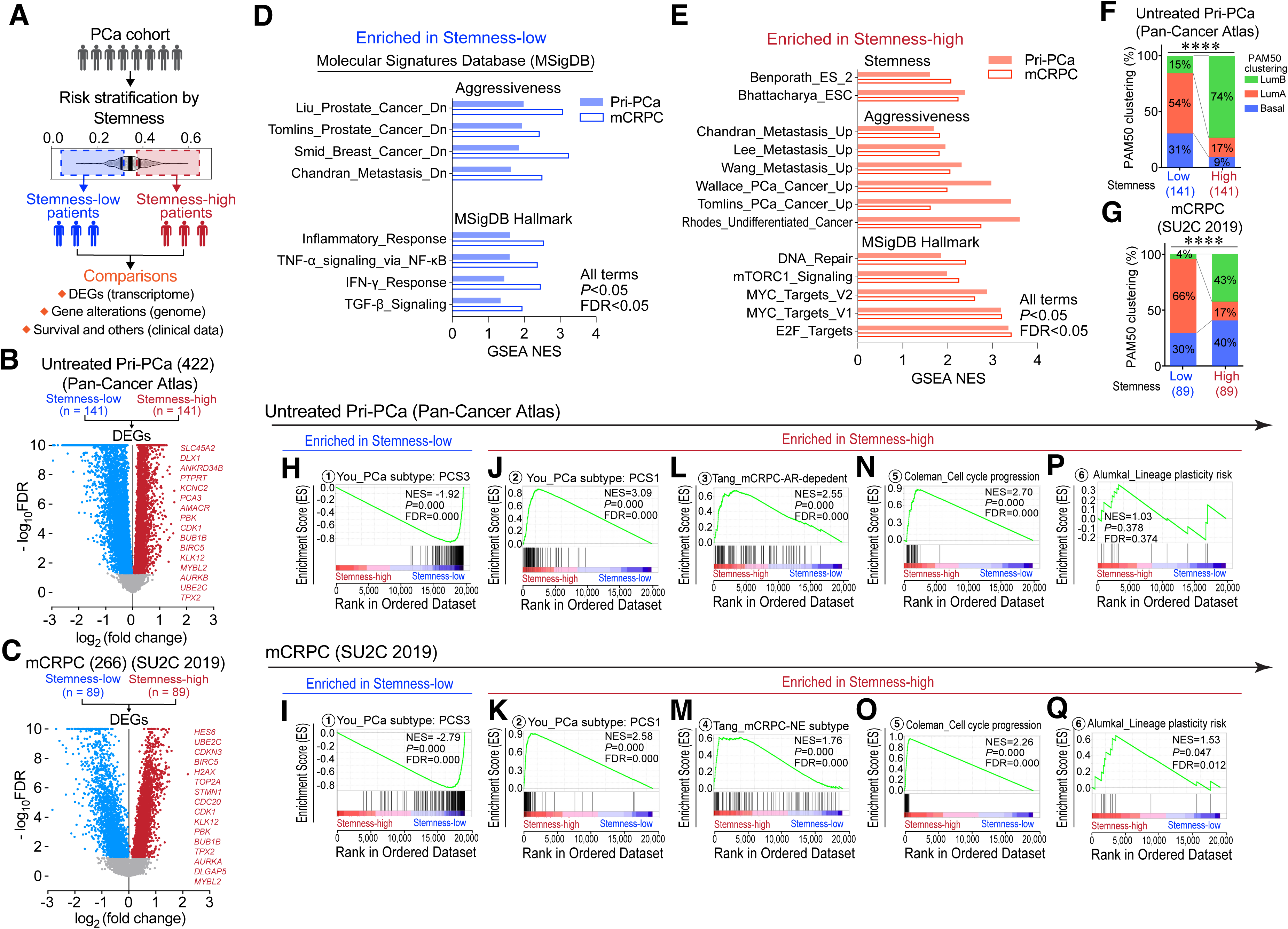
Stemness-high PCa are associated with aggressive molecular subtypes and a newly developed 12-gene PCa-Stem signature prognosticates poor patient survival. **A,** Schematic illustrating the profiling strategy used to assess associations between Stemness and molecular features in PCa. Patients were ranked by Stemness score and stratified into Stemness-high (top 33%) and Stemness-low (bottom 33%) groups for downstream analyses. Genome-wide functional analyses were performed using transcriptomic, genomic, and clinical data. **B-C,** Volcano plots showing differentially expressed genes (DEGs) between Stemness-high and Stemness-low groups in Pri-PCa (**B**) and mCRPC (**C**) cohorts. Red and blue indicate upregulated and downregulated DEGs, respectively. Genes with FDR > 0.05 are shown in gray. Sample sizes (n) of each group are indicated. See Table S3 for complete DEG lists. **D-E,** GSEA showing low Stemness is associated with immune signaling and inflammatory response (**E**) while high Stemness in both cohorts is enriched for embryonic stem cell traits, aggressiveness, DNA repair, MYC activation, and mTORC1 signaling (**F**). All enrichments are significant (*P* < 0.05, FDR < 0.05). **F-G,** PAM50 subtyping showing a higher frequency of aggressive LumB subtype in Stemness-high versus Stemness-low samples in Pri-PCa (**G**) and mCRPC (**H**). (****, *P* < 0.0001, χ^2^ test). **H-K,** GSEA showing enrichment of the aggressive PCS1 signature in Stemness-high groups and the less aggressive PCS3 signature in Stemness-low groups in Pri-PCa (**H**, **J**) and mCRPC (**I**, **K**). The Prostate Cancer Subtype (PCS) classification system stratifies tumors into three molecular subtypes based on transcriptomic features: PCS1 (most aggressive), PCS2 (intermediate), and PCS3 (least aggressive). See Table S2 for PCS gene signature. **L-Q,** GSEA showing AR signature enrichment in Pri-PCa Stemness-high (**L**), NEPC signature enrichment in mCRPC Stemness-high (**M**), and shared enrichment of cell cycle (**N**, **O**) and lineage plasticity (**P**, **Q**) programs in both cohorts. See Table S5 for gene sets.

Gene set enrichment analysis (GSEA) revealed that Stemness-high PCa samples were enriched in gene sets associated with embryonic stem cells (ESC) as well as aggressive cancer phenotypes such as undifferentiated cancer and metastasis in both Pri-PCa and mCRPC whereas Stemness-low PCa samples were enriched in gene signatures linked to lower aggressiveness and inflammatory pathways (Fig. 5D-E). Additionally, hallmark pathways such as E2F targets, MYC targets, mTORC1 signaling, and DNA repair were upregulated in Stemness-high PCa (Fig. 5E), while TGF-β signaling, TNF-α signaling, and IFN-γ responses were prominent in Stemness-low PCa (Fig. 5D).

Subsequent PCa molecular subtype analysis revealed that the aggressive PAM50-LumB molecular subtype [80–82] was significantly more prevalent in Stemness-high PCa, accounting for 74% in Pri-PCa (compared to 15% in Stemness-low PCa; Fig. 5F) and 43% in mCRPC (compared to 4% in Stemness-low PCa; Fig. 5G). The PAM50-Basal subtype [80, 81] was also increased in Stemness-high mCRPC but not in Pri-PCa (Fig. 5F-G), aligning with the characteristics of mCRPC, which involves lineage plasticity and acquisition of basal, mesenchymal, neural and stem-like phenotypes when developing ARPI resistance [7, 82, 83]. The PCS1 subtype [84, 85], known for its association with ADT resistance and the highest risk of progression to advanced disease in comparison with PCS2 or PCS3, was enriched in both Stemness-high Pri-PCa and mCRPC whereas PCS3 signature was enriched in Stemness-low Pri-PCa and mCRPC (Fig. 5H-K). Moreover, in PCa subtypes epigenetically stratified by chromatin accessibility [86], Stemness-high samples remained largely AR-dependent in Pri-PCa (Fig. 5L) but transitioned to AR-independent NE subtype in mCRPC (Fig. 5M). Of interest, Stemness-high PCa was characterized by high proliferation in both Pri-PCa and mCRPC as evidenced by their enrichment in a 31-gene cell-cycle progression signature [82] (Fig. 5N-O; Table S5). Finally, we examined the association between Stemness and a signature of lineage plasticity risk in mCRPC after Enza treatment [12] and found that Stemness-high tumors was at risk for this virulent form of treatment resistance (Fig. 5P-Q).

Taken together, these findings indicate that *PCa with high Stemness exhibit increased aggressiveness, enhanced proliferative signaling, lineage plasticity, and therapy-resistant transcriptional programs, and are associated with poor patient survival*.

### The 12-gene PCa-Stem signature defines a coordinated proliferative program with functional and prognostic relevance

To derive a PCa-specific transcriptional measure of Stemness, we intersected genes significantly upregulated (FC ≥ 2, FDR < 0.05) in Stemness-high versus Stemness-low tumors in both treatment-naïve Pri-PCa and mCRPC (Fig. 5B-C). This analysis identified a shared **12-gene PCa-Stem signature** comprising *HMMR*, *AURKB*, *CENPA*, *DEPDC1B*, *HJURP*, *PBK*, *MELK*, *UBE2C*, *DLGAP5*, *NEK2*, *BIRC5*, and *KLK12* (Fig. 6A-B). PCa-Stem signature scores increased progressively across the PCa continuum and correlated strongly with PCa disease progression scores (pseudotime) derived from trajectory inference analysis of the Bolis PCa transcriptome atlas [36] (Pearson’s r = 0.72; Fig. 6C). Strikingly, gene expression analysis revealed that 11 of the 12 genes (except *KLK12*) in the PCa-Stem signature exhibited stage-related increases and positively correlated with the PCa disease progression (Fig. S7A-B). In contrast, among the genes commonly downregulated in Stemness-high PCa samples in both Pri-PCa and mCRPC (Table S3), some (e.g., *PTGDS*, *SPARCL1*, *CLU*, *GJA1* and *S100A4*) were involved in regulating prostate-specific glandular structures and luminal functions while others (e.g., *FBLN1*, *SFRP1*, *IGFBP6*, *GAS1* and *TIMP2*) played general roles in epithelial differentiation (Fig. S7C). The PCa-Stem signature also showed enhanced discrimination of high-versus low-grade Pri-PCa and mCRPC versus treatment-naïve Pri-PCa compared with the global Stemness Index (AUC = 0.662 and 0.865, respectively; Fig. S7D– E).

**Figure 6.**
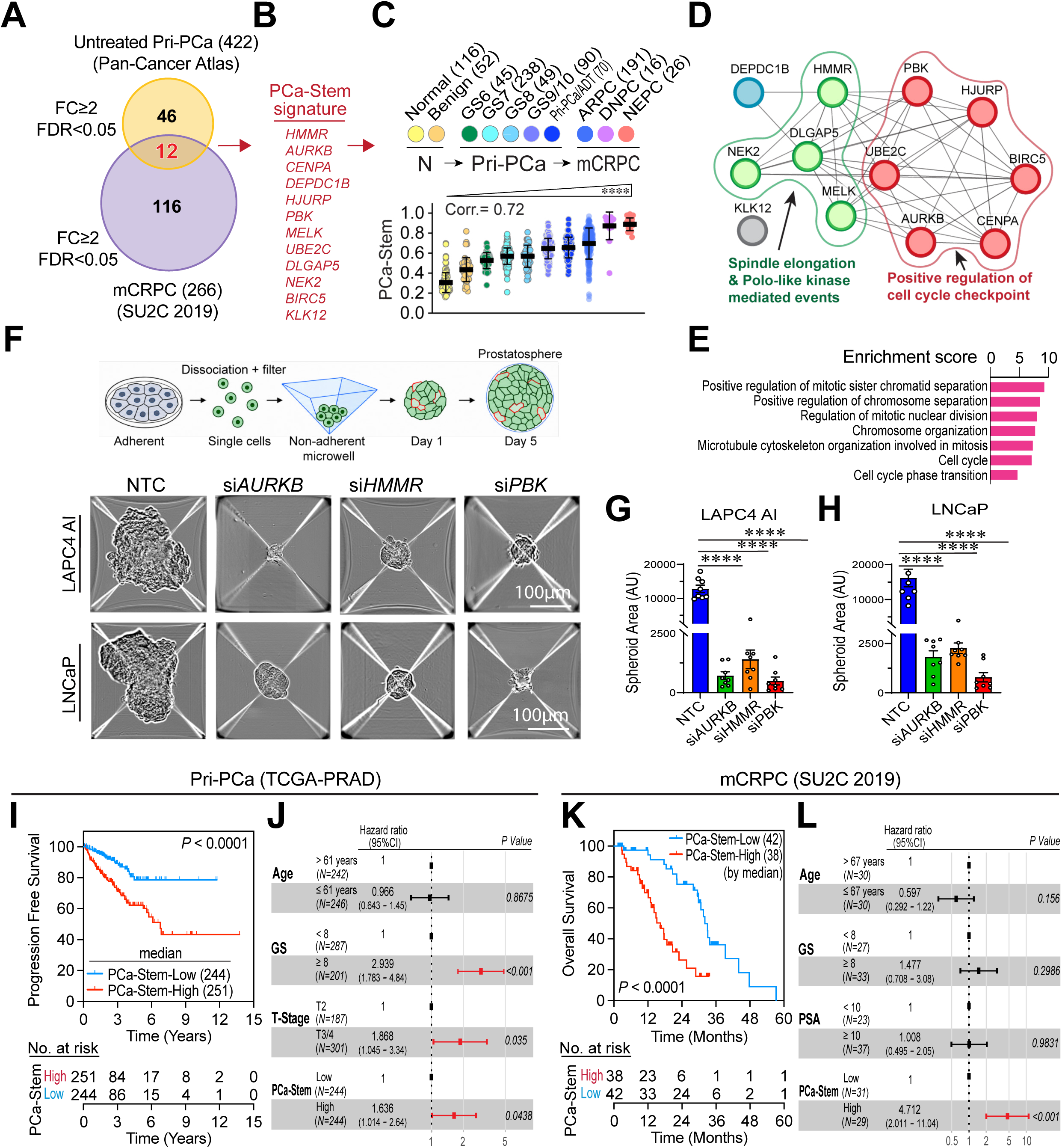
Derivation, biological characterization, functional validation, and clinical significance of the 12-gene PCa-Stem signature. **A-B,** Derivation of the 12-gene PCa-Stem signature by overlapping genes significantly upregulated in Stemness-high versus Stemness-low tumors from both Pri-PCa and mCRPC cohorts (fold change ≥ 2, FDR < 0.05). **A,** Venn diagram showing the overlap. **B,** The resulting 12-gene PCa-Stem signature. **C,** Upper panel, schematic representation of the PCa progression continuum analyzed in this study. Sample sizes are indicated for each disease stage, and the corresponding color scheme is used throughout the progression analyses. Lower panel, dot plot showing progressively increasing PCa-Stem signature scores across the PCa progression continuum. Pearson’s correlation coefficient (Corr.) indicates the association between PCa progression scores, derived from the integrated PCa transcriptomic continuum (Bolis et al., 2021), and PCa-Stem signature scores. Data are presented as mean ± SD, and trends across disease stages were assessed using the Jonckheere–Terpstra trend test. **D,** STRING protein–protein interaction (PPI) network of the 12-gene PCa-Stem signature. Eleven of the twelve PCa-Stem signature genes formed a highly interconnected interaction network, whereas KLK12 was relatively isolated because of its secreted extracellular protease function. Functional clustering identified two major biological modules associated with positive regulation of the cell-cycle checkpoint (red) and spindle elongation/mitotic progression (green), supporting coordinated regulation of mitotic progression and proliferative Stemness. **E,** Functional enrichment analysis of the 12-gene PCa-Stem signature. Shown is the Gene Ontology (GO) biological process analysis. Enrichment scores are presented as −log₁₀-transformed FDR values. **F–H,** Functional validation of PCa-Stem signature genes using AggreWell-based prostatosphere assays in LAPC4-AI and LNCaP cells following siRNA-mediated knockdown of *HMMR*, *AURKB*, or *PBK*. **F,** Schematic of the AggreWell-based prostatosphere assay workflow (top) and representative images of prostatospheres generated from NTC-, siAURKB-, siHMMR-, and siPBK-transfected LAPC4-AI and LNCaP cells at day 5 (bottom). Scale bars, 100 μm. Additional representative prostatosphere images are shown in Fig. S9I-J. **G–H,** Quantification of prostatosphere area in LAPC4-AI (**G**) and LNCaP (**H**) cells. Each dot represents an individual prostatosphere formed within a single AggreWell microwell. Data are presented as mean ± SD. ****, *P* < 0.0001. **I–L,** Kaplan-Meier analyses showing that the 12-gene PCa-Stem signature predicts worse progression-free survival (PFS) in TCGA Pri-PCa (**I**) and worse overall survival (OS) in SU2C mCRPC (**K**). Multivariable Cox models adjusting for clinicopathologic variables confirm the high PCa-Stem signature score as an independent prognostic factor for PFS (**J**) and OS (**L**).

Notably, 11 of the 12 PCa-Stem signature genes (except *KLK12*), i.e., *HMMR* (Hyaluronan Mediated Motility Receptor), *AURKB* (Aurora Kinase B), *CENPA* (Centromere Protein A), *DEPDC1B* (DEP Domain Containing 1B), *HJURP* (Holliday Junction Recognition Protein), *PBK* (PDZ-Binding Kinase), *MELK* (Maternal Embryonic Leucine Zipper Kinase), *UBE2C* (Ubiquitin Conjugating Enzyme E2 C), *DLGAP5* (DLG Associated Protein 5), *NEK2* (NIMA Related Kinase 2), and, *BIRC5* (Baculoviral IAP Repeat Containing 5), have all been implicated in functionally regulating cell proliferation, cell division, cell-cycle timing and control, mitosis and mitotic control, chromosome movement and stability, and G2/M checkpoint control. Therefore, we asked whether the PCa-Stem signature genes might constitute a coordinated proliferation-associated program, and we found that, PCa-Stem signature proteins indeed formed a highly interconnected STRING network, with *KLK12*, a secreted protease, remaining relatively isolated (Fig. 6D). Functional enrichment converged on mitotic cell-cycle regulation, including chromosome segregation, spindle assembly, and mitotic checkpoint control, with enrichment of AURORA A/B and FOXM1 pathways (Fig. 6E; Fig. S8A-C). Network-based visualization further demonstrated extensive connectivity among these functional processes (Fig. S8D). Extending the network to established stemness and lineage plasticity regulators revealed extensive functional connectivity between the PCa-Stem signature genes and canonical stemness programs (Fig. S8E). An IPA-inferred hierarchical network further connected upstream RB1/TP53, E2F, FOXM1, and MYC-family regulators through plasticity-associated intermediates—including YAP1, TCF4, ASCL1, NOTCH3, and LEF1—to the downstream PCa-Stem signature genes (Fig. S8F), and broader stemness-associated regulatory networks (Fig. S8G). Together, these analyses support the PCa-Stem signature as a coordinated proliferative Stemness program rather than a collection of unrelated genes.

Next, we functionally interrogated three representative PCa-Stem signature genes, *AURKB*, *HMMR*, and *PBK*. AURKB and PBK represent closely connected mitotic regulators within the PCa-Stem network (Fig. 6D; Fig. S8) whereas HMMR, well-known to regulate cell invasion and spindle elongation (Fig. 6D), has also been reported as a marker and functional regulator of stemness-associated lineage plasticity and neuroendocrine differentiation in advanced PCa [57]. Efficient siRNA-mediated depletion of each gene was confirmed at both mRNA and protein levels in androgen-dependent LNCaP and androgen-independent LAPC4-AI cells (Fig. S9A–D). In LAPC4-AI cells, knockdown of these genes impaired invasive and clonogenic phenotypes and altered three-dimensional spheroid growth and cytotoxicity (Fig. S9E–H). Importantly, AggreWell-based prostatosphere assays demonstrated that depletion of *HMMR*, *AURKB*, or *PBK* markedly reduced prostatosphere growth in both LAPC4-AI and LNCaP cells (Fig. 6F–H; Fig. S9I–J), providing functional support for their contribution to stemness-associated growth across distinct PCa contexts. Consistent with their roles in mitotic regulation (Fig. 6D), pH3-S10/tubulin co-staining revealed altered mitotic spindles and microtubule organization following *AURKB*, *HMMR*, or *PBK* depletion in both cell models (Fig. S9K). Interestingly, preliminary studies showed that their knockdown was also associated with reduced γ-H2AX levels in both cell lines (Fig. S9L), potentially linking PCa-Stem signature genes to DNA damage and repair signaling. Collectively, these functional assays support a role for individual components of the PCa-Stem signature in conferring aggressive behavior and stemness-associated traits in PCa cells.

Finally, we observed that elevated PCa-Stem signature scores predicted adverse clinical outcomes in both localized and advanced disease. Briefly, PCa-Stem-high Pri-PCa, compared to PCa-Stem-low Pri-PCa, showed shorter progression-free survival after multivariable adjustment (HR = 1.636, 95% CI 1.014– 2.640, *P* = 0.0438) whereas PCa-Stem-high mCRPC showed much shorter overall survival than PCa-Stem-low mCRPC patients (HR = 4.712, 95% CI 2.011–11.040, *P* < 0.001) (Fig. 6I–L).

These transcriptomic, network, functional, and clinical analyses, in aggregate, suggest that our newly developed 12-gene PCa-Stem signature reports the proliferative aggressiveness of PCa cells and track poor patient survival.

### Single-cell and spatial transcriptomic analyses localize the PCa-Stem program to lineage plasticity-related epithelial cells and reveal its progressive expansion during PCa progression

To determine whether the PCa-Stem program exhibits spatial organization within prostate tumors, we first analyzed Visium spatial transcriptomic data containing the pathologist-validated adjacent AR pathway-positive prostate adenocarcinoma (i.e., ARPC) and de novo NEPC regions (Fig. 7A). We observed that global Stemness (mRNAsi) was preferentially enriched within the NEPC region relative to the adjacent ARPC region (Fig. 7A–B). Consistently, PCa-Stem scores showed similar spatial enrichment and positively correlated with mRNAsi across individual spatial spots (Fig. 7C–D), supporting spatial concordance between the PCa-specific and transcriptome-wide global Stemness programs.

**Figure 7.**
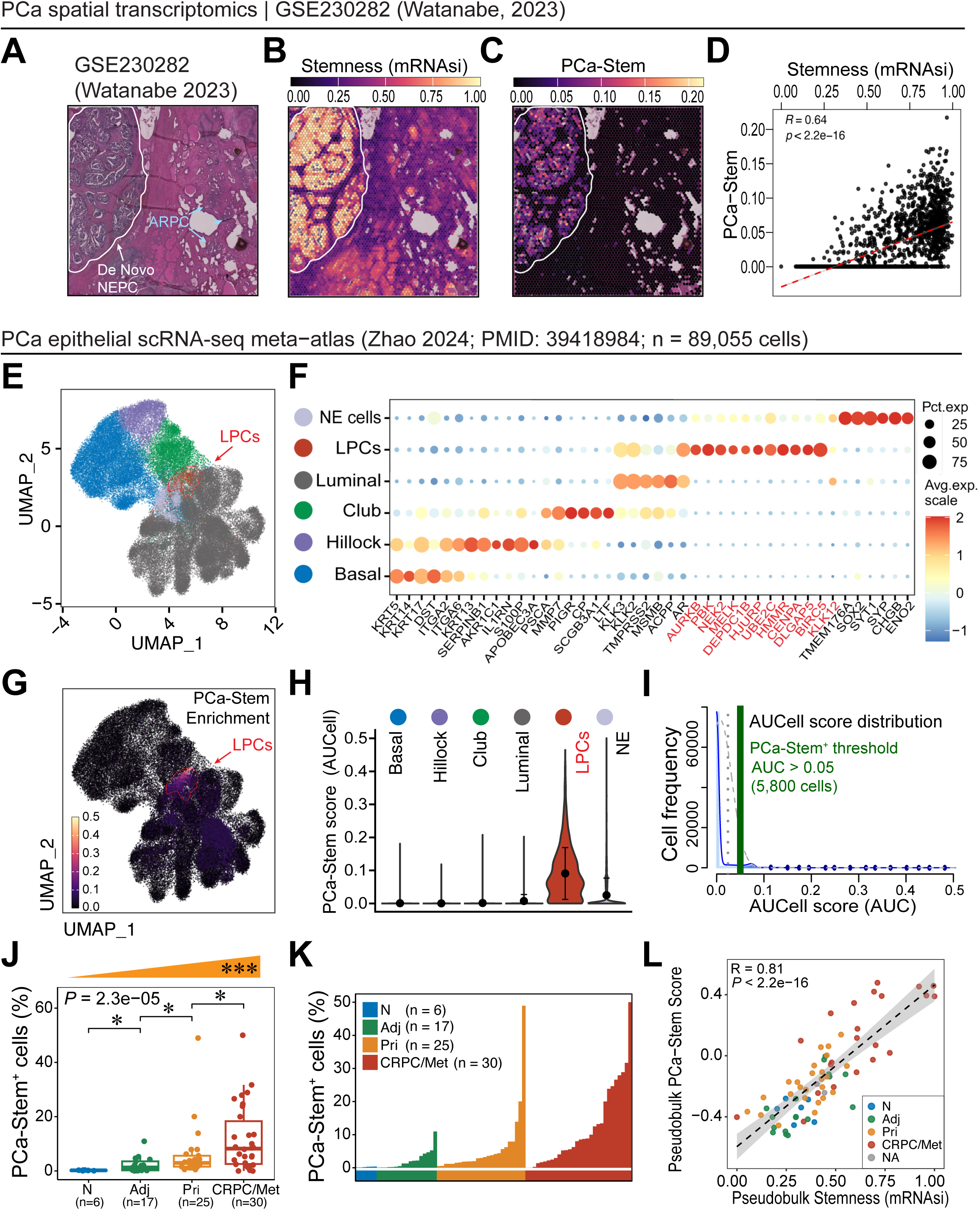
Spatial and single-cell transcriptomic analyses localize the PCa-Stem program to lineage plasticity-related epithelial cells and reveal its progressive enrichment during PCa progression. **A–D,** Spatial distribution of Stemness in a human PCa tissue section containing the pathologist-validated adjacent de novo neuroendocrine PCa (NEPC) and AR pathway-positive prostate adenocarcinoma (ARPC) regions (GSE230282; Watanabe, 2023; ref. 56). **(A)** Histopathological annotation of the tissue section showing de novo NEPC and ARPC regions. **(B–C)** Spatial distribution of the transcriptome-based Stemness Index (mRNAsi) (**B**) and PCa-Stem score (**C**). **(D)** Correlation between PCa-Stem and mRNAsi scores across individual spatial spots (n = 4,397 spots). Pearson’s correlation coefficient and P value are indicated. **E–L,** Single-cell analysis of the PCa-Stem program using the integrated PCa epithelial scRNA-seq meta-atlas and cell-type annotations established by Zhao et al. (2024) (PMID: 39418984). Published cell-type annotations were retained for the present analyses. **(E)** UMAP visualization of epithelial cell populations, including basal, hillock, club, luminal, lineage plasticity-related cells (LPCs), and neuroendocrine (NE) cells. LPCs (red arrow) are indicated. **(F)** Dot plot showing expression of established epithelial lineage markers and PCa-Stem signature genes across the annotated epithelial populations. The 12 PCa-Stem signature genes are highlighted in red. Dot size indicates the percentage of cells expressing each gene, whereas color intensity represents scaled average expression. **(G)** UMAP visualization of PCa-Stem activity quantified by AUCell, showing preferential enrichment in LPCs. **(H)** Comparison of PCa-Stem AUCell scores across epithelial cell populations, demonstrating highest activity in LPCs and secondary enrichment in NE cells. **(I)** Distribution of PCa-Stem AUCell scores across epithelial cells. The automatically selected AUCell threshold was used to identify PCa-Stem⁺ cells; cells with AUC scores >0.05 were classified as PCa-Stem⁺. **(J)** Comparison of the percentage of PCa-Stem⁺ epithelial cells among normal prostate (N), adjacent benign tissue (Adj), Pri-PCa (Pri), and CRPC/metastatic disease (CRPC/Met). The wedge above indicates the Jonckheere–Terpstra (J-T) trend test across the ordered disease groups (*P* = 1 × 10⁻⁴), demonstrating a progressive increase in PCa-Stem⁺ epithelial cells with PCa progression. Pairwise statistical comparisons are indicated by brackets. **(K)** Percentage of PCa-Stem⁺ epithelial cells in individual samples across N, Adj, Pri, and CRPC/Met groups. Each bar represents an individual sample, ordered according to the percentage of PCa-Stem⁺ cells within each disease group. **(L)** Correlation between pseudobulk PCa-Stem scores and mRNAsi across individual samples. Spearman’s correlation coefficient and *P* value are indicated.

We next examined the cellular distribution of the PCa-Stem program using an integrated PCa epithelial scRNA-seq meta-atlas (Zhao 2024) [57]. Within the established epithelial lineage framework, most PCa-Stem signature genes were preferentially expressed in lineage plasticity-related epithelial cells or LPCs [57] and, to a lesser extent, NE cells (Fig. 7E-F), further linking the PCa-Stem program to epithelial states associated with lineage plasticity. Consistently, AUCell analysis localized the composite PCa-Stem activity predominantly to LPCs, with secondary enrichment in NE cells (Fig. 7G–H). AUCell analysis further identified a distinct PCa-Stem⁺ epithelial population (Fig. 7I), whose frequency progressively increased from normal prostate to Pri-PCa and advanced CRPC/metastatic disease (Fig. 7J–K). At the pseudobulk sample level, PCa-Stem scores correlated strongly with mRNAsi (Fig. 7L), further supporting the PCa-Stem signature as a ‘compact’ molecular representation of the broader global Stemness program.

To independently validate these observations, we analyzed the Cheng 2022 PCa scRNA-seq dataset [58] using cell-type annotations following the LPC classification framework by Zhao *et al.* (2024) [57]. PCa-Stem activity was, again, preferentially enriched in LPCs, with lower but detectable activity in NE cells (Fig. S10A–C). Consistent with the discovery meta-atlas, individual PCa-Stem signature genes were preferentially expressed in LPCs and NE cells (Fig. S10D).

Taken together, these spatial and single-cell transcriptomic analyses identify LPCs as the principal epithelial population harboring the PCa-Stem program and reveal the progressive expansion of PCa-Stem^+^ epithelial cells during PCa progression.

### Stemness-high PCa exhibit increased genomic alterations and distinct genomic features

To explore potential genomic differences between global Stemness-high and Stemness-low prostate tumors, we first compared genomic alteration burden using faction of genome altered (FGA) and tumor mutation burden (TMB) and found that mCRPC exhibited a greater FGA (Fig. 8A) and higher TMB (Fig. 8B) than Pri-PCa. Furthermore, within both Pri-PCa and mCRPC, the Stemness-high tumors displayed a significantly higher FGA (Fig. 8A) and TMB (Fig. 8B) than Stemness-low tumors, despite no differences in patient age between the groups (Fig. 8C). These findings support a strong association between oncogenic Stemness and genomic instability. We next examined the distribution of recurrent cancer-associated alterations in Stemness-defined subgroups across disease states (Fig. 8D–E). In treatment-naïve Pri-PCa, Stemness-high tumors were enriched for tumor-suppressor lesions (including *PTEN* and *RB1* deletions (DEL) and *TP53* mutations (MUT)), and alterations in known PCa drivers such as *SPOP* and *FOXA1* MUT and focal *MYC* amplification (AMP) (Fig. 8D). In contrast, the Stemness-high mCRPC exhibited markedly increased frequencies of therapy-selected drivers such as AR alterations (*AR* MUT, *AR* AMP; Fig. 8F). Furthermore, PI3K-pathway lesions, including *PIK3C2B* AMP (Fig. 8E), were enriched in Stemness-high mCRPC in contrast to their low prevalence in Pri-PCa. Several additional amplified loci—including *ELK4*, *RECQL4*, *SLC45A3*, and *ELOC*—were present in both disease stages but showed substantially higher frequencies in mCRPC (Fig. 8D-E). Notably, copy-number gain of *SLC45A3*, an androgen-regulated gene, was enriched in Stemness-high tumors and increased in frequency in mCRPC (Fig. 8E). These results support selective amplification of prostate lineage–associated loci under ARPI treatment and during progression to mCRPC.

**Figure 8.**
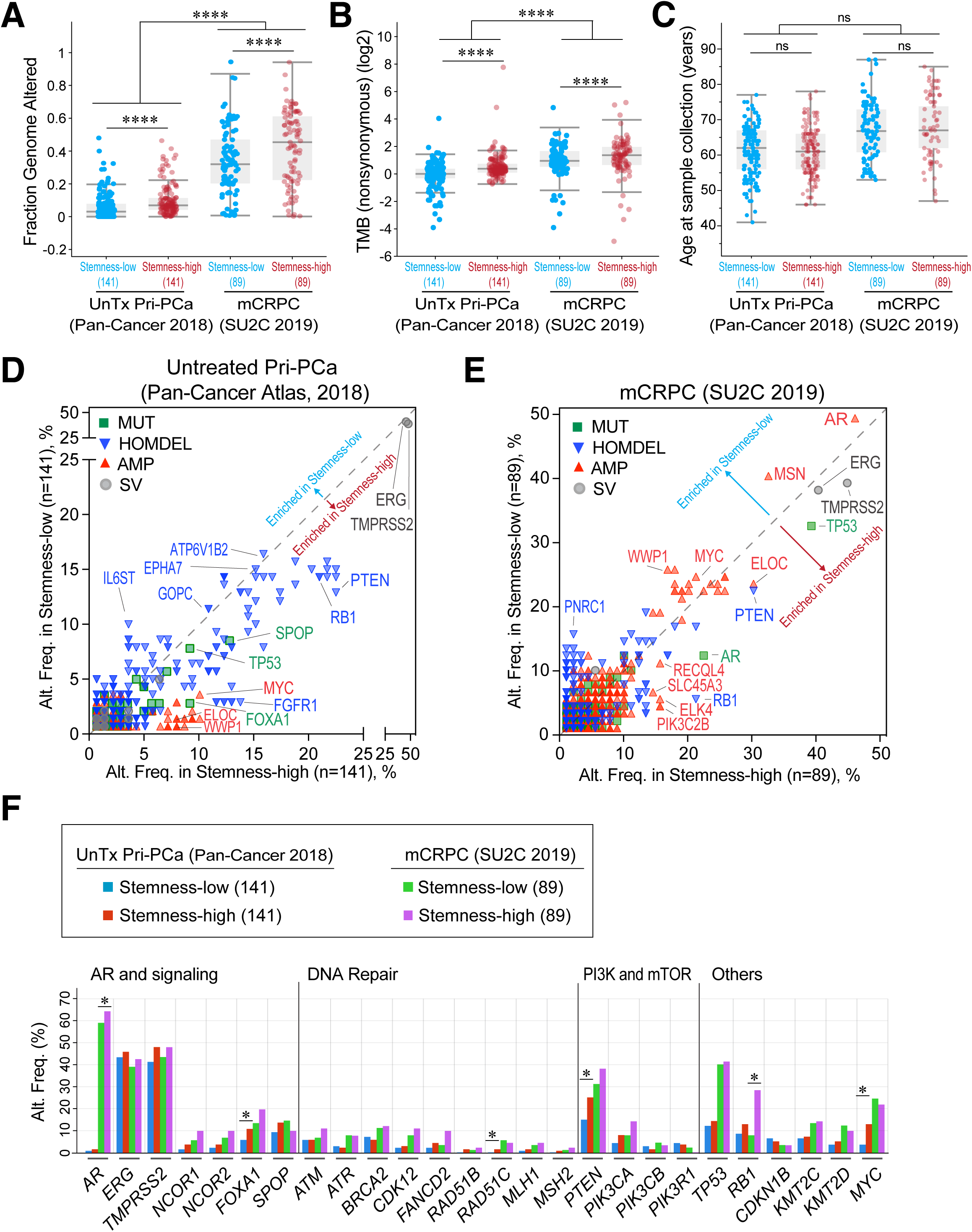
Stemness-high PCa is associated with increased genomic alteration burden. **A-C,** Box plots showing that Stemness-high subgroup in both Pri-PCa (n = 141) and mCRPC (n = 89) displays a higher fraction of altered genome (**A**) and higher TMB (**B**), but no age difference (**C**) compared to Stemness-low cases. Wilcoxon rank-sum test was used to determine significance (****, *P* < 0.0001; ns, not significant). Box plots represent the median, interquartile range (IQR), and whiskers extending to 1.5× the IQR. **D-F,** Comparisons of genomic alteration frequencies (Alt. Freq) in cancer-associated genes between Stemness-high and Stemness-low subgroups in TCGA Pri-PCa (**D**, **F**) and SU2C mCRPC (**E**, **F**) cohorts. Statistical significance was determined by two-sided Fisher’s Exact test (*, *P* < 0.05).

Globally, both Pri-PCa and mCRPC exhibited mutations in more than a dozen genes although most of these alterations were <10% (Fig. S11A; green symbols), consistent with reports that PCa is genomically heterogeneous with a broad spectrum of genomic alternations of low penetrance [87, 88]. On the other hand, homozygous deletion (HOMDEL) events, particularly, loss of *PTEN* and *RB1*, occurred frequently in both Pri-PCa and mCRPC (Fig. S11A; blue symbols) whereas AMP events in oncogenes such as *MYC* were more prevalent in mCRPC (Fig. S11A; red symbols). Nevertheless, *PTEN* and *RB1* HOMDEL represented among the most prevalent alterations in early-stage (GS6) PCa (Fig. S11B-D). Analysis of genetic alterations across the spectrum of PCa progression revealed that among the 3 tumor suppressors (*RB1, PTEN* and *TP53*), while *RB1* HOMDEL remained relatively constant at 13%, 6%, 10% and 10% in low-grade (GS6), high-grade (GS9/10), Pri-PCa/ADT and mCRPC, respectively, *PTEN* HOMDEL (9%, 23%, 30% and 26%) and *TP53* MUT (0%, 21%, 27% and 37%) continued to increase along this progression (Fig. S11B-D). Among the 3 oncogenic events analyzed, while high levels of *TMPRSS2* fusions (∼40%) remained unchanged from low-grade Pri-PCa to mCRPC, *MYC* (2%, 12%, 19% and 24%) and *AR* (0%, 1%, 6% and 49%) AMP continued to increase along the progression trajectory (Fig. S11B-C, E). In fact, prevalent *AR* MUT (11%) were observed only in the most aggressive and Stemness-highest mCRPC (Fig. S11B-C). Analyzing these major genomic alterations in association with PCa Stemness, we observed that *RB1* HOMDEL, *FOXA1* (together with *SPOP*, *IDH1*, and *CHD1*) MUT, and especially *MYC* AMP were significantly associated with increased Stemness in Pri-PCa (Fig. S11F) whereas in mCRPC, only *RB1* HOMDEL exhibited a statistically significant association with higher Stemness (Fig. S11G).

### MYC signaling functionally contributes to the proliferative Stemness program during PCa progression

The above studies (Fig. 8; Fig. S11) provided genomic correlates of PCa stemness. *In following studies, we explored 1) MYC signaling, 2) noncanonical (i.e., castration-induced) AR activity, and 3) tumor suppressor loss as potential drivers of persistently increasing Stemness during PCa progression*. We showed that in early PCa, increased c_AR-A may drive elevated Stemness (Fig. 1D-F); however, treatment-naïve Pri-PCa displayed continually increasing Stemness but decreasing c_AR-A (Fig. 2). As *AR* AMP and MUT are exclusively treatment-induced and there were virtually no *AR* genomic alterations in treatment-naïve Pri-PCa (Fig. S11B-C, E), what could be driving high Stemness during Pri-PCa progression? We found that Stemness-high Pri-PCa, compared to Stemness-low Pri-PCa, were enriched in MSigDB Hallmark MYC_TARGETS (Fig. 5E) and had more prevalent *MYC* AMP (Fig. 8D, F). High-grade (GS9/10) Pri-PCa also had more *MYC* AMP events than low-grade (GS6) Pri-PCa, and in fact, *MYC* AMP continued to increase along the PCa continuum (Fig. S11B-C, E) and was associated with increased Stemness in Pri-PCa (Fig. S11F). These observations led us to hypothesize that MYC may represent an important regulator of the Stemness program during both spontaneous Pri-PCa progression and mCRPC transition. To test this hypothesis, we quantified MYC signaling activity [61] (**MYC-sig**; See Methods and Table S2 for gene signature information) across the PCa progression spectrum. We observed that: 1) both *MYC* mRNA and MYC-sig increased in Pri-PCa compared to matched benign samples (Fig. 9A-B); 2) MYC-sig continued to go up during Pri-PCa progression (Fig. 9C); 3) unlike nADT-induced concomitant decline in both c_AR-A and Stemness (Fig. 1D-F; Fig. S4), nADT showed no discernible effect on MYC-sig (Fig. S12A-D); and 4) MYC-sig continued to increase across the disease spectrum with mCRPC exhibiting the highest MYC activity (Fig. 9D-E) and with MYC-sig positively correlated with the Stemness (Fig. 9F-H).

**Figure 9.**
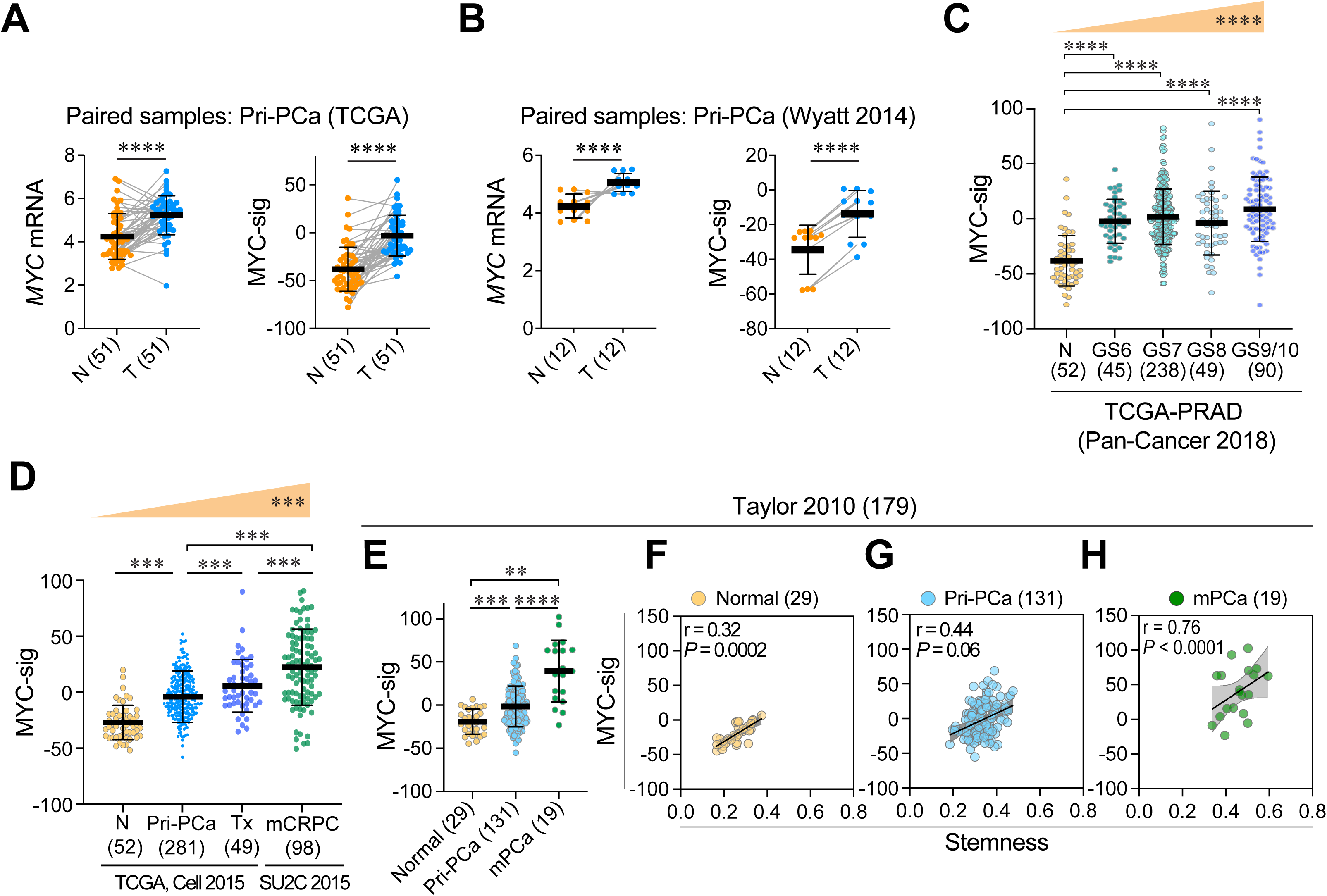
MYC activation contributes to increased Stemness during spontaneous PCa progression. **A-B,** Pairwise comparisons showing increased *MYC* mRNA levels and MYC signaling activity (MYC-sig) in primary tumors (T) versus matched adjacent benign tissues (N) in TCGA-PRAD cohort (n=51; **A**) and Wyatt (n=12; **B**) cohorts. Data is shown as mean ± SD; paired samples are linked by grey lines. Significance was determined using two-tailed paired Student’s *t*-test (****, *P* < 0.0001). **C-H,** MYC activity increases during PCa progression, peaking in mCRPC. In treatment-naïve Pri-PCa (**C**), MYC-sig is significantly increased in all-grade (GS) tumors as compared to tumor-adjacent benign tissues (N) with the most advanced GS9/10 tumors exhibiting the highest MYC-sig. In **D**, MYC-sig increases from normal to treatment-naïve Pri-PCa and further in advanced Pri-PCa treated with adjuvant ADT (Tx) in TCGA-PRAD, while mCRPC shows the highest MYC-sig in both SU2C 2015 (**D**) and Taylor 2010 (**E**) cohorts. Data is presented as mean ± SD. Statistical significance was assessed by one-way ANOVA with Tukey’s multiple-comparison test (**C-E**). J-T trend test was used in (C, D) to assess the significance of progressive increase. **, *P* < 0.01; ***, *P* < 0.001; ****, *P* < 0.0001; ns, not significant. **F-H,** Scatter plots showing positive correlations between MYC activity and Stemness in the Taylor 2010 cohort across different stages. Pearson’s r and P-values are indicated.

To further test whether MYC functionally promotes stemness, we first analyzed an independent RNA-seq dataset generated in LNCaP cells with MYC knockdown (KD) (GSE135942) [52]. MYC silencing markedly reduced MYC activity (Fig. S12E), confirming effective suppression of the MYC transcriptional output. Importantly, MYC-KD significantly reduced the global Stemness score (Fig. S12E). We next examined three independent pharmacological inhibition datasets (GSE178869, ref. 53; GSE135800 and GSE135876, ref. 54; GSE98069, ref. 55). Direct MYC inhibition using MYCi975 significantly reduced MYC activity as well as Stemness in AR-positive 22Rv1 CRPC cells (Fig. S12F) whereas both MYCi975 and MYCi361 similarly suppressed MYC activity and Stemness in AR-negative PC3 cells (Fig. S12G–H). Finally, treatment with the BET bromodomain inhibitor JQ1, an indirect suppressor of MYC transcription, reduced MYC activity, Stemness, *MYC* mRNA, and *KLK3* expression without significantly altering *AR* mRNA levels (Fig. S12I). Across multiple PCa models, the magnitude of Stemness reduction was highly correlated with suppression of MYC activity (Fig. S12I), supporting a close functional relationship between MYC signaling and the Stemness program.

Together, these genetic and pharmacological perturbation studies support that MYC functionally contributes to the proliferative Stemness program in diverse PCa models.

### Elevated castration-reprogrammed AR activity (cr_AR-A) and tumor suppressor-loss signatures are also associated with increasing Stemness during PCa progression

Having established MYC as a driver of Stemness during PCa progression, we next asked how therapeutic pressure may remodel AR signaling to sustain and amplify the stem-like programs. Androgen deprivation and AR-pathway inhibition remodel the AR cistrome and redirect AR toward non-canonical survival/plasticity programs [14–17]. To capture this shift, we derived the **cr_AR-A** by integrating AR and H3K27ac ChIP-seq data from Pomerantz *et al.* (GSE130408; [16]) with mRNA profiling data from Taylor *et al.* (GSE21032; [47]) (Fig. S13A; Table S2). We intersected genes that exhibited CRPC-enriched AR binding, enhancer activation, and transcriptional up-regulation to obtain a 63-gene set representing AR-regulated outputs preferentially active in mCRPC (Fig. S13A; Table S6).

As expected, cr_AR-A increased sharply in androgen-independent (AI) versus androgen-dependent (AD) LNCaP xenografts, whereas the canonical AR program (c_AR-A) declined (Fig. S13B). Genome-browser views confirmed that enhancer-level reprogramming [16] for *AR*, *UBE2C*, *CDK1*, *MELK*, *BIRC5*, and many other oncogenic and proliferation-associated genes displayed stronger AR and H3K27ac peaks in CRPC than in Pri-PCa (Fig. S13C; data not shown), while *CNN1* and *TNS1* showed the opposite pattern (Fig. S13D). To further benchmark the cr_AR-A, we compared it with the Wang “**cr_AR-Growth**” 8-gene set [14] curated from AR-bound, enhancer-validated and AI LNCaP-abl growth drivers (e.g., *UBE2C*, *CDK1*, *CDC20*) (Fig. S13E). Despite distinct derivations, the two signatures, with only one overlapped gene (i.e., *CDK1*), were highly concordant (r = 0.93, *P* < 0.0001; Fig. S13F–H) and both increased from normal/benign to Pri-PCa to mCRPC (Fig. 10A; S13I), underscoring a shared transcriptional logic linking therapy-reprogrammed AR signaling with proliferative Stemness. Together, these data reveal a state-dependent bifurcation of AR programs: the c_AR-A tracks differentiation in androgen-intact contexts whereas the cr_AR-A couples AR signaling to stem-like aggressiveness under therapeutic pressure, establishing cr_AR-A as a transcriptomic bridge between AR adaptation and PCa evolution.

**Figure 10.**
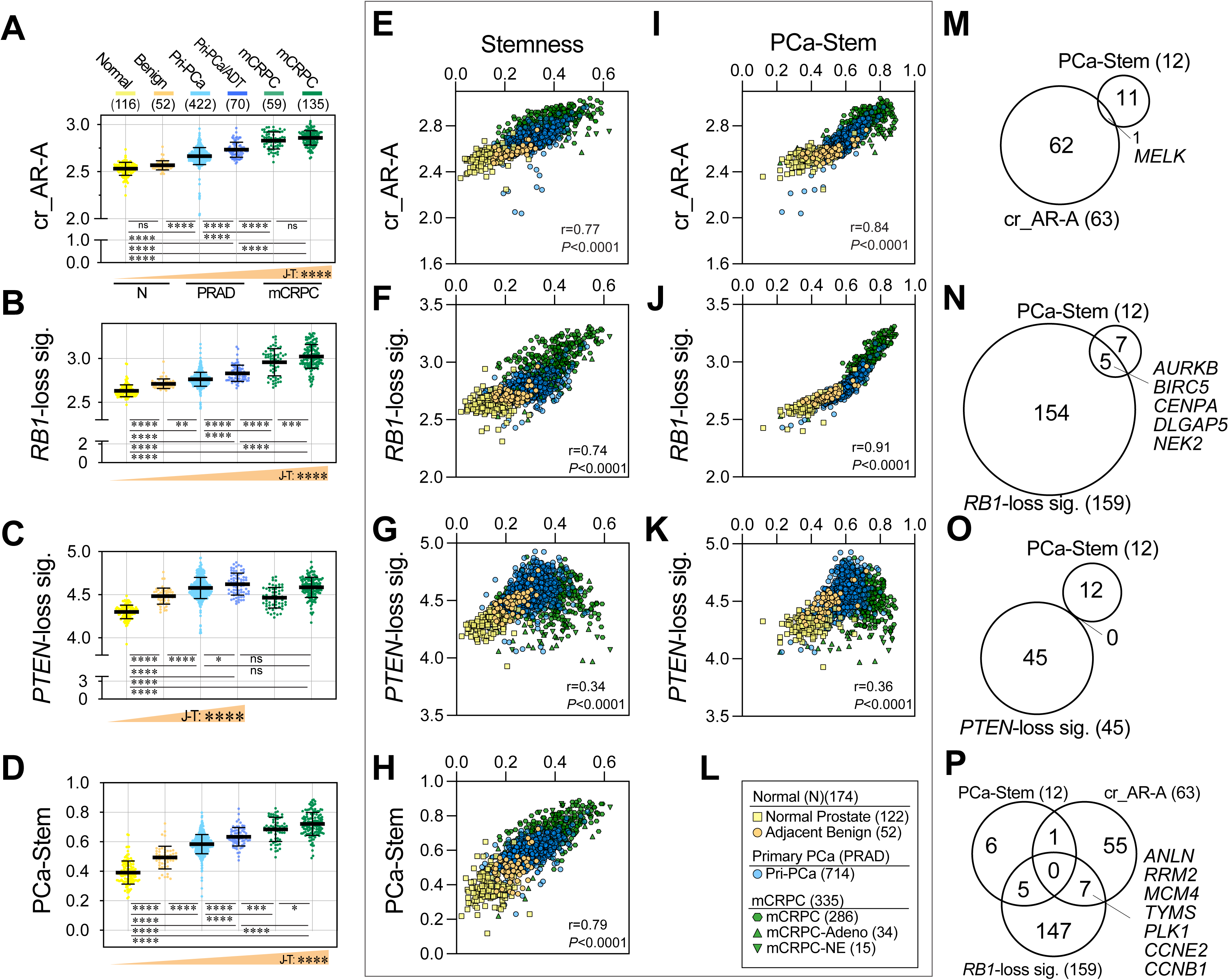
Castration-reprogrammed AR activity (cr_AR-A) and RB1-loss signature contribute to increased Stemness in mCRPC. **A–D,** The cr_AR-A and genomic *RB1*-loss signature increase progressively from normal and Pri-PCa to mCRPC. Shown are cr_AR-A (**A**) and *RB1*-loss (**B**), *PTEN*-loss (**C**) and PCa-Stem **(D)** signature scores across the disease spectrum using the Bolis integrated PCa cohort: normal prostates (GTEx, n=116), tumor-adjacent benign tissues (TCGA-PRAD, n=52), treatment-naïve Pri-PCa (TCGA-PRAD, n=422), aggressive Pri-PCa treated with adjuvant ADT (Pri-PCa/ADT) (n=70), and two mCRPC cohorts including GSE126078 mCRPC (n=59) and SU2C 2015 (n=135). Sample sizes are indicated in parentheses. Shown at the bottom are Jonckheere–Terpstra (J-T) trend-test results (wedges). **E–H,** Scatter plots illustrating correlations between cr_AR-A (**E**), *RB1*-loss (**F**), *PTEN*-loss (**G**), PCa-Stem (**H**) scores with Stemness across Bolis integrated cohort (total n=1,223; including normal (n=174), Pri-PCa (n=714), and mCRPC (n=335)). Pearson’s correlation coefficients (*r*) and *P* values are shown. Details of symbol colors and shapes are shown in (**L**). **I–L,** Inter-signature relationships among cr_AR-A, *RB1*-loss, *PTEN*-loss, and PCa-Stem signatures in the Bolis integrated cohort (total n=1,223, normal n=174, Pri-PCa n=714, mCRPC n=335). Correlation plots demonstrate strong positive correlations between cr_AR-A and PCa-Stem (**I**), and between *RB1*-loss and PCa-Stem (**J**), and a moderate but significant correlation between *PTEN*-loss and PCa-Stem (**K**). Details of symbol colors and shapes are shown in (**L**). **M-P,** Venn diagrams showing overlapping genes among cr_AR-A (**M, P**), *RB1*-loss (**N, P**), *PTEN*-loss (**O**), and PCa-Stem signatures, highlighting shared components (e.g., *AURKB, BIRC5, CENPA, PLK1, ANLN)* involved mitotic regulation and the self-renewing proliferation typical of stem-like tumor cells. All dot plots data are shown as mean ± SD. Statistical significance among groups was determined using one-way ANOVA followed by Tukey’s multiple-comparison test, and J-T trend tests were used to assess significance across disease states. Pearson correlation coefficients (*r*) were calculated for all pairwise associations. *ns*, not significant; *, *P* < 0.05; **, *P* < 0.01; ***, *P* < 0.001; ****, *P* < 0.0001. See Table S4 for full statistical summaries and data sources.

Finally, we examined whether transcriptomic signatures associated with tumor suppressor losses may also reinforce therapy-driven Stemness. Prior genomic analyses (Fig. 8; Fig. S11) showed that, in Pri-PCa, *RB1* DEL, *FOXA1* MUT, and especially *MYC* AMP associate with elevated Stemness whereas *PTEN* loss is prevalent early and persists into advanced disease and that in mCRPC, *RB1* DEL significantly correlated with high Stemness. To further assess these trends transcriptionally, we quantified the *RB1*-loss [62] and *PTEN*-loss [63] gene signatures, respectively, and found that the ***RB1*-loss signature** increased progressively from benign to Pri-PCa to mCRPC (Fig. 10B) whereas the ***PTEN*-loss signature** rose sharply from normal prostate to Pri-PCa but plateaued thereafter, peaking in aggressive Pri-PCa/ADT and remaining high in mCRPC (Fig. 10C). These analyses suggest that *PTEN* loss might function as an early and persistent driver of spontaneous progression whereas *RB1* loss represents a more prominent contributor to Stemness and proliferative reprogramming in therapy-resistant disease.

### Convergence of cr_AR-A, *RB1*-loss, and MYC signaling with global Stemness and PCa-Stem signature scores

Building on the above analyses showing progressive increase in cr_AR-A (Fig. 10A) and distinct dynamics of *RB1*-and *PTEN*-loss signatures (Fig. 10B-C), we next examined how these programs relate quantitatively to the general Stemness score and PCa-Stem signature (Fig. 10D-P; Fig. S13J-P). Consistent with its progressive rise across disease states, the PCa-Stem score increased from normal prostate and Pri-PCa to mCRPC (Fig. 10D), broadly tracking the trajectory of cr_AR-A and *RB1*-loss (Fig. 10A-B). Across the Bolis integrated cohort (n = 1,223), the cr_AR-A correlated strongly with both general Stemness (Fig. 10E; see Fig. 10L for symbol definitions) and the PCa-Stem (Fig. 10I). In parallel, the LNCaP-abl-derived cr_AR-Growth_Wang signature also correlated with the two Stemness indices (Fig. S13J-K). Similarly, the *RB1*-loss signature correlated with the global Stemness (r = 0.74, *P* < 0.0001; Fig. 10F) and even more strongly with PCa-Stem (r = 0.91, *P* < 0.0001; Fig. 10J) whereas the *PTEN*-loss signature exhibited a more modest association with the two Stemness indices (Fig. 10G, K).

We next integrated MYC signaling into this framework. Consistent with our earlier MYC analyses (Fig. 9), MYC-sig increased along the PCa evolution continuum in the Bolis integrated PCa cohort (Fig. S13M) and correlated strongly with both the global Stemness (Fig. S13N) and PCa-Stem (Fig. S13O). Of interest, despite this strong transcriptional convergence, the MYC-sig and PCa-Stem showed no direct gene overlap (Fig. S13P), indicative of their convergence at the level of coordinated downstream proliferative programs rather than shared gene membership.

Across the same cohort, PCa-Stem displayed the strongest positive correlation with the general Stemness (Fig. 10H), consistent with the PCa-Stem being derived from Stemness-stratified DEGs (Fig. 6A-B). Gene-overlap analyses further supported convergence among these programs. The PCa-Stem shared *MELK* with the 63-gene cr_AR-A signature (Fig. 10M) and shared *UBE2C* with the 8-gene cr_AR-Growth_Wang (Fig. S13L). The PCa-Stem also showed broader overlap with the *RB1*-loss signature (Fig. 10N), including *AURKB*, *BIRC5*, *CENPA*, *DLGAP5*, and *NEK2*, genes largely involved in cell-cycle progression and mitotic regulation. In contrast, no direct gene overlap was observed between *PTEN*-loss and PCa-Stem (Fig. 10O). When signatures were integrated (Fig. 10P), the proliferative core was most evident between cr_AR-A and the *RB1*-loss signature (i.e., *ANLN*, *RRM2*, *MCM4*, *TYMS*, *PLK1*, *CCNE2*, and *CCNB1*), supporting convergent selection for cell-cycle machinery characteristic of stem-like, self-renewing tumor cells.

## Discussion

The current project was initiated to quantitatively co-analyze the global changes of and interrelationship between cancer stemness and canonical AR signaling activity during PCa evolution. By employing gene expression-based general Stemness and c_AR-A signature scores and through developing a new 12-gene PCa-Stem signature, we show that while differentiation-associated c_AR-A steadily declines in the PCa continuum, progressively increasing Stemness quantitatively tracks PCa aggressiveness, plasticity and progression, and represents a poor prognosticator for patient outcomes. Stemness-high PCa possess unique genomic features including elevated TMBs and are transcriptionally enriched in aggressive molecular subtypes such as PAM50-LumB [80–82] and PCS1 [84, 85]. Our data further suggest that elevated Stemness in early PCa is causally associated with increased c_AR-A while castration-reprogrammed AR-A (cr_AR-A), MYC activation and tumor suppressor alterations (especially *RB1* loss) likely cooperate to propagate the pervasively increasing Stemness observed in mCRPC (where c_AR-A is suppressed).

Early studies reported that AR-regulated genes decline in high-grade Pri-PCa [89, 90] and inhibition of AR signaling promotes stem-like phenotypes [8, 51, 91, 92]. However, global pro-differentiation AR activity and oncogenic stemness have not been systematically and quantitatively co-analyzed across the PCa continuum. By integrating 33 transcriptomic datasets encompassing normal prostate, tumor-adjacent benign tissues, treatment-naïve Pri-PCa, nADT-treated tumors, (m)CRPC subtypes and model systems, we demonstrate that PCa progression is accompanied by a continual increase in Stemness and progressive decline in canonical c_AR-A, with the two inversely correlated (Fig. S14). Importantly, both the global Stemness and PCa-Stem signature prognosticate poor patient survival, establishing Stemness as a quantifiable and clinically relevant indicator of PCa progression. To our knowledge, this is the first comprehensive study to simultaneously quantify the oncogenic stemness and c_AR-A and to investigate the potential drivers of cancer stemness across the PCa continuum. Our pan-cohort approach enables signature validation in rare archival biopsy samples and cross-platform integration — a translational advantage not addressed by other studies.

The normal prostatic epithelium consists of AR^+^ secretory luminal cells and AR^-^basal cells. We find that NHP luminal cells have higher c_AR-A whereas basal cells display higher Stemness, consistent with biological studies from our lab [21] and others [19, 20, 76, 77] showing stem/progenitor cell properties enriched within the basal cell compartment. Compared to adjacent benign tissue, Pri-PCa exhibit elevated Stemness, which is associated with and likely driven by increased c_AR-A. Early genomic alterations including *PTEN* and *RB1* loss in GS6 tumors (9% and 13%, respectively; Fig. S11D) may further contribute to this initial rise in Stemness. Strikingly, tumor-adjacent benign tissues, the so-called ‘normal’ tissues used in most comparative studies, displayed higher Stemness than the true normal prostate obtained from cancer-free organ donors in GTEx, indicating that the commonly used “benign” controls may already exhibit transcriptional shifts toward cancer-associated programs.

Our studies suggest that the drivers of increased Stemness differ between spontaneous Pri-PCa progression and mCRPC transition. In treatment-naïve Pri-PCa, where tumor progression is characterized by a gradual decline in c_AR-A while Stemness shows a continual elevation (Fig. S14), increased Stemness may likely be driven by oncogenic events such as *MYC* amplification and activation, with additional contributions from early loss of tumor-suppressors such as *PTEN* and *RB1*. On the other hand, in mCRPC, which develops under sustained therapeutic pressure from long-term ADT and ARPI treatment, the c_AR-A is further suppressed while Stemness reaches its highest levels. This paradoxical sharp increase in Stemness despite reduced c_AR-A not only correlates tightly with *RB1* loss and MYC activation but also coincides with emergence of the cr_AR-A, which is enriched for cell-cycle and proliferation-associated genes (Fig. S14). Thus, the high Stemness in mCRPC is reinforced by AR functional reprogramming that cooperates with genomic alterations and MYC activity to sustain aggressive, proliferative and therapy-resistant stem-like states. Regardless, our analyses further strengthen the functional connection between MYC signaling and Stemness. Beyond the strong transcriptomic association observed across clinical cohorts, the genetic *MYC* knockdown and pharmacological MYC inhibition consistently reduced the Stemness across AR-negative PCa and CRPC models. These findings support MYC activity as an important regulator of the proliferative Stemness program in PCa and suggest that targeting MYC-associated transcriptional programs may represent a therapeutic strategy for Stemness-high PCa.

What biological correlates do our Stemness metrics measure? ***First***, it measures transcriptional features and relative abundance of stem cells and CSCs. Therefore, the NHP basal/stem cells display higher Stemness than luminal cells (consistent with literature [19, 21, 76, 77]); ARPI treatment, while shutting down the c_AR-A, enriches CSCs [7, 51, 91, 92] and upregulates Stemness (this study); and PCa cell lines, xenografts and PDX, known to be clonally derived from cancer stem/progenitor cells, demonstrate the highest Stemness. ***Second***, it measures cellular and lineage plasticity [8] induced by treatment and genetic alterations. Thus, mCRPC, which harbor more abundant treatment-reprogrammed AR^-/lo^ cells and PCa cells with stem-like features or in an AR-indifferent state [8, 12, 13, 23, 46, 60, 83, 86], manifests the highest Stemness. Our analysis further substantiates the aggressive nature of (AR^-/lo^) CRPC-NEPC and DNPC compared to ARPC, as evidenced by their elevated Stemness scores (Fig. S5B). Importantly, our single-cell analyses provide cellular-resolution support for this association by localizing the PCa-Stem program predominantly to lineage plasticity-related epithelial cells [57], with secondary enrichment in NE cells (Fig. 7; Fig. S10). The abundance of PCa-Stem⁺ epithelial cells progressively increased during disease progression, while spatial transcriptomics demonstrated regional concordance between the PCa-Stem and global Stemness (Fig. 7). Together, these findings support a shift in the phenotypic landscape of advanced PCa toward more plastic, stem-like and aggressive epithelial states. Likewise, aggressive DKO and TKO murine prostate tumors have undergone lineage transformation and exhibit higher Stemness than indolent SKO tumors. ***Third***, both the global Stemness and PCa-Stem metrics likely report the coordinated proliferative Stemness program and high proliferative robustness characteristic of advanced PCa, mCRPC and NEPC, as supported by our network, functional, and single-cell analyses. Indeed, 11 of the 12 PCa-Stem signature genes are well-characterized regulators of G2/M transition, mitosis, cell division and cell proliferation. ***Finally***, the Stemness metrics quantitatively measure PCa aggressiveness as high Stemness correlates not only with more advanced and aggressive stages of PCa but also with aggressive molecular subtypes and poor patient survival.

It’s worth reiterating that the newly derived 12-gene PCa-Stem signature further refines the concept of proliferative Stemness by providing a compact PCa-specific molecular representation. Although derived solely by differential expression between Stemness-high and Stemness-low tumors shared between Pri-PCa and mCRPC, 11 of the 12 PCa-Stem signature genes formed a highly interconnected protein-protein interaction network and converged on mitotic chromosome segregation, spindle assembly, and cell-cycle checkpoint regulation. Upstream regulatory analyses further connected the PCa-Stem program with established drivers of lineage plasticity and therapy resistance, including RB1, TP53, MYC, FOXM1, E2F, and YAP1. Coupled with our prospective studies with knockdown of 3 representative PCa-Stem signature genes (i.e., *HMMR*, *AURKB*, and *PBK*), these analyses provide a biologically coherent framework linking proliferative reprogramming with stemness during PCa progression.

A key implication of our study is that stemness in PCa represents a dynamic cell-state continuum rather than a fixed CSC population. Studies in normal adult tissues have demonstrated that regenerative capacity is maintained through the coexistence of quiescent reserve stem cells and fast-cycling active stem cells, with cells reversibly transitioning between these states in response to stress or injury [93]. Applying this framework to PCa, our data suggest that both the general Stemness Index and PCa-Stem signature preferentially reflect a fast-cycling, proliferation-enriched stem-like state. This framework explains why Stemness progressively increases despite declining c_AR-A associated with cellular maturation: therapeutic pressure and accumulating genomic alterations select for proliferating and plastic cell states rather than a static CSC compartment. Our functional validation provides support for this proliferation-driven cancer stemness -depletion of mitotic kinases *AURKB* and *PBK* suppressed invasion, clonogenic growth, and 3D spheroid viability (Fig. 6F-H; Fig. S9E-H). Notably, this proliferative Stemness state mirrors transcriptional programs captured by cr_AR-A and cr_AR-growth signatures, both enriched for cell-cycle genes and strongly correlated with global Stemness and PCa-Stem scores. Together, these data support convergence on a fast-cycling, therapy-adapted stem-like state in aggressive PCa, consistent with recently reported convergence of CSC states across tumor types [94].

Of clinical significance, both the general Stemness Index and the 12-gene PCa-Stem signature correlate with poor patient survival, suggesting the possibility of developing the Stemness metrics into a predictive biomarker to distinguish aggressive from indolent primary tumors and a prognostic indicator of unfavorable clinical outcomes in mCRPC patients. Notably, 11 of the 12 PCa-Stem signature genes correlate with PCa progression (Fig. S7B), supporting their individual associations with the increased Stemness in advanced Pri-PCa and mCRPC. Analysis using data from The Human Protein Atlas (www.proteinatlas.org) reveals that these 11 genes are also recognized as unfavorable prognostic markers in PCa and several other cancers (Liu and Tang, unpublished), raising the potential utility of the PCa-Stem signature as a prognostic marker not only for PCa but also for other malignancies. Coupled with earlier studies linking increased Stemness with aggressive breast and brain cancer subtypes [6], our findings herein underscore the increased Stemness as a ‘universal’ and fundamental characteristic of cancer progression and aggressiveness. Beyond its ability to track tumor dedifferentiation and lineage plasticity across the PCa continuum, the PCa-Stem signature demonstrated improved disease-focused classification compared with the pan-cancer Stemness Index (Fig. S7D-E). This does not diminish the biological value of the general Stemness Index; rather, PCa-Stem complements it by providing increased PCa–specific sensitivity.

In recent years, several studies [e.g., 36, 57, 95-97] have proposed stemness-based PCa classifications or prognostic models using either single-cell RNA-seq, bulk RNA-seq or integrative analyses. These studies reinforce the biological relevance of stemness and highlight oncogenic pathways such as MYC activation, *RB1*/*PTEN* loss, and lineage plasticity in PCa progression. Our findings extend these observations in several important aspects. First, we establish quantitative Stemness metrics and validate them across a large pan-cohort framework spanning the entire PCa continuum. Second, by integrating spatial transcriptomics with two independent scRNA-seq datasets, we localize the PCa-Stem program to lineage plasticity-related epithelial cells and demonstrate progressive expansion of Stemness-associated epithelial cells during PCa progression. Finally, the compact 12-gene PCa-Stem signature provides a biologically interpretable and experimentally tractable molecular representation of global oncogenic Stemness while retaining strong prognostic performance. Of note, our 12-gene PCa-Stem signature, derived from bulk transcriptomic stratification, shows strong biological overlap with the lineage plasticity signature (LPSig) reported by Zhao *et al*. using scRNA-seq [57]. Our independent single-cell analyses further localize the PCa-Stem program to the same lineage plasticity-related epithelial cell population, providing orthogonal support for the biological relevance of this proliferative Stemness program. scRNA-seq analysis showed that individual PCa-Stem signature genes exhibited expression patterns largely consistent with the composite PCa-Stem program (Fig. S10). Among the shared genes, *HMMR* emerges as a critical node of convergence as it has been shown that *HMMR* promotes NE differentiation and aggressive behavior in CRPC models [57]. The presence of *HMMR*, along with *DLGAP5*, *AURKB*, and *BIRC5* in our PCa-Stem signature, highlights the translational potential of this gene set in targeting NEPC and other plasticity-driven CRPC subtypes. Supporting this, Zheng *et al*. used multi-omics integration and machine learning to define three stemness-based PCa subtypes and constructed a nine-gene prognostic signature [95] that also included *HMMR* and *DLGAP5*. *KLK12*, another PCa-Stem signature gene, was recently identified as a unique marker for SOX9^High^ stem-like luminal epithelial cells enriched following androgen deprivation [98]. Interestingly, we observed *KLK12* to be significantly enriched in NE cells (Fig. S10D). Additionally, our findings that *MYC* amplification, *RB1* deletion, and *PTEN* loss are enriched in Stemness-high tumors are consistent with the previously reported chromatin accessibility (ATAC-seq) patterns [86] and multi-omics spatial data [98]. It is worth noting that although our PCa-Stem signature was initially designed for risk stratification in mCRPC, the analytical framework is inherently flexible as the same platform can be extended to derive subtype-specific signatures.

We recognize that the global Stemness algorithm utilized in our studies was not developed specifically for PCa and therefore, its sensitivity to distinguish subtle intra-group differences remains to be fully validated. Furthermore, cellular localization and biological relevance of the PCa-Stem signature initially derived from bulk transcriptomic data, although independently validated using spatial transcriptomics together with two scRNA-seq datasets, should be further investigated in future studies by incorporating additional longitudinal single-cell datasets. In addition, incorporating CytoTRACE-based analyses [99, 100], which infer cellular developmental potential and differentiation state, may provide a complementary measure of developmental Stemness and help distinguish developmental potential from the progression-associated cancer Stemness captured by the current transcriptomic framework. Cancer-specific genomic or cell-free DNA biomarkers [101] may further improve the clinical applicability of Stemness assessment. Finally, although our network analyses nominate coordinated regulatory modules underlying the PCa-Stem program, the predicted interactions remain hypothesis-generating and warrant future mechanistic validation.

In conclusion, we establish transcriptome-based signature scores to assess pro-differentiation AR activity, MYC signaling, and oncogenic stemness across the PCa continuum. Stemness progressively increases during spontaneous PCa progression and becomes much heightened in mCRPC through AR reprogramming, *RB1*-loss associated transcriptional programs, and MYC activation. These findings provide a quantitative framework linking PCa progression, oncogenic dedifferentiation, proliferative reprogramming, lineage plasticity and therapy resistance, and identify stemness-associated transcriptional programs as potential clinical vulnerabilities. Together, the signature scores reported herein have the potential to serve as clinical classifiers for identifying aggressive disease and patients who may benefit from stemness-targeting therapies.

## Data Availability

Data Availability: All data analyzed in this study were obtained from publicly available repositories, including GEO (RRID:SCR_005012), cBioPortal (RRID:SCR_014555), the NCI GDC Portal, the UCSC Xena Functional Genomics Portal (RRID:SCR_018938), EGA (RRID:SCR_004944), ENA (RRID:SCR_006515), and Zenodo (RRID:SCR_004129), as detailed in Table S1 and described in the Materials and Methods section. No new sequencing or genomic data were generated in this study.

## Code availability

All custom R scripts used for quantification of Stemness Score (i.e., mRNAsi), PCa-Stem signature ssGSEA scoring, Jonckheere–Terpstra trend testing, multivariable Cox modeling, and regression analysis are available on GitHub: https://github.com/DaltonLiu2026/ProstateCancer-Stemness-Scripts. A permanently archived, citable version of this repository has been deposited in Zenodo: https://zenodo.org/records/15620990. The GitHub and Zenodo repository include all scripts, example input/output files, a README, dependency list, and a code-reproducibility overview.

## Acknowledgements

We gratefully acknowledge Dr. X. Qiu for kindly providing supporting data for AR cistrome, H3K27ac profiles and transcriptomic data in prostate cancer tissue samples (Pomerantz *Nat Genet* 2020); the cloud computing resources provided by the Center for Computational Research (CCR) at the University at Buffalo (http://hdl.handle.net/10477/79221); Mr. W. Tian and Dr. L. Yan for assistance in survival analysis; Drs. J. Yong and H. He for bioinformatics assistance and all other Tang lab members for helpful discussions and suggestions. We apologize to the colleagues whose work was not cited due to space constraint.

## Funding

Work in Tang lab was supported, in part, by grants from the U.S. National Institutes of Health (NIH) National Cancer Institute (NCI) R01CA237027, R01CA240290 and 2R01CA240290-06A1, grants from the U.S. Department of Defense (DOD) PC220137 and PC220273, Roswell Park Comprehensive Cancer Center and the NCI Center grant P30CA016056, Roswell Park Alliance Foundation (RPAF) and the George Decker Endowment fund. Work in Goodrich lab was support by NIH/NCI R01CA234162 and R01CA230913. Work in the labs of Goodrich, Tang and Chatta was further supported by the Prostate Cancer Foundation (PCF) Challenge Award 2022CHAL3788 (PI: Goodrich). S. Liu and J. Wang were supported by NIH/NCI U24CA232979 and U24CA274159. Work in Alumkal lab was supported by NIH/NCI R01CA251245, PCF Challenge Award, National Comprehensive Cancer Network (NCCN)/Astellas Pharma Global Development Award, Michigan Prostate SPORE (NCI P50 CA186786), and Joint Institute for Translational and Clinical Research.

## Authors’ Contributions

**X. Liu**: Conceptualization, data curation, formal analysis, statistical analysis, validation, visualization, investigation, writing–original draft, writing–review and editing, equal contribution as co-first author and co-senior author. **A. Jamroze**: Data curation, investigation, writing–review and editing, equal contribution as co-first author. **E. Cortes**: Data curation, formal analysis, statistical analysis, validation, investigation, writing–original draft, writing–review and editing, equal contribution as co-first author. **Y. Ji**: Study conception and design, data curation, validation, visualization. **K. Zhao**: Data curation, formal analysis, validation, visualization. **J. Ho**: Data curation, formal analysis, validation, visualization. **Y. Liu**: Data curation, formal analysis, validation, visualization. **E. Davicioni**: Data curation, investigation, writing–review and editing. **S. Wu**: Data curation and formal analysis. **W. Li**: Data curation and formal analysis. **F. Zhao**: Data curation and formal analysis. **Z. Xia**: Data curation and formal analysis. **F. Feng**: Writing–review and editing. **J. Alumkal**: Writing–review and editing. **D. Spratt**: Writing–review and editing. **C. Sweeney**: Writing– review and editing. **H. Yu**: Data curation, statistical analysis, writing–review and editing. **Q. Hu**: Data curation, statistical analysis, investigation. **C. Zou**: Data curation, investigation, writing–review and editing. **D. Zhang**: Data curation, investigation, writing–review and editing. **K. Lin**: Data curation and formal analysis. **Y. Lu**: Data curation and formal analysis. **G. Chatta**: Data curation, investigation, writing–review and editing. **K. Nastiuk**: Data curation, investigation, writing–review and editing. **D. Goodrich**: Obtained main funding for this project, data curation, writing–review and editing. **K. Rycaj**: Investigation, Data curation, visualization, writing–review and editing. **J. Kirk**: Data curation, investigation, writing–original draft, writing–review and editing. **I. Puzanov**: Writing–review and editing. **S. Liu**: Study conception and design, supervision, writing–review and editing, funding acquisition, equal contribution as co-senior author. **J. Wang**: Study conception and design, supervision, writing–review and editing, funding acquisition, equal contribution as co-senior author. **D. Tang**: Conceptualization, supervision, writing–original draft, writing– review, editing and finalization, funding acquisition, equal contribution as co-senior author, the lead corresponding author.

## Ethics declarations

### Ethics approval and consent to participate

Not applicable.

## Consent for Publication

Not applicable.

## Competing interests

All authors declare no competing interests relative to this work.

## Notes

### Competing Interest Statement

The authors have declared no competing interest.

### Summary of Updates

This revised version substantially expands the biological, functional, and single-cell characterization of prostate cancer Stemness and the 12-gene PCa-Stem signature. The manuscript now contains 10 main Figures, 14 Supplementary Figures, and 6 Supplementary Tables, including 2 new main Figures (Figs. 6 and 7) and 2 new Supplementary Figures (Figs. S8 and S10). Authorship was updated by moving Dr. A. Jamroze to the second co-contributing author position and adding 4 new authors (Drs. F. Zhao, S. Wu, W. Li, and Z. Xia) for their contributions to the new single-cell RNA-seq, spatial transcriptomic, and related bioinformatic analyses. The title was also slightly modified to use "cancer stemness" to more precisely define the Stemness measured in this study. Major scientific additions include expanded network and functional analyses linking the PCa-Stem signature to proliferative and lineage-plasticity programs; additional functional validation of representative PCa-Stem genes (AURKB, HMMR, and PBK), including AggreWell-based prostatosphere assays; and new spatial and single-cell transcriptomic analyses localizing the PCa-Stem program predominantly to lineage plasticity-related epithelial cells and demonstrating its progressive expansion during PCa progression. We also expanded MYC perturbation analyses and further examined relationships among Stemness, AR signaling reprogramming, tumor-suppressor alterations, genomic alteration burden, and AR-indifferent disease states.

