## Supplementary Figure S1 to Figure S14 for "Increasing Cancer Stemness Drives Prostate Cancer Progression, Plasticity, Therapy Resistance and Poor Patient Survival"

---

**Supplementary Table S1.** Summary of datasets used in this study.

**Supplementary Table S2.** Summary of gene signatures used in this study.

**Supplementary Table S3.** DEGs between Stemness-high and Stemness-low subgroups in Pri-PCa (TCGAPRAD) and mCRPC (SU2C 2019) cohort (*six sheets).

**Supplementary Table S4.** Summary of statistical significance of differences using Tukey's multiple comparisons test (*nine sheets).

**Supplementary Table S5.** Summary of gene signatures and gene sets used in Gene Set Enrichment Analysis (GSEA).

**Supplementary Table S6.** CRPC-enriched AR-binding, enhancer-marked, and transcriptionally up-regulated gene sets used for cr_AR-A signature derivation (*six sheets).

**Related to Fig. 1**

**Supplementary Fig. S1.** Schematic illustration of bioinformatic datasets and signature score pipelines used for comparisons across the spectrum of PCa evolution.

**Supplementary Fig. S2.** Transcriptome-based signature scores for canonical (pro-differentiation) AR signaling activity (c_AR-A).

**Supplementary Fig. S3.** Prostate luminal cell compartment harbors higher c_AR-A but lower Stemness as compared to prostate basal cell compartment.

**Supplementary Fig. S4.** nADT decreases both c_AR-A and Stemness.

**Related to Fig. 3**

**Supplementary Fig. S5.** Increased Stemness represents a 'universal' feature of PCa progression and aggressiveness.

**Related to Fig. 4**

**Supplementary Fig. S6.** High Stemness correlates with poor patient survival.

**Related to Fig. 6**

**Supplementary Fig. S7.** PCa-Stem signature gene expression correlates with PCa progression.

**Supplementary Fig. S8.** Biological network organization and functional annotation of the 12-gene PCa-Stem signature.

**Supplementary Fig. S9.** Functional and molecular characterization of representative PCa-Stem signature genes *AURKB*, *HMMR*, and *PBK* in PCa models.

**Related to Fig. 7**

**Supplementary Fig. S10.** Single-cell expression patterns of the 12 PCa-Stem signature genes in the Cheng 2022 PCa scRNA-seq dataset.

**Related to Fig. 8**

**Supplementary Fig. S11.** Unique genomic features of Stemness-high PCa.

**Related to Fig. 9**

**Supplementary Fig. S12.** nADT does not alter MYC activity whereas genetic and pharmacological MYC inhibition suppresses MYC activity and Stemness in diverse PCa models

**Related to Fig. 10**

**Supplementary Fig. S13.** Integrated multi-omic derivation and cross-validation of the castration-reprogrammed AR activity (cr_AR-A) signature.

**Supplementary Fig. S14**. Schematic summary of cancer Stemness changes across stages of PCa progression. See Text and figure legend.

Fig. S1

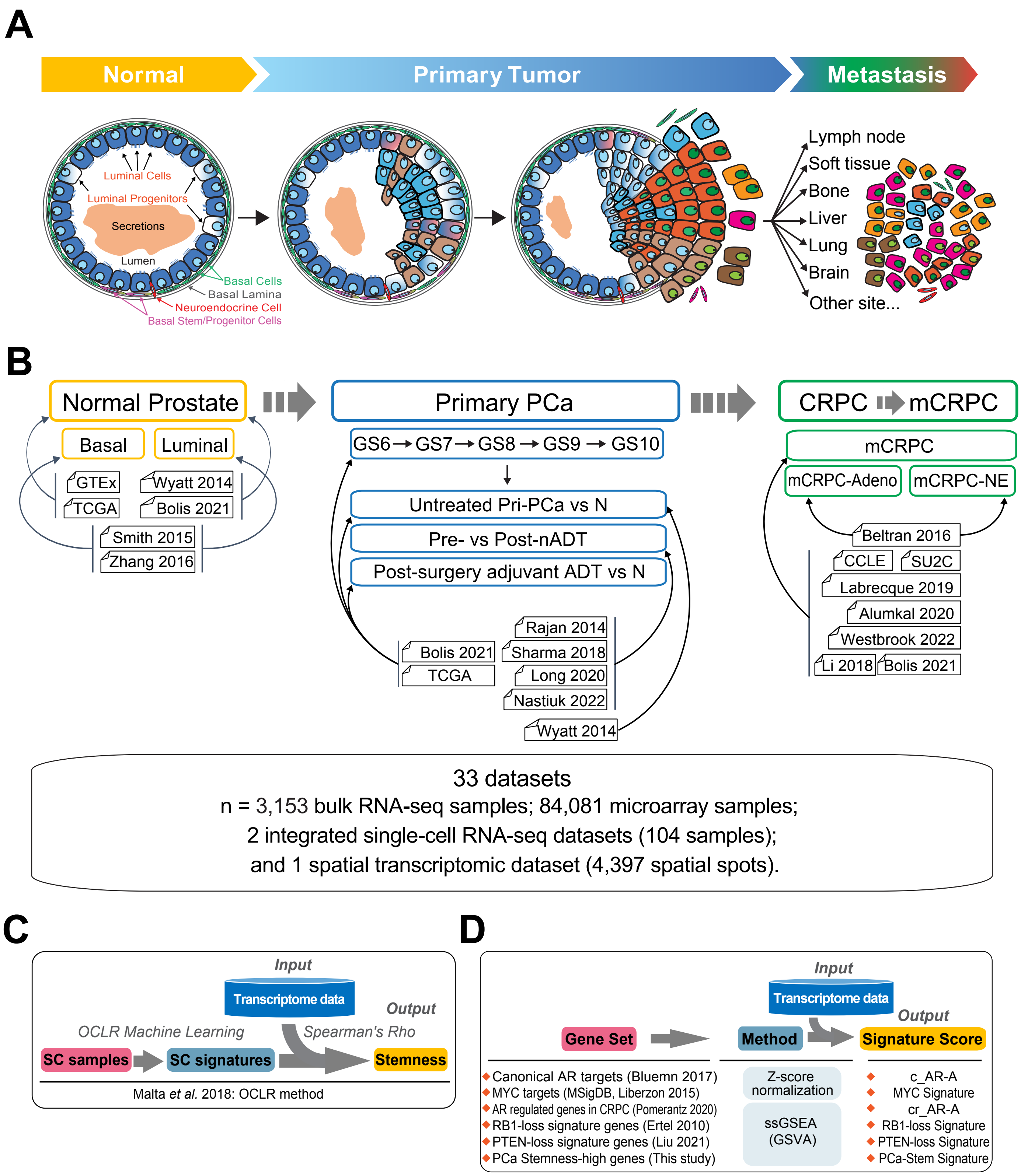
A
Normal
Primary Tumor
Metastasis
Luminal Cells
Luminal Progenitors
Secretions
Lumen
Basal Cells
Basal Lamina
Neuroendocrine Cell
Basal Stem/Progenitor Cells
Lymph node
Soft tissue
Bone
Liver
Lung
Brain
Other site...
B
Normal Prostate
Basal
Luminal
GTEx
TCGA
Wyatt 2014
Bolis 2021
Smith 2015
Zhang 2016
Primary PCa
GS6 → GS7 → GS8 → GS9 → GS10
Untreated Pri-PCa vs N
Pre- vs Post-nADT
Post-surgery adjuvant ADT vs N
Rajan 2014
Bolis 2021
Sharma 2018
TCGA
Long 2020
Nastiuk 2022
Wyatt 2014
CRPC mCRPC
mCRPC
mCRPC-Adeno
mCRPC-NE
Beltran 2016
CCLE
SU2C
Labrecque 2019
Alumkal 2020
Westbrook 2022
Li 2018
Bolis 2021
33 datasets
n = 3,153 bulk RNA-seq samples; 84,081 microarray samples;
2 integrated single-cell RNA-seq datasets (104 samples);
and 1 spatial transcriptomic dataset (4,397 spatial spots).
C
Input
Transcriptome data
Output
OCLR Machine Learning
Spearman's Rho
SC samples
SC signatures
Stemness
Malta et al. 2018: OCLR method
D
Input
Transcriptome data
Output
Gene Set
Method
Signature Score
Canonical AR targets (Bluemn 2017)
MYC targets (MSigDB, Liberzon 2015)
AR regulated genes in CRPC (Pomerantz 2020)
RB1-loss signature genes (Ertel 2010)
PTEN-loss signature genes (Liu 2021)
PCa Stemness-high genes (This study)
Z-score normalization
ssGSEA (GSVA)
c_AR-A
MYC Signature
cr_AR-A
RB1-loss Signature
PTEN-loss Signature
PCa-Stem Signature

**Fig. S1. Schematic illustration of bioinformatic datasets and signature score pipelines used for comparisons across the spectrum of PCa evolution; related to Fig. 1.**

**(A-B)** Schematic illustrating the spectrum of PCa development, therapy resistance, and metastatic progression (A), and related datasets used for comparisons in different contexts (B). Panel A illustrates the PCa progression continuum from Normal Prostate (Normal) through Primary PCa (Pri-PCa), to Castration-Resistant PCa (CRPC) and Metastatic CRPC (mCRPC), with each stage marked by specific changes in cellular composition and architectural organization. Panel B details the datasets utilized for comparative analyses across these stages, with arrows indicating the relationship of each dataset to specific PCa stages. In total, 87,339 transcriptomic profiles from 33 datasets were analyzed in the current study, including 3,153 bulk RNA-seq samples, 84,081 microarray samples, 2 integrated single-cell RNA-seq datasets comprising 104 samples, and 1 spatial transcriptomic dataset comprising 4,397 spatial spots. See Supplementary Table S1 for dataset details. (A is adapted from ref. 7).

Detailed Description of PCa Stages:

- Normal Prostate: Depicted as a glandular organ with a pseudostratified two-layer epithelium consisting of basal and luminal cells, interspersed with rare neuroendocrine cells. Luminal cells express cytokeratins CK8 and CK18, androgen receptor (AR), and prostate-specific antigen (PSA), and represent the primary secretory cells. Also depicted are rare luminal progenitor cells within the luminal cell layer, which represent less than 2 % of the luminal cell population and are characterized by the co-expression of CK5 and CK8 and a low level of CD38 (reviewed in ref. 8). Basal cells express CK5 and CK14, interfacing with the stroma through the basal lamina. The basal cell compartment harbors stem/progenitor cells capable of bidirectional differentiation (ref. 8). Also see an enlarged depiction of a normal human prostatic gland in Supplementary Fig. S3A.

- Localized Pri-PCa: Progresses from well-differentiated structures (low Gleason score) to poorly differentiated configurations (high Gleason score), indicating a shift towards a more stem-like state with loss of glandular architecture. This transition involves a reduction in basal cells and an expansion of progenitor-like cells, with the combined Gleason score (GS) reflecting the architectural patterns of cancerous glands. Gleason pattern 1 represents the most well-differentiated glandular structures and Gleason pattern 5 consists of the most poorly differentiated cells lacking glandular structures. Given that PCa is often multifocal, the combined GS is the sum of the two most prevalent patterns, with GS $\leq$ 7 representing low-grade PCa and GS $\geq$ 8 representing high-grade PCa. Illustrated here are two representative tumor glands with a decrease/loss in basal cells and expansion of luminal progenitor-like cells.

- Metastatic CRPC: Features increased cellular heterogeneity and the presence of cells with stem-like and neuroendocrine-like characteristics, which drive therapy resistance and disease progression.

**(C-D)** Schematic presentation of transcriptome-based stemness quantification method mRNAsi (C, ref. 6), and transcriptome-based signature scores for canonical AR signaling activity levels, MYC signaling activity levels, castration reprogrammed AR regulated genes, RB1-loss signature genes, PTEN-loss signature genes, and PCa-Stem signature (D). See Supplementary Table S2 for gene signature information.

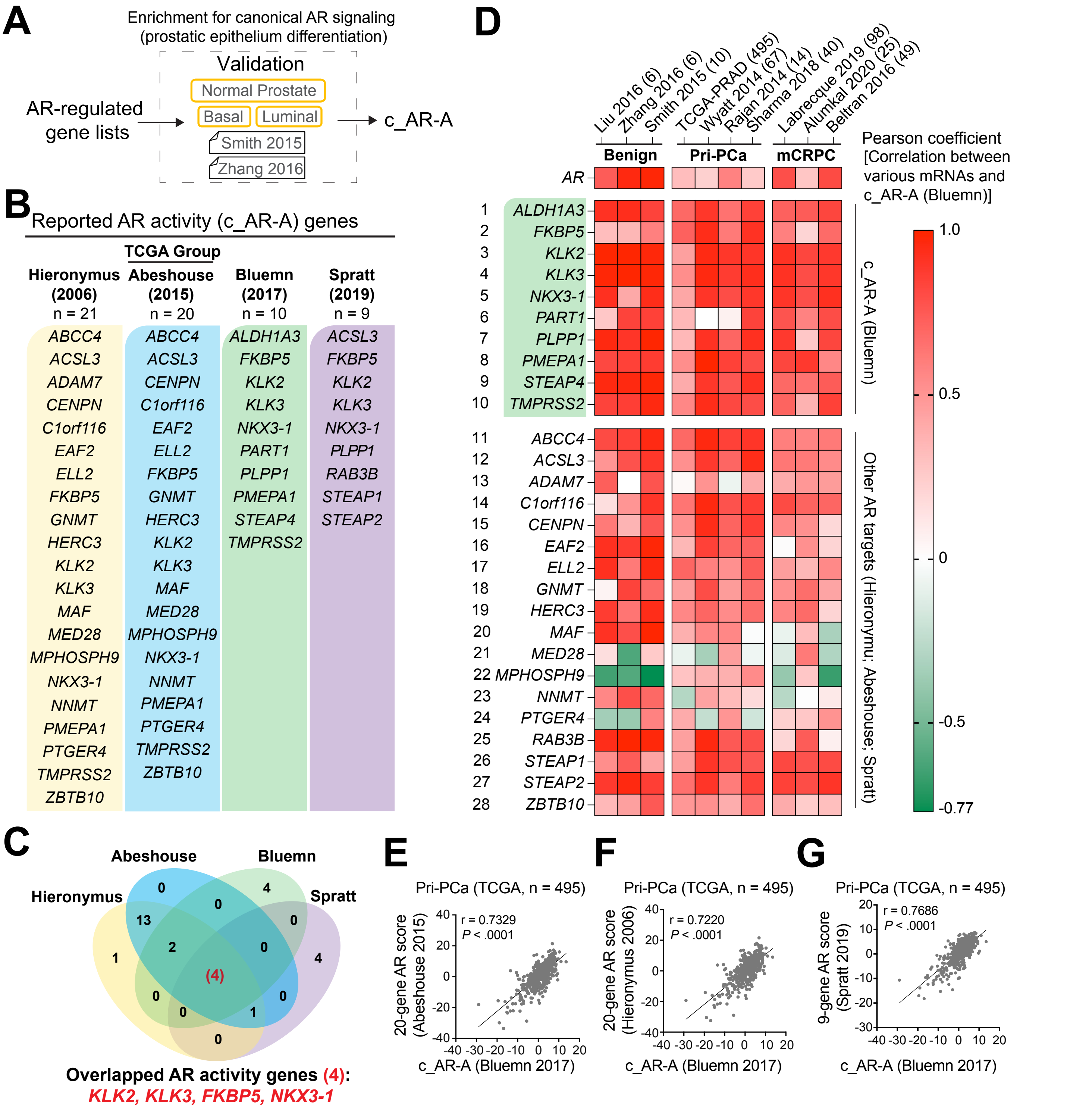


**Fig. S2. Transcriptome-based signature scores for canonical AR signaling activity (c_AR-A); related to Fig. 1.**

**(A)** Workflow for validating canonical AR signaling activity (c_AR-A) signatures from different AR-regulated gene lists.

**(B-C)** Comparison of AR downstream signaling targets from four studies. The 4 genes commonly shared in all four c_AR-A signatures are highlighted (C). Genes were listed using updated HGNC symbols (the HUGO gene nomenclature from the Human Genome Organization). Note that in the original publications, some c_AR-A genes were denoted by their (gene/protein) aliases, as exemplified by *CENPN (BM039), C1orf116 (SARG), KLK3 (PSA), PLPP1 (PPAP2A), and PMEPA1 (TMEPAI)*.

**(D)** c_AR-A signature score highly correlates with mRNA expression levels of canonical AR signaling targets in various PCa cohorts. See Supplementary Table S1 for dataset details.

**(E-G)** Strong positive linear relationships between the Bluemn c_AR-A and the other three c_AR-A signatures (Hieronymus 2006; TCGA Group/Abeshouse 2015, Spratt 2019) using primary PCa cohort (TCGA-PRAD). r indicates Pearson correlation coefficient.

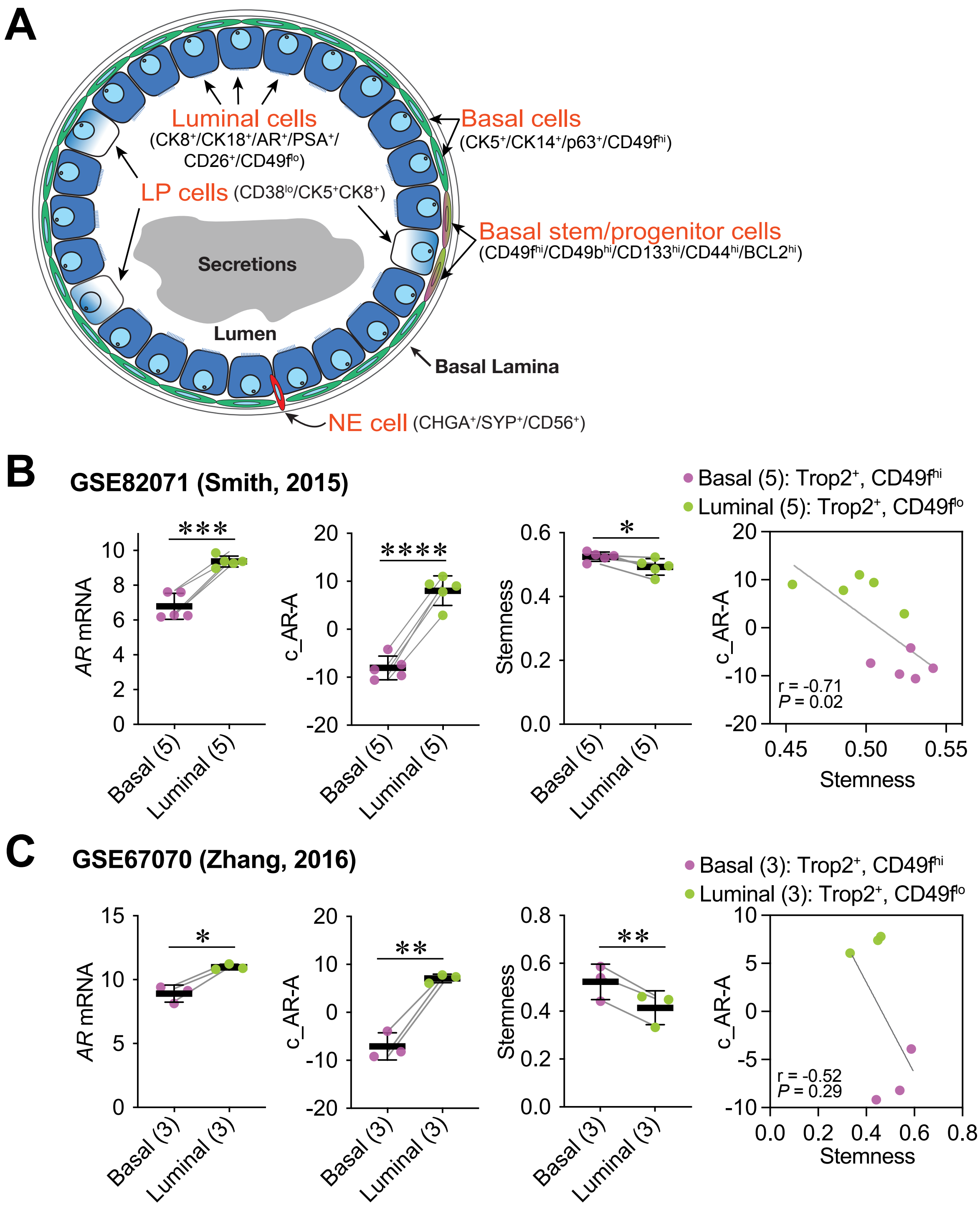


**Fig. S3. Prostate luminal cell compartment harbors higher c_AR-A but lower Stemness as compared to prostate basal cell compartment; related to Fig. 1.**

**(A)** Depiction of the major cell types in human prostatic glands. Both basal and luminal cell compartments harbor differentiated as well as less mature stem/progenitor cells. LP, luminal progenitor; NE, neuroendocrine. Cellular markers are highlighted for different cell types. Adapted from ref. 8.

**(B-C)** Prostate luminal cells possess higher c_AR-A but lower Stemness as compared to prostate basal cells, and Stemness inversely correlates with c_AR-A in the context of prostatic basal and luminal cells. Results are shown as mean ± SD. The pairs are linked with a grey line. Significance was calculated by two-tailed paired Student's *t*-test (*, $P < 0.05$; **, $P < 0.01$; ***, $P < 0.001$; ****, $P < 0.0001$). Pearson correlation coefficients (Pearson's r) were used to calculate the correlations between Stemness and c_AR-A.

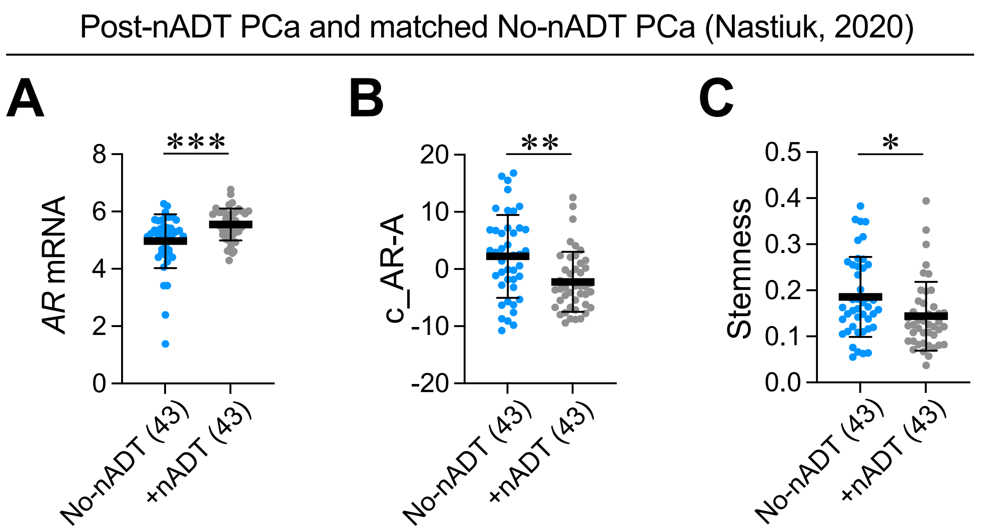


**Fig. S4. nADT decreases both c_AR-A and Stemness; related to Fig. 1.**

Matched comparison of *AR* mRNA levels (A), c_AR-A (B), and Stemness (C) showing significant decrease of both c_AR-A and Stemness in neo-adjuvant ADT (+nADT) group as compared to the samples without nADT (No-nADT) from stage-, tumor grade-, age-matched PCa individuals (RPCI Nastiuk cohort). Sample sizes are indicated. See Supplementary Table S1 for cohort and RNAseq data information. Within the plots, results are shown as mean ± SD. Significance was calculated by two-tailed unpaired Student's *t*-test (*, $P < 0.05$; **, $P < 0.01$; ***, $P < 0.001$).

### Fig. S5

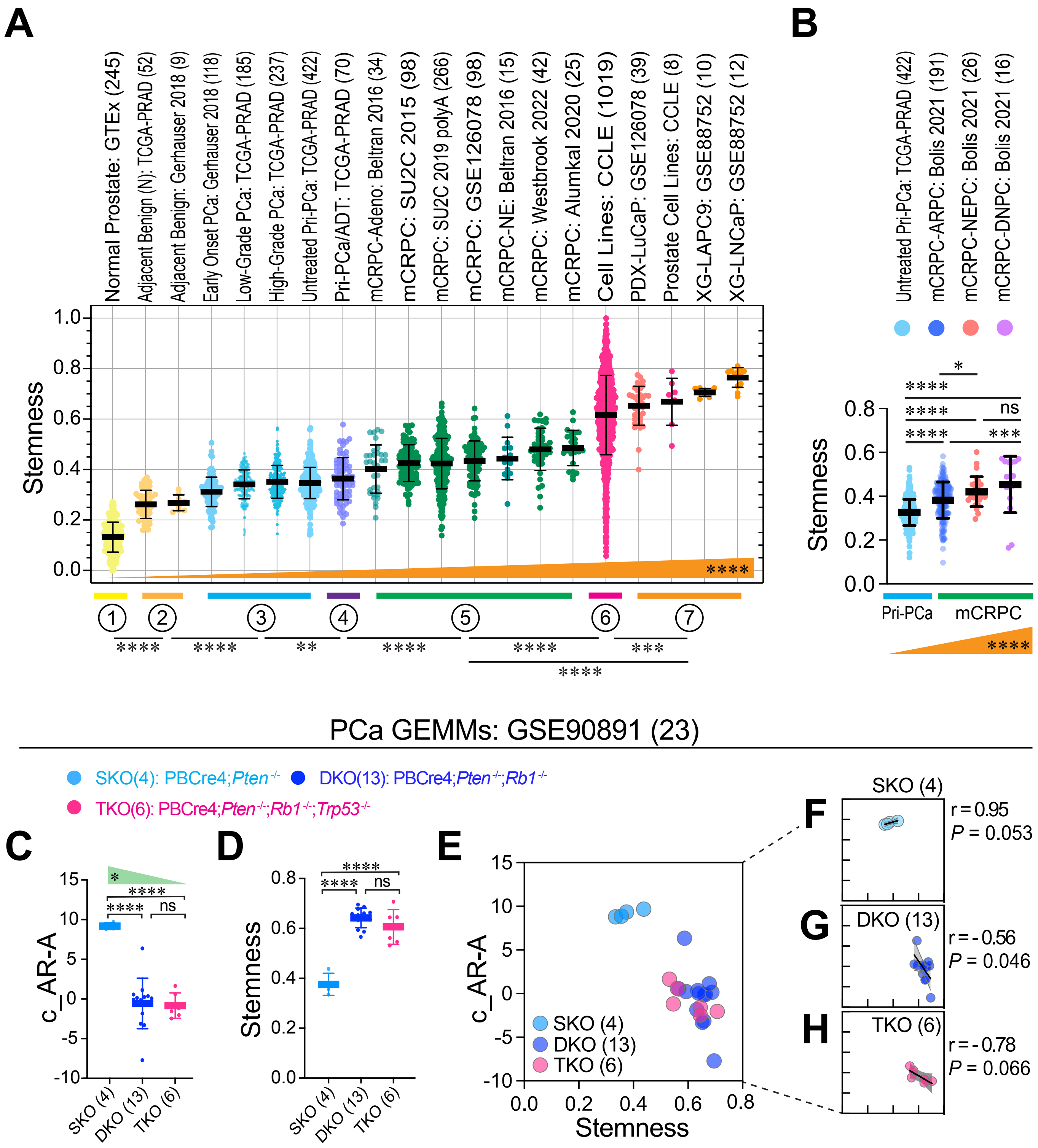
A
B
Normal Prostate: GTEx (245)
Adjacent Benign (N): TCGA-PRAD (52)
Adjacent Benign: Gerhauser 2018 (9)
Early Onset PCa: Gerhauser 2018 (118)
Low-Grade PCa: TCGA-PRAD (185)
High-Grade PCa: TCGA-PRAD (237)
Untreated Pri-PCa: TCGA-PRAD (422)
Pri-PCa/ADT: TCGA-PRAD (70)
mCRPC-Adeno: Beltran 2016 (34)
mCRPC: SU2C 2015 (98)
mCRPC: SU2C 2019 polyA (266)
mCRPC: GSE126078 (98)
mCRPC-NE: Beltran 2016 (15)
mCRPC: Westbrook 2022 (42)
mCRPC: Alumkal 2020 (25)
Cell Lines: CCLE (1019)
PDX-LuCaP: GSE126078 (39)
Prostate Cell Lines: CCLE (8)
XG-LAPC9: GSE88752 (10)
XG-LNCaP: GSE88752 (12)
Stemness
****
① ② ③ ④ ⑤ ⑥ ⑦
**** **** ** **** **** *** ****
Untreated Pri-PCa: TCGA-PRAD (422)
mCRPC-ARPC: Bolis 2021 (191)
mCRPC-NEPC: Bolis 2021 (26)
mCRPC-DNPC: Bolis 2021 (16)
* **** **** ns **** ***
Pri-PCa
mCRPC
PCa GEMMs: GSE90891 (23)
SKO(4): PBCre4;Pten-/-
DKO(13): PBCre4;Pten-/-;Rb1-/-
TKO(6): PBCre4;Pten-/-;Rb1-/-;Trp53-/-
C
D
E
F
G
H
c_AR-A
SKO (4)
DKO (13)
TKO (6)
r = 0.95
P = 0.053
r = -0.56
P = 0.046
r = -0.78
P = 0.066

**Fig. S5. Increased Stemness represents a 'universal' feature of PCa progression and aggressiveness; related to Fig. 3.**

**(A)** Increasing Stemness accompanies PCa progression and aggressiveness. Stemness scores were shown in various PCa cohorts representing the PCa evolutionary spectrum (from top to bottom): GTEx normal prostate → TCGA tumor-adjacent benign prostate tissue → Treatment-naïve Pri-PCa → aggressive Pri-PCa treated with adjuvant hormone therapy (Pri-PCa/ADT) → Treatment-failed CRPC → mCRPC → Long-term PCa models including PDX, xenografts (XG) and cell lines. Sample sizes (n) are indicated in the parentheses. See Supplementary Table S1 for dataset details and Supplementary Table S4 for the statistical significance between different groups.

**(B)** Progressive increase in Stemness from primary PCa to advanced mCRPC subtypes. This violin plot displays increasing Stemness scores across four PCa groups, defined by treatment status and pathway activity: treatment-naïve Pri-PCa from TCGA, and three subtypes of mCRPC categorized by AR and NE pathway activities - AR pathway-active (ARPC), neuroendocrine (NEPC), and double-negative (DNPC). These three subtypes were delineated based on established signatures (Bluemn *et al.*, 2017, ref. 60; Labrecque *et al.,* 2019, ref. 46). Displayed from left to right are: Untreated Pri-PCa (n = 422), mCRPC-ARPC (191), mCRPC-NEPC (26), and mCRPC-DNPC (16). mCRPC samples were derived from studies by Sharp *et al.* (2019, ref. 42, GSE118435), Labrecque *et al.* (2019, ref. 46, GSE126078), Nyquist *et al.* (2020, ref. 43, GSE147250), Lim *et al.* (2021, ref. 45, GSE171729), and Beltran *et al.* (2016, ref. 41, PRJNA282856; dbGaP: phs000909). Normalized RNAseq data were sourced from the Bolis 2021 integrated cohort (ref. 36). For dataset details, see Supplementary Table S1.

**(C-H)** PCa progression in GEMMs is characterized by decreasing c_AR-A and increasing Stemness in aggressive DKO and TKO tumors. The Stemness and c_AR-A were analyzed in distinct genetically engineering mouse models (GEMMs) of PCa due to individual or combined deletion of 3 tumor suppressor genes, *Pten*, *Rb1* and *Trp53* (Ku *et al*., 2017, ref. 49). Briefly, *Pten*$^{-/-}$ single knockout (SKO) prostate tumors develop around 9 weeks, and mice rarely develop metastasis with a median lifespan of 48 weeks. In contrast, the *Pten*$^{-/-}$;*Rb1*$^{-/-}$ double knockout (DKO) mice develop highly metastatic PCa that shortens median survival to ~38 weeks. DKO tumors are initially castration-sensitive but eventually become castration-resistant and, notably, castration-resistant DKO tumors turn into an AR$^{-/lo}$ phenotype. When *Trp53* is further deleted in DKO background, the triple KO (TKO; *Pten*$^{-/-}$*;Rb1*$^{-/-}$*;Trp53*$^{-/-}$) tumors are exclusively AR$^{-}$ NEPC and castration-resistant *de novo* with high metastatic rate and lifespan of ~16 weeks. c_AR-A was dramatically reduced (C) but Stemness was significantly increased (D) in the aggressive DKO and TKO PCa compared to indolent SKO prostate tumors. E-H, Scatter plots illustrating the correlations between c_AR-A and Stemness in combined cases in PCa GEMMs (E), SKO (F, Pearson's r=0.95, $P$ = 0.053), DKO (G, Pearson's r= –0.56, $P$ = 0.053), and TKO (H, Pearson's r= –0.78, $P$ = 0.066).

All data in dot plots and violin plots are shown as mean ± SD. Statistical significance was determined using one-way analysis of variance (ANOVA) followed by Tukey's multiple-comparison test. Jonckheere-Terpstra's trend (J-T) test was used to calculate the statistical significance of the trend across different groups. Pearson correlation coefficients (Pearson's r) were used to calculate the correlations between the Stemness and c_AR-A. Note: ns, not significant; *, $P$ < 0.05; **, $P$ < 0.01; ***, $P$ < 0.001; ****, $P$ < 0.0001.

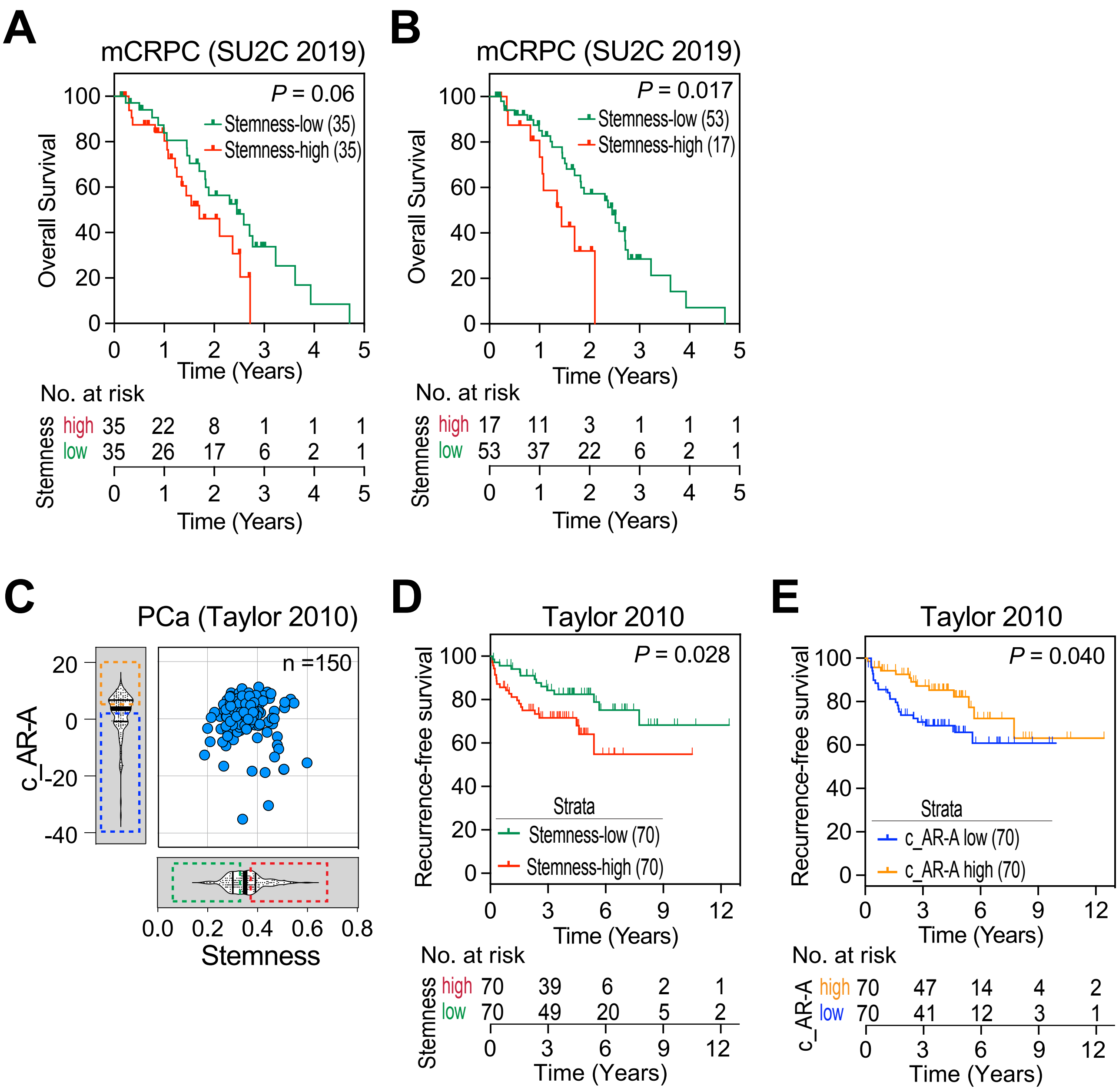


**Fig. S6. High Stemness correlates with poor patient survival; related to Fig. 4.**

**(A-B)** High Stemness correlates with poorer overall survival in SU2C mCRPC cohort. The SU2C 2019 mCRPC cohort was divided into Stemness-high and Stemness-low using Stemness median split (A) or the analysis was done by comparing the 25% of highest Stemness samples with the 75% of lowest Stemness samples (B). *P*-value was determined using the Log-Rank test.

**(C-E)** High Stemness and low c_AR-A correlate with worse recurrence-free survival in PCa patients in the Taylor 2010 PCa cohort. Scatter plot illustrating the relationship between Stemness and c_AR-A in PCa samples from the Taylor 2010 cohort, with marginal violin plots depicting data distributions (median and interquartile range) (C). Stemness (D) or c_AR-A (E) was used to fractionate Taylor cohort for Kaplan-Meier analysis. The Taylor patient cohort was divided into Stemness-high (n = 70) and Stemness-low (n = 70) using Stemness median split (D) or c_AR-A high (n = 70) and c_AR-A low (n =70) using c_AR-A median split (E) followed by the Kaplan-Meier analyses. Note: there are 150 PCa samples with transcriptome data in the Taylor cohort but only 140 PCa samples with survival data. *P*-value was determined using the Log-Rank test.

Fig. S7

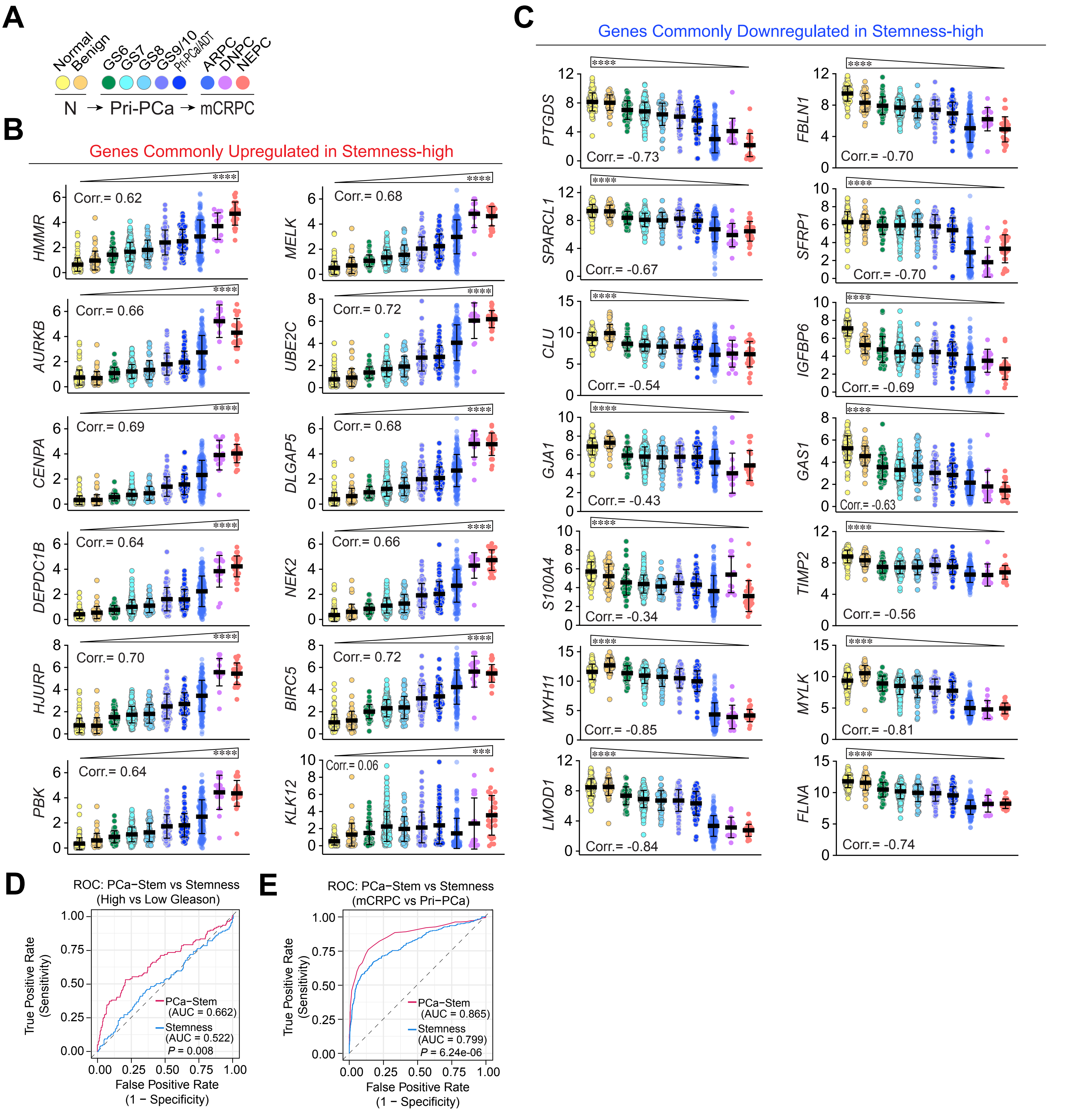
A
Normal
Benign
GS6
GS7
GS8
GS9/10
Pri-PCa/ADT
ARPC
DNPC
NEPC
N → Pri-PCa → mCRPC
B
Genes Commonly Upregulated in Stemness-high
HMMR Corr.= 0.62
MELK Corr.= 0.68
AURKB Corr.= 0.66
UBE2C Corr.= 0.72
CENPA Corr.= 0.69
DLGAP5 Corr.= 0.68
DEPDC1B Corr.= 0.64
NEK2 Corr.= 0.66
HJURP Corr.= 0.70
BIRC5 Corr.= 0.72
PBK Corr.= 0.64
KLK12 Corr.= 0.06
C
Genes Commonly Downregulated in Stemness-high
PTGDS Corr.= -0.73
FBLN1 Corr.= -0.70
SPARCL1 Corr.= -0.67
SFRP1 Corr.= -0.70
CLU Corr.= -0.54
IGFBP6 Corr.= -0.69
GJA1 Corr.= -0.43
GAS1 Corr.= -0.63
S100A4 Corr.= -0.34
TIMP2 Corr.= -0.56
MYH11 Corr.= -0.85
MYLK Corr.= -0.81
LMOD1 Corr.= -0.84
FLNA Corr.= -0.74
D
ROC: PCa−Stem vs Stemness (High vs Low Gleason)
True Positive Rate (Sensitivity)
False Positive Rate (1 − Specificity)
PCa−Stem (AUC = 0.662)
Stemness (AUC = 0.522)
P = 0.008
E
ROC: PCa−Stem vs Stemness (mCRPC vs Pri−PCa)
True Positive Rate (Sensitivity)
False Positive Rate (1 − Specificity)
PCa−Stem (AUC = 0.865)
Stemness (AUC = 0.799)
P = 6.24e-06

**Fig. S7. PCa-Stem signature gene expression correlates with PCa progression; related to Fig. 6.**

**(A)** Color-coded key depicting the PCa progression continuum from normal/benign prostate (N) through various stages of primary PCa (Pri-PCa) to treatment-failed mCRPC. The color scheme in (A) applies to the dot plots in (B–C). Sample groups include:

- Two subgroups in N (Normal/Benign), including GTEx normal prostate (n=116) and TCGA tumor-adjacent benign prostate tissue (n=52).
- Five subgroups in Pri-PCa, including GS6 (n=45), GS7 (n=238), GS8 (n=49), combined GS9 and GS10 (GS9/10; n=90), and aggressive Pri-PCa treated with post-surgery adjuvant hormone therapy (Pri-PCa/ADT; n=70).
- Three subtypes of mCRPC, including AR-active PCa (ARPC; n=191), double-negative AR-null/neuroendocrine-null PCa (DNPC; n=16), and neuroendocrine CRPC (NEPC; n=26).

**(B)** Dot plots showing the mRNA expression of the 12 PCa-Stem signature genes across PCa progression. 11 of the 12 genes (i.e., *HMMR, AURKB, CENPA, DEPDC1B, HJURP, PBK, MELK, UBE2C, DLGAP5, NEK2,* and *BIRC5)* exhibited stage-related increases and positively correlated with PCa progression, whereas *KLK12* did not show a significant progression-associated trend (see Figure 6A; Supplementary Table S3).

**(C)** Dot plots showing the mRNA expression of 14 genes commonly downregulated in Stemness-high groups (i.e., *PTGDS, SPARCL1, CLU, GJA1, S100A4, MYH11, LMOD1, FBLN1, SFRP1, IGFBP6, GAS1, TIMP2, MYLK, and FLNA; see* Supplementary Table S3) all displayed stage-related downregulation and negatively correlated with PCa progression.

**(D)** Receiver operating characteristic (ROC) curves assessing the ability of the 12-gene PCa-Stem signature versus the general Stemness Index to distinguish high-grade (GS >7) from low-grade (GS $\leqslant$7) primary prostate tumors. The PCa-Stem signature outperformed the general Stemness Index, reflecting its enhanced ability to capture aggressiveness-associated transcriptional programs in Pri-PCa.

**(E)** ROC analysis evaluating the discriminative performance of the PCa-Stem signature versus the general Stemness Index in separating metastatic CRPC (mCRPC) from treatment-naïve Pri-PCa. The PCa-Stem signature achieved a higher area under the ROC curve (AUC), indicating improved discriminatory performance for identifying advanced, Stemness-enriched disease states.

For the dot plots in (B–C), y-axes represent normalized mRNA expression on a $\log_2$ scale. Pearson's correlation coefficients (Corr.) indicate associations between individual gene-expression levels and PCa progression scores derived from trajectory inference analysis of the PCa transcriptome atlas (Bolis et al., 2021; ref. 36). Data are presented as mean ± SD, and trends across disease stages were assessed using the Jonckheere-Terpstra (J-T) trend test (wedges above). ***, $P < 0.001$; ****, $P < 0.0001$; ns, not significant. Normalized RNA-seq data across the indicated disease stages were obtained from the integrated transcriptomic framework of PCa (Bolis 2021; ref.36).

### Fig. S8

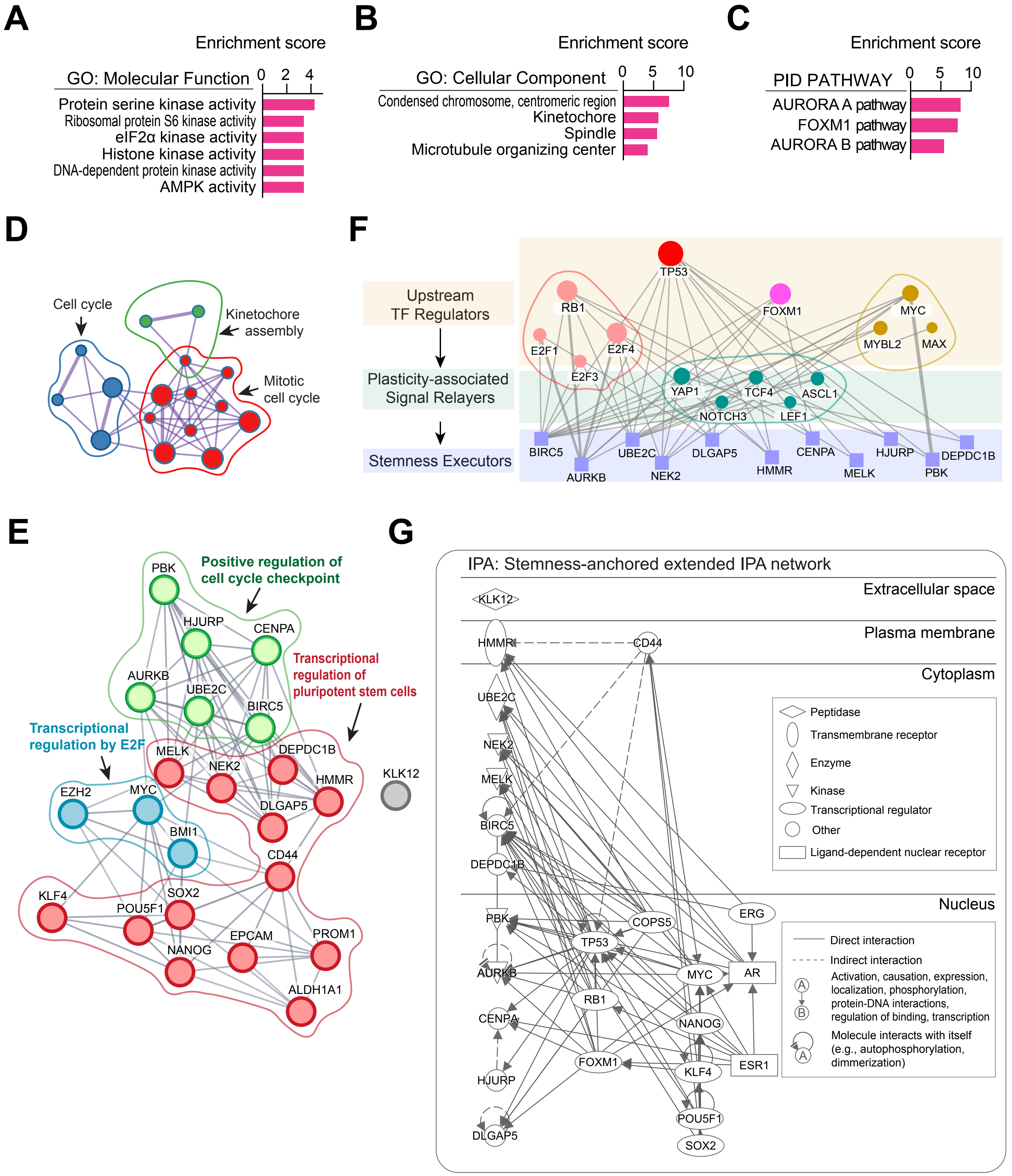
A
Enrichment score
GO: Molecular Function
0
2
4
Protein serine kinase activity
Ribosomal protein S6 kinase activity
eIF2α kinase activity
Histone kinase activity
DNA-dependent protein kinase activity
AMPK activity
B
Enrichment score
GO: Cellular Component
0
5
10
Condensed chromosome, centromeric region
Kinetochore
Spindle
Microtubule organizing center
C
Enrichment score
PID PATHWAY
0
5
10
AURORA A pathway
FOXM1 pathway
AURORA B pathway
D
Cell cycle
Kinetochore assembly
Mitotic cell cycle
F
Upstream TF Regulators
Plasticity-associated Signal Relayers
Stemness Executors
TP53
RB1
FOXM1
MYC
E2F1
E2F4
MYBL2
MAX
E2F3
YAP1
TCF4
ASCL1
NOTCH3
LEF1
BIRC5
UBE2C
DLGAP5
CENPA
HJURP
DEPDC1B
AURKB
NEK2
HMMR
MELK
PBK
E
Positive regulation of cell cycle checkpoint
Transcriptional regulation of pluripotent stem cells
Transcriptional regulation by E2F
PBK
HJURP
CENPA
AURKB
UBE2C
BIRC5
MELK
NEK2
DEPDC1B
HMMR
DLGAP5
EZH2
MYC
BMI1
CD44
KLK12
KLF4
SOX2
POU5F1
NANOG
EPCAM
PROM1
ALDH1A1
G
IPA: Stemness-anchored extended IPA network
Extracellular space
KLK12
Plasma membrane
HMMR
CD44
Cytoplasm
UBE2C
NEK2
MELK
BIRC5
DEPDC1B
Peptidase
Transmembrane receptor
Enzyme
Kinase
Transcriptional regulator
Other
Ligand-dependent nuclear receptor
Nucleus
PBK
COPS5
ERG
TP53
AURKB
MYC
AR
RB1
NANOG
CENPA
FOXM1
ESR1
KLF4
HJURP
POU5F1
DLGAP5
SOX2
Direct interaction
Indirect interaction
Activation, causation, expression, localization, phosphorylation, protein-DNA interactions, regulation of binding, transcription
Molecule interacts with itself (e.g., autophosphorylation, dimmerization)

**Fig. S8. Biological network organization and functional annotation of the 12-gene PCa-Stem signature; related to Fig. 6.**

**(A–C)** Functional enrichment analyses of the 12-gene PCa-Stem signature. **(A)** Gene Ontology (GO) Molecular Function analysis showing enrichment of the indicated GO terms. **(B)** GO Cellular Component analysis showing enrichment of condensed chromosome centromeric region, kinetochore, spindle, and microtubule organizing center, consistent with the predominant involvement of the signature in mitotic cell-cycle regulation. **(C)** PID pathway analysis showing significant enrichment of the AURORA A, FOXM1, and AURORA B pathways. Enrichment scores in (A–C) are presented as $-\log_{10}$−transformed FDR values.

**(D)** Metascape network visualization of enriched biological processes associated with the 12-gene PCa-Stem signature. Functionally related enriched terms are grouped into interconnected clusters, highlighting predominant enrichment of mitotic cell cycle, cell cycle, and kinetochore assembly.

**(E)** Stemness-anchored extended protein–protein interaction (PPI) network generated using STRING. The 12 PCa-Stem signature genes were integrated with representative stemness- and lineage plasticity-associated regulators, including pluripotency factors (POU5F1/OCT4, SOX2, NANOG, and KLF4), MYC, BMI1, EZH2, CD44, EPCAM, PROM1 (CD133), and ALDH1A1. Functional modules associated with transcriptional regulation by E2F, positive regulation of cell-cycle checkpoints, and transcriptional regulation of pluripotent stem cells are indicated.

**(F)** Hierarchical upstream regulatory network inferred from Ingenuity Pathway Analysis (IPA). Upstream transcriptional regulators were organized into three functional layers: Level 1, upstream transcription-factor regulators (RB1, TP53, FOXM1, MYC/MYBL2/MAX, and E2F family members); Level 2, plasticity-associated signal relayers (YAP1, TCF4, ASCL1, NOTCH3, and LEF1); and Level 3, stemness executors, represented by the PCa-Stem signature genes. This organization provides a conceptual framework linking established regulators of lineage plasticity and proliferative signaling with the downstream PCa-Stem program.

**(G)** Stemness-anchored extended regulatory network generated by IPA. The PCa-Stem genes were integrated with established stemness-associated regulators and signaling molecules. Molecules are arranged according to their predominant subcellular localization (extracellular space, plasma membrane, cytoplasm, and nucleus). Solid and dashed lines denote direct and indirect interactions, respectively, as defined by the IPA Knowledge Base.

Fig. S9

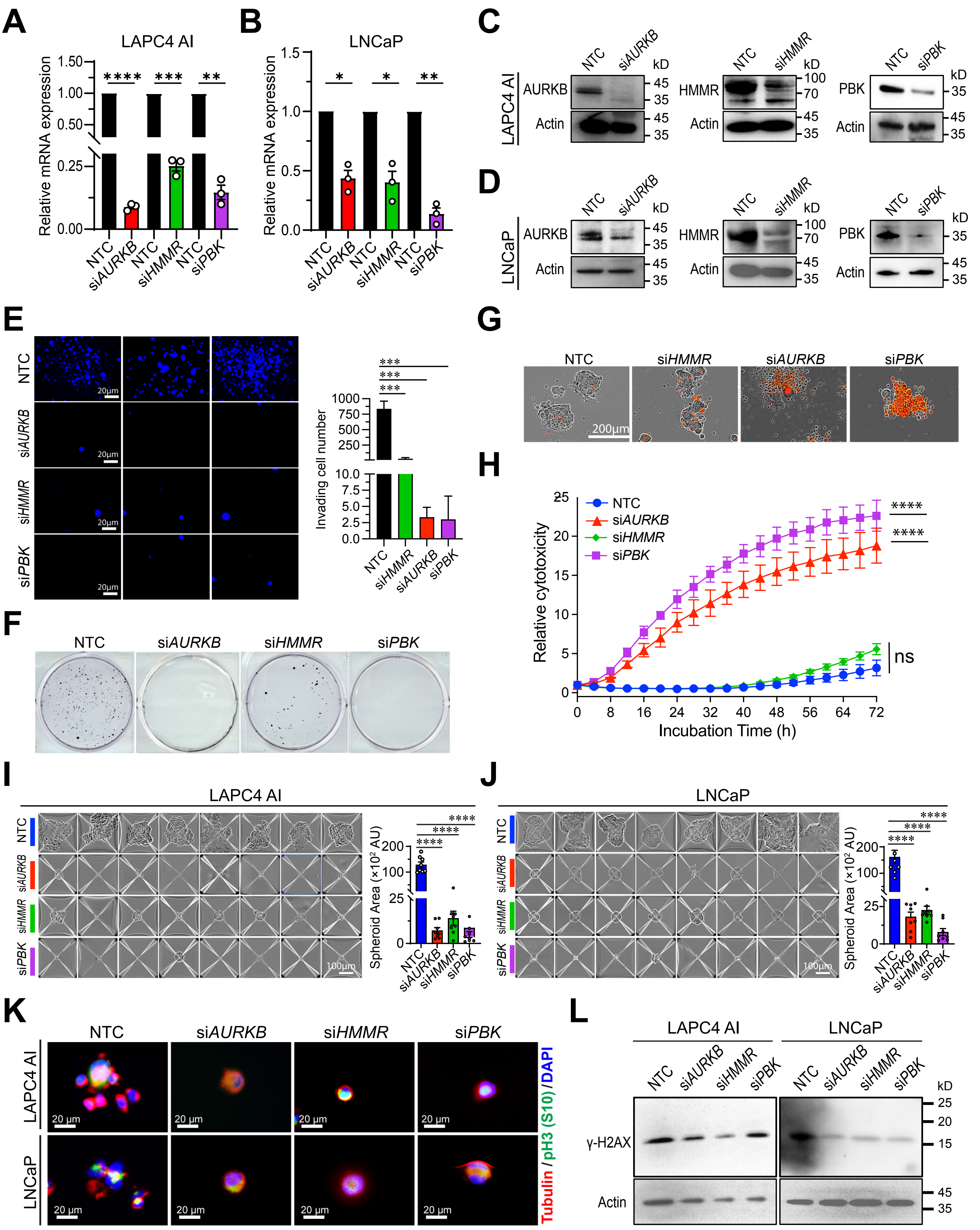

A
LAPC4 AI
Relative mRNA expression
NTC
siAURKB
NTC
siHMMR
NTC
siPBK
B
LNCaP
Relative mRNA expression
C
LAPC4 AI
AURKB
Actin
HMMR
PBK
kD
D
LNCaP
E
NTC
siAURKB
siHMMR
siPBK
20μm
Invading cell number
F
G
200μm
H
Relative cytotoxicity
Incubation Time (h)
ns
I
LAPC4 AI
Spheroid Area (×10² AU)
100μm
J
LNCaP
K
20 μm
Tubulin / pH3 (S10) / DAPI
L
γ-H2AX

**Fig. S9. Functional and molecular characterization of representative PCa-Stem signature genes *AURKB*, *HMMR*, and *PBK* in PCa models; related to Fig. 6.**

**(A–D)** Validation of siRNA-mediated knockdown of *AURKB*, *HMMR*, and *PBK* in LAPC4-AI and LNCaP cells. **(A–B)** qRT-PCR analysis of *AURKB*, *HMMR*, and *PBK* mRNA expression following transfection with the indicated gene-specific siRNAs or non-targeting control (NTC) siRNA in LAPC4-AI (A) and LNCaP (B) cells. Gene expression was normalized to the corresponding NTC (n = 3 biological replicates; mean ± SD; one-sample *t*-test). **(C–D)** Immunoblot validation of AURKB, HMMR, and PBK protein depletion following siRNA-mediated knockdown in LAPC4-AI (C) and LNCaP (D) cells. β-Actin served as a loading control.

**(E)** Matrigel-based Transwell invasion assays in LAPC4-AI cells following siRNA-mediated knockdown of *HMMR*, *AURKB*, or *PBK*. Left, representative fluorescence images of DAPI-stained invading cells; scale bars, 20 µm. Right, quantification of invading cell numbers per field (n = 3 biological replicates; mean ± SD; ordinary one-way ANOVA).

**(F)** Clonogenic survival assay in LAPC4-AI cells following siRNA-mediated knockdown of *HMMR*, *AURKB*, or *PBK*. Cells were replated at clonal density after transfection and cultured for 14 days before fixation and staining. Representative images are shown.

**(G–H)** Three-dimensional spheroid growth and real-time cytotoxicity following knockdown of *HMMR*, *AURKB*, or *PBK* in LAPC4-AI cells. **(G)** Representative phase-contrast/fluorescence overlay images showing spheroid growth and dead-cell accumulation (red reporter); scale bar, 200 µm. **(H)** Real-time cytotoxicity measured over 72 h using the Incucyte platform and a cell-impermeant dead-cell dye. Data are presented as mean ± SD. Note that in this specific spheroid growth assay, *HMMR* knockdown did not elicit significant cytotoxicity (H).

**(I–J)** AggreWell-based prostatosphere assays following siRNA-mediated knockdown of *HMMR*, *AURKB*, or *PBK* in LAPC4-AI (I) and LNCaP (J) cells. Left, representative bright-field images of prostatospheres formed within AggreWell microwells; scale bars, 100 µm. Right, quantification of prostatosphere area (×$10^2$ AU; n = 3 biological replicates; mean ± SD; ordinary one-way ANOVA). The AggreWell-based prostatosphere assay workflow and enlarged representative prostatosphere images are shown in Figure 6F.

**(K)** Double IF staining of phospho-histone H3 (Ser10) (pH3-S10; green), tubulin (red), and DAPI (blue) in LAPC4-AI and LNCaP cells transfected with the indicated siRNAs. Scale bars, 20 µm.

**(L)** Immunoblot analysis of the DNA damage marker γ-H2AX following siRNA-mediated knockdown of *HMMR*, *AURKB*, or *PBK* in LAPC4-AI and LNCaP cells. β-Actin served as a loading control.

Statistical significance is indicated as shown in the individual panels: ns, not significant; *, $P < 0.05$; **, $P < 0.01$; ***, $P < 0.001$; ****, $P < 0.0001$.

### Fig. S10

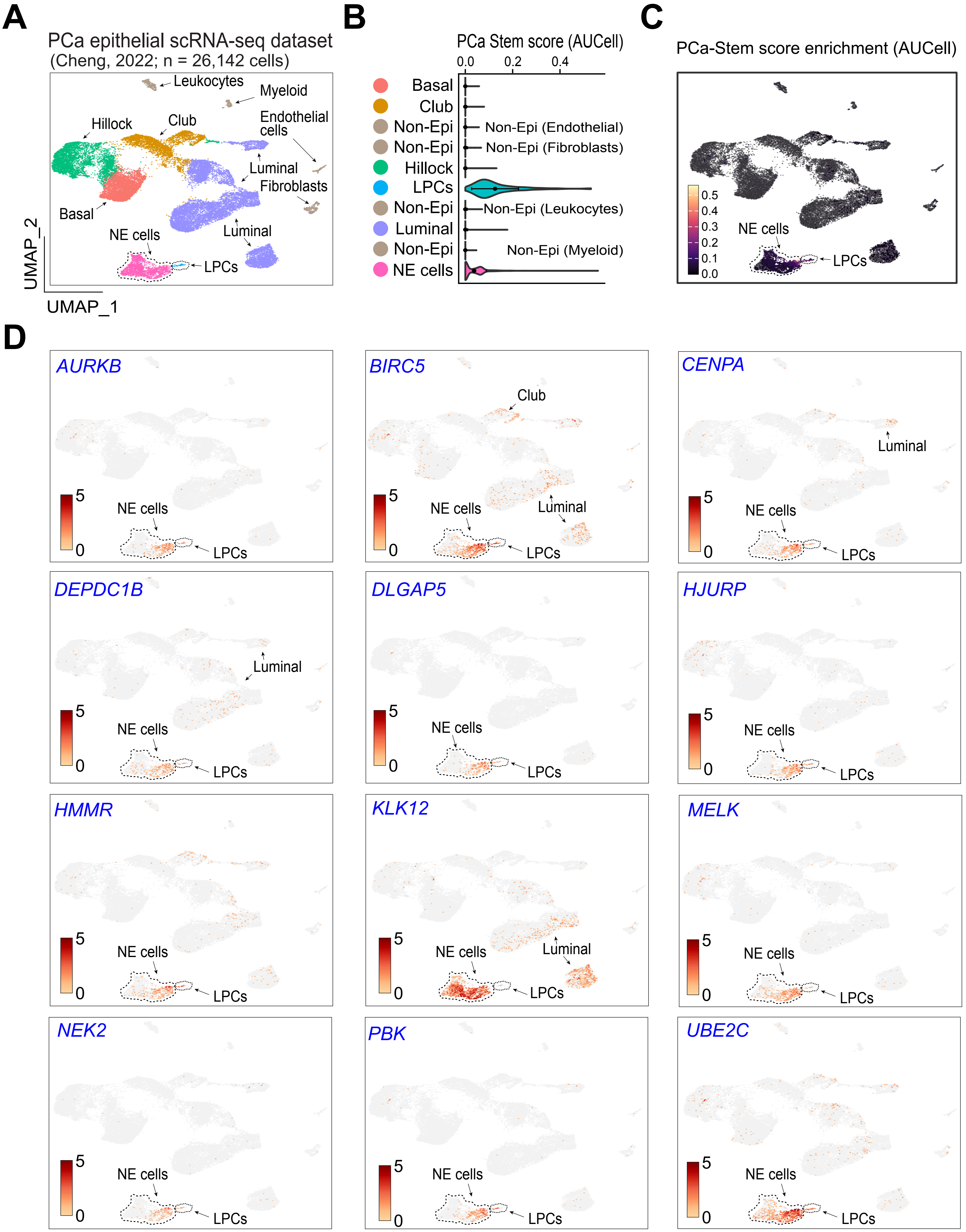
A
PCa epithelial scRNA-seq dataset
(Cheng, 2022; n = 26,142 cells)
Leukocytes
Myeloid
Hillock
Club
Endothelial cells
Luminal
Fibroblasts
Basal
NE cells
LPCs
UMAP_2
UMAP_1
B
PCa Stem score (AUCell)
0.0
0.2
0.4
Basal
Club
Non-Epi
Non-Epi
Hillock
LPCs
Non-Epi
Luminal
Non-Epi
NE cells
Non-Epi (Endothelial)
Non-Epi (Fibroblasts)
Non-Epi (Leukocytes)
Non-Epi (Myeloid)
C
PCa-Stem score enrichment (AUCell)
0.5
0.4
0.3
0.2
0.1
0.0
NE cells
LPCs
D
AURKB
BIRC5
CENPA
DEPDC1B
DLGAP5
HJURP
HMMR
KLK12
MELK
NEK2
PBK
UBE2C
Club
Luminal
NE cells
LPCs
5
0

**Fig. S10. Single-cell expression patterns of the 12 PCa-Stem signature genes in the Cheng 2022 PCa scRNA-seq dataset; related to Fig. 7.**

**(A)** UMAP visualization showing annotated cell populations in the remapped Cheng 2022 PCa scRNA-seq dataset (PMID: 35058087; n = 26,142 cells; ref. 58). The dataset includes treatment-naïve Pri-PCa with matched adjacent benign tissues, locally recurrent CRPC/SCNC, and mCRPC specimens. Major epithelial populations include basal, hillock, club, luminal, lineage plasticity-related cells (LPCs), and neuroendocrine (NE) cells, together with residual non-epithelial populations including endothelial cells, fibroblasts, leukocytes, and myeloid cells. Cell-type annotations followed the classification framework established by Zhao *et al.* (2024) (PMID: 39418984; ref. 57).

**(B)** Comparison of PCa-Stem AUCell scores across the annotated cell populations. PCa-Stem activity was preferentially enriched in LPCs, with lower but detectable enrichment in NE cells.

**(C)** UMAP visualization of PCa-Stem activity quantified by AUCell, showing preferential localization of the PCa-Stem program within LPCs and, to a lesser extent, NE cells.

**(D)** UMAP FeaturePlots showing normalized expression of the individual genes comprising the 12-gene PCa-Stem signature, arranged alphabetically: *AURKB, BIRC5, CENPA, DEPDC1B, DLGAP5, HJURP, HMMR, KLK12, MELK, NEK2, PBK,* and *UBE2C*. All FeaturePlots are displayed using the same $\log_2$ expression scale (0–5) to facilitate direct comparison across genes. Most PCa-Stem signature genes are preferentially expressed in LPCs and NE cells, whereas their expression is generally low in the remaining epithelial and non-epithelial populations.

### Fig. S11

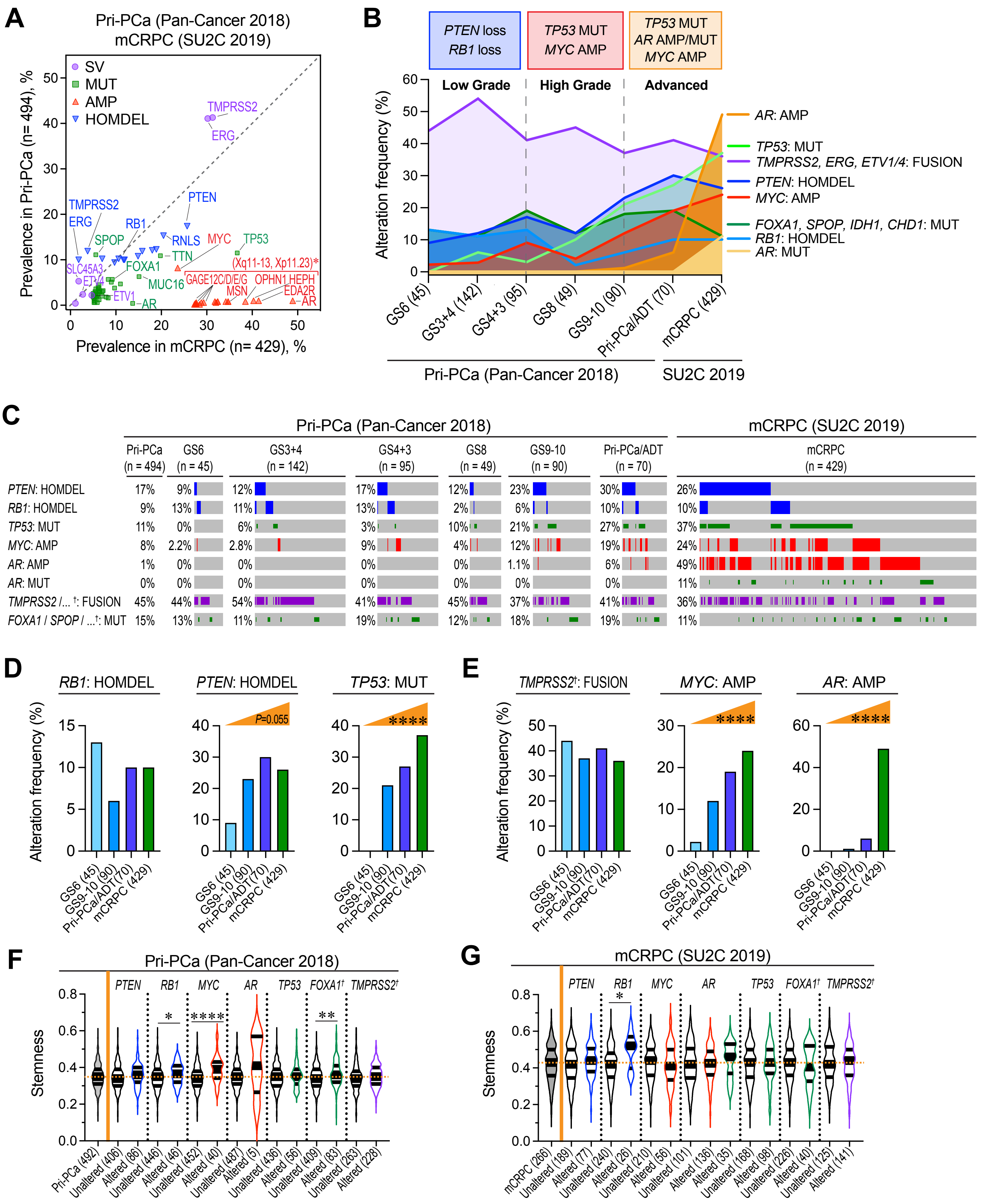

A
Pri-PCa (Pan-Cancer 2018)
mCRPC (SU2C 2019)
SV
MUT
AMP
HOMDEL
TMPRSS2
ERG
PTEN
RB1
SPOP
RNLS
MYC
TP53
TTN
SLC45A3
ETV4
FOXA1
MUC16
ETV1
AR
(Xq11-13, Xp11.23)*
GAGE12C/D/E/G
OPHN1 HEPH
MSN
EDA2R
Prevalence in Pri-PCa (n= 494), %
Prevalence in mCRPC (n= 429), %
B
PTEN loss
RB1 loss
TP53 MUT
MYC AMP
TP53 MUT
AR AMP/MUT
MYC AMP
Low Grade
High Grade
Advanced
Alteration frequency (%)
AR: AMP
TP53: MUT
TMPRSS2, ERG, ETV1/4: FUSION
PTEN: HOMDEL
MYC: AMP
FOXA1, SPOP, IDH1, CHD1: MUT
RB1: HOMDEL
AR: MUT
GS6 (45)
GS3+4 (142)
GS4+3 (95)
GS8 (49)
GS9-10 (90)
Pri-PCa/ADT (70)
mCRPC (429)
Pri-PCa (Pan-Cancer 2018)
SU2C 2019
C
Pri-PCa (Pan-Cancer 2018)
mCRPC (SU2C 2019)
Pri-PCa (n = 494) | GS6 (n = 45) | GS3+4 (n = 142) | GS4+3 (n = 95) | GS8 (n = 49) | GS9-10 (n = 90) | Pri-PCa/ADT (n = 70) | mCRPC (n = 429)
PTEN: HOMDEL 17% 9% 12% 17% 12% 23% 30% 26%
RB1: HOMDEL 9% 13% 11% 13% 2% 6% 10% 10%
TP53: MUT 11% 0% 6% 3% 10% 21% 27% 37%
MYC: AMP 8% 2.2% 2.8% 9% 4% 12% 19% 24%
AR: AMP 1% 0% 0% 0% 0% 1.1% 6% 49%
AR: MUT 0% 0% 0% 0% 0% 0% 0% 11%
TMPRSS2 /... †: FUSION 45% 44% 54% 41% 45% 37% 41% 36%
FOXA1 / SPOP / ...†: MUT 15% 13% 11% 19% 12% 18% 19% 11%
D
RB1: HOMDEL
PTEN: HOMDEL
TP53: MUT
P=0.055
****
Alteration frequency (%)
GS6 (45)
GS9-10 (90)
Pri-PCa/ADT (70)
mCRPC (429)
E
TMPRSS2†: FUSION
MYC: AMP
AR: AMP
****
F
Pri-PCa (Pan-Cancer 2018)
PTEN RB1 MYC AR TP53 FOXA1† TMPRSS2†
Stemness
Pri-PCa (492)
Unaltered (406)
Altered (86)
Unaltered (446)
Altered (46)
Unaltered (452)
Altered (40)
Unaltered (487)
Altered (5)
Unaltered (436)
Altered (56)
Unaltered (409)
Altered (83)
Unaltered (263)
Altered (228)
G
mCRPC (SU2C 2019)
Stemness
mCRPC (266)
Unaltered (189)
Altered (77)
Unaltered (240)
Altered (26)
Unaltered (210)
Altered (56)
Unaltered (101)
Altered (136)
Altered (35)
Unaltered (168)
Altered (98)
Unaltered (226)
Altered (40)
Unaltered (125)
Altered (141)

**Fig. S11. Unique genomic features of Stemness-high PCa; related to Fig. 8.**

**(A)** Scatterplot of observed frequencies of genomic alterations in cancer genes reported in Pri-PCa (TCGA) versus mCRPC (SU2C) with data retrieved from cBioPortal. SV (purple), structural variants/fusions; MUT (green), mutation; AMP (red), amplification in DNA copy numbers (i.e., copy number alterations (CNA) ≥ 2); HOMDEL (blue), homozygous deletion or deep deletion.

**(B)** Genomic alteration frequency plot showing that (early-stage) low-grade PCa have prevalent *PTEN* and *RB1* loss and high-grade PCa have frequent *TP53* MUT and *MYC* AMP whereas advanced PCa have prevalent *TP53* MUT, *MYC* AMP and *AR* AMP/MUT.

**(C)** cBioPortal Oncoprint showing the frequencies of major genetic alteration events along the PCa progression continuum.

**(D-E)** Bar graphs showing genomic alteration frequencies of tumor suppressors (D; *RB1* loss, *PTEN* loss, *TP53* MUT) or oncogenic drivers (E; *TMPRSS2* fusion, *MYC* AMP, AR AMP) across low-grade GS6, high-grade GS9-10, Pri-PCa/ADT, and mCRPC. Colored wedges shown on top indicate the direction of the observed trend. Trend significance across ordered groups was assessed using the Cochran–Armitage trend test results (two-sided). ****, $P < 0.0001$.

**(F-G)** Association of major genomic alterations in PCa with Stemness. Grey, no alterations; Blue, HOMDEL; Green, MUT; Red, AMP; Purple, Fusion. TMPRSS2 /...†: FUSION (in C) or TMPRSS2† (in D-E) stands for combined fusion events in *TMPRSS2*, *ERG*, *ETV1* and *ETV4*; FOXA1 / SPOP /...†: MUT (in C) or FOXA1† (in D-E) represent combined mutation events in *FOXA1*, *SPOP*, *IDH1* and *CHD1*. Note that data presented represent the major mutation subtypes in Pri-PCa (TCGA PRAD cohort) reported in the Cancer Genome Atlas Research Network 2015 (ref. 22) and alterations (mutations, structural variants and copy number) of unknown significance were excluded. Within the violin plots, the center lines represent median values, and box edges are 75th and 25th percentiles. The dashed orange line denotes the median Stemness value calculated across all samples within the indicated cohort, including both altered and unaltered cases. Significance was calculated by Mann-Whitney test (*, $P < 0.05$; **, $P < 0.01$; ****, $P < 0.0001$).

### Fig. S12

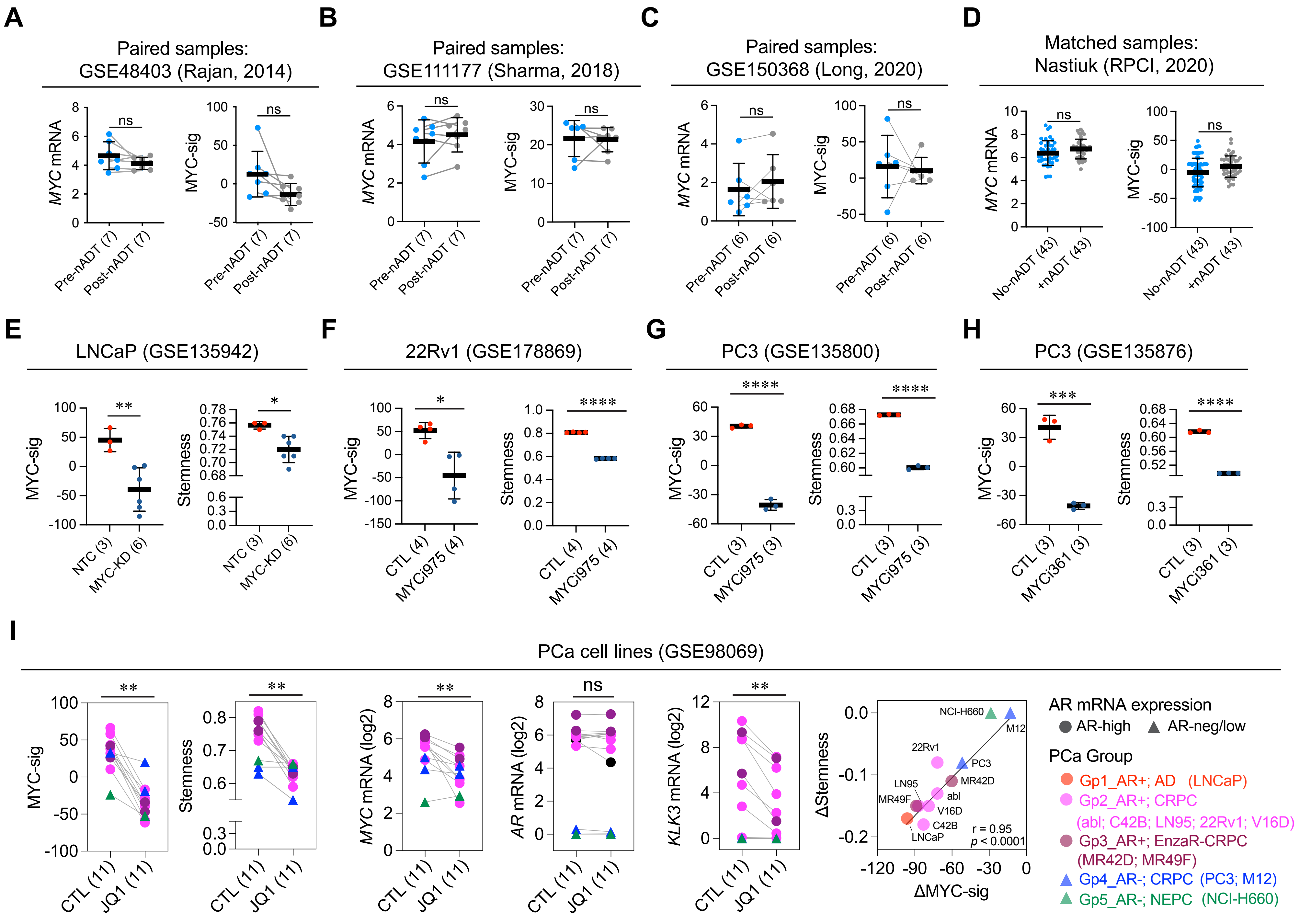
A
Paired samples: GSE48403 (Rajan, 2014)
MYC mRNA
MYC-sig
ns
Pre-nADT (7)
Post-nADT (7)
B
Paired samples: GSE111177 (Sharma, 2018)
C
Paired samples: GSE150368 (Long, 2020)
Pre-nADT (6)
Post-nADT (6)
D
Matched samples: Nastiuk (RPCI, 2020)
No-nADT (43)
+nADT (43)
E
LNCaP (GSE135942)
Stemness
NTC (3)
MYC-KD (6)
F
22Rv1 (GSE178869)
CTL (4)
MYCi975 (4)
G
PC3 (GSE135800)
CTL (3)
MYCi975 (3)
H
PC3 (GSE135876)
MYCi361 (3)
I
PCa cell lines (GSE98069)
CTL (11)
JQ1 (11)
MYC mRNA (log2)
AR mRNA (log2)
KLK3 mRNA (log2)
ΔStemness
ΔMYC-sig
NCI-H660
M12
22Rv1
PC3
MR42D
LN95
abl
MR49F
V16D
C42B
LNCaP
r = 0.95
p < 0.0001
AR mRNA expression
AR-high
AR-neg/low
PCa Group
Gp1_AR+; AD (LNCaP)
Gp2_AR+; CRPC
(abl; C42B; LN95; 22Rv1; V16D)
Gp3_AR+; EnzaR-CRPC
(MR42D; MR49F)
Gp4_AR-; CRPC (PC3; M12)
Gp5_AR-; NEPC (NCI-H660)

**Fig. S12. nADT does not alter MYC activity whereas genetic and pharmacological MYC inhibition suppresses MYC activity and Stemness in diverse PCa models; related to Fig. 9.**

**(A–D)** Matched comparisons showing no significant differences in either *MYC* mRNA expression or MYC activity (MYC-sig) following neoadjuvant ADT (Post-nADT) compared with matched pre-treatment samples (Pre-nADT) from the same individuals in **(A)** the Rajan 2014 cohort (GSE48403), **(B)** the Sharma 2018 cohort (GSE111177), and **(C)** the Long 2020 cohort (GSE150368), or in **(D)** the RPCI Chatta/Nastiuk cohort (ref. 29), comprising matched PCa samples from patients who received neoadjuvant ADT (+nADT) and patients who did not receive ADT (no-ADT), matched for age, tumor stage, and Gleason grade. In panels A–C, each line connects paired samples from the same patient.

**(E)** RNA-seq analysis of LNCaP cells (GSE135942) demonstrates that MYC knockdown (KD) suppresses MYC activity (MYC-sig) and Stemness. Two independent ORF-targeting shMYC constructs (shMYC-512 and shMYC-637), each performed in triplicate, were compared with non-targeting controls (NTC).

**(F)** RNA-seq analysis of AR-positive CRPC 22Rv1 cells treated with the MYC inhibitor MYCi975 (GSE178869). MYCi975 significantly reduced MYC activity (MYC-sig) and Stemness.

**(G-H)** Independent validation in AR-negative PC3 cells treated with MYCi975 (GSE135800) **(G)** or MYCi361 (GSE135876) **(H)**. Both MYC inhibitors consistently reduced MYC activity and Stemness despite the absence of functional AR signaling.

**(I)** RNA-seq analysis of a panel of PCa cell lines treated with the BET bromodomain inhibitor JQ1 (GSE98069). JQ1 significantly reduced MYC activity (MYC-sig), Stemness, and *MYC* and *KLK3* (but not *AR*) mRNA expression. Across all cell lines, the magnitude of Stemness reduction strongly correlated with the reduction in MYC activity (Pearson $r = 0.95$, $P < 0.0001$), supporting a tight association between MYC activity and Stemness across multiple PCa models. AR-high and AR-low/negative cell lines are indicated.

Sample sizes are indicated in parentheses. Results are presented as the mean ± SD. Paired samples in panels A–C and I are connected by gray lines. Statistical significance was determined using two-tailed paired Student's *t*-tests (A–C), two-tailed unpaired Student's *t*-tests (D–H), and the Wilcoxon matched-pairs signed-rank test (I). *ns*, not significant; *, $P < 0.05$; **, $P < 0.01$; ***, $P < 0.001$; ****, $P < 0.0001$. Gene signatures are described in Supplementary Table S2.

### Fig. S13

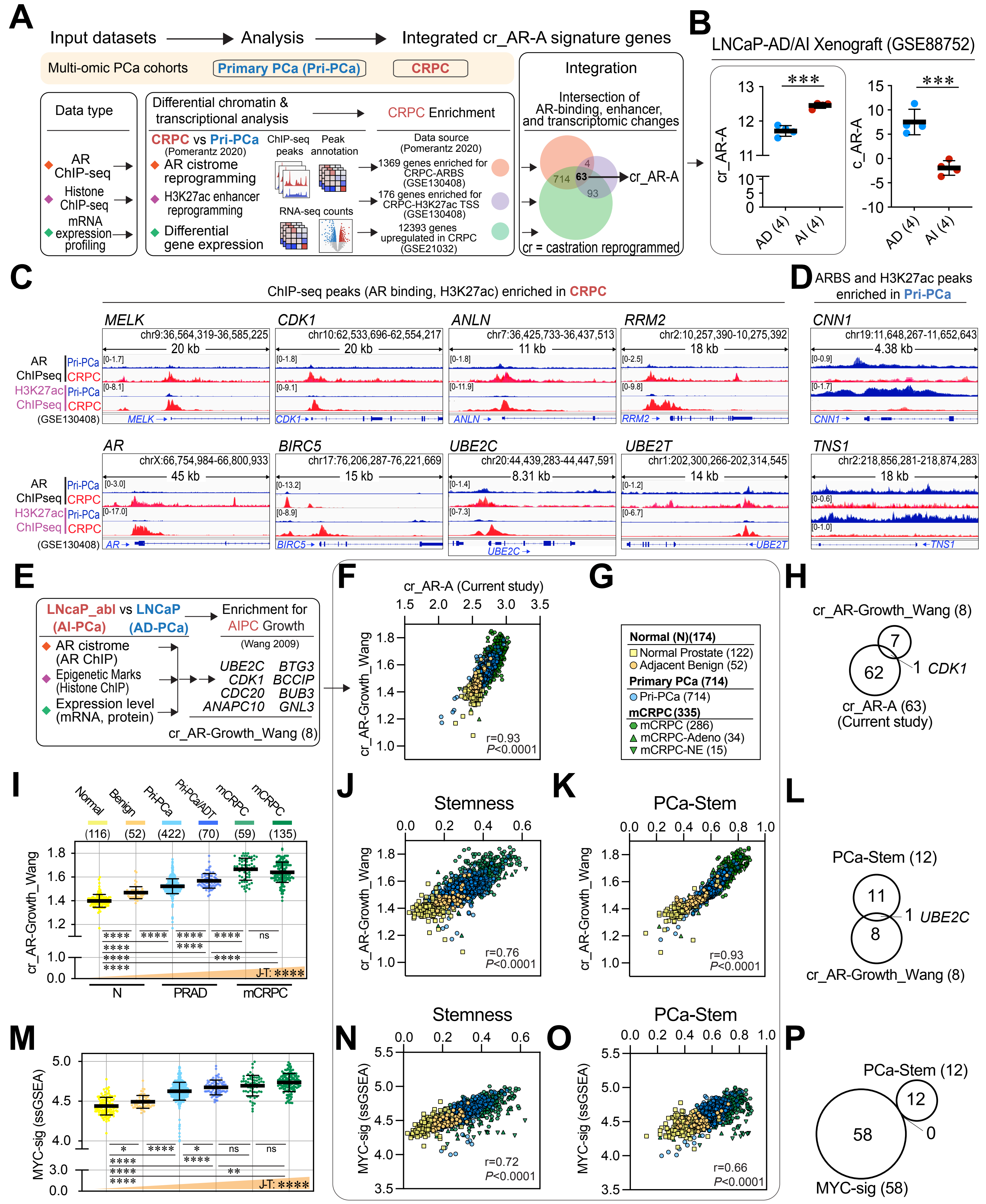
A
Input datasets → Analysis → Integrated cr_AR-A signature genes
Multi-omic PCa cohorts
Primary PCa (Pri-PCa)
CRPC
Data type
AR ChIP-seq
Histone ChIP-seq
mRNA expression profiling
Differential chromatin & transcriptional analysis
CRPC Enrichment
CRPC vs Pri-PCa
(Pomerantz 2020)
AR cistrome reprogramming
H3K27ac enhancer reprogramming
Differential gene expression
ChIP-seq peaks
Peak annotation
RNA-seq counts
Data source (Pomerantz 2020)
1369 genes enriched for CRPC-ARBS (GSE130408)
176 genes enriched for CRPC-H3K27ac TSS (GSE130408)
12393 genes upregulated in CRPC (GSE21032)
Integration
Intersection of AR-binding, enhancer, and transcriptomic changes
714
63
4
93
cr_AR-A
cr = castration reprogrammed
B
LNCaP-AD/AI Xenograft (GSE88752)
cr_AR-A
c_AR-A
AD (4)
AI (4)
***
C
ChIP-seq peaks (AR binding, H3K27ac) enriched in CRPC
AR ChIPseq
H3K27ac ChIPseq
Pri-PCa
CRPC
(GSE130408)
MELK chr9:36,564,319-36,585,225 20 kb
CDK1 chr10:62,533,696-62,554,217 20 kb
ANLN chr7:36,425,733-36,437,513 11 kb
RRM2 chr2:10,257,390-10,275,392 18 kb
AR chrX:66,754,984-66,800,933 45 kb
BIRC5 chr17:76,206,287-76,221,669 15 kb
UBE2C chr20:44,439,283-44,447,591 8.31 kb
UBE2T chr1:202,300,266-202,314,545 14 kb
D
ARBS and H3K27ac peaks enriched in Pri-PCa
CNN1 chr19:11,648,267-11,652,643 4.38 kb
TNS1 chr2:218,856,281-218,874,283 18 kb
E
LNCaP_abl (AI-PCa) vs LNCaP (AD-PCa)
Enrichment for AIPC Growth (Wang 2009)
AR cistrome (AR ChIP)
Epigenetic Marks (Histone ChIP)
Expression level (mRNA, protein)
UBE2C BTG3
CDK1 BCCIP
CDC20 BUB3
ANAPC10 GNL3
cr_AR-Growth_Wang (8)
F
cr_AR-A (Current study)
cr_AR-Growth_Wang
r=0.93
P<0.0001
G
Normal (N)(174)
Normal Prostate (122)
Adjacent Benign (52)
Primary PCa (714)
Pri-PCa (714)
mCRPC (335)
mCRPC (286)
mCRPC-Adeno (34)
mCRPC-NE (15)
H
cr_AR-Growth_Wang (8)
7
62
1 CDK1
cr_AR-A (63)
(Current study)
I
Normal
Benign
Pri-PCa
Pri-PCa/ADT
mCRPC
mCRPC
(116) (52) (422) (70) (59) (135)
cr_AR-Growth_Wang
ns
J-T: ****
N
PRAD
mCRPC
J
Stemness
r=0.76
P<0.0001
K
PCa-Stem
r=0.93
P<0.0001
L
PCa-Stem (12)
11
1 UBE2C
8
cr_AR-Growth_Wang (8)
M
MYC-sig (ssGSEA)
J-T: ****
N
Stemness
r=0.72
P<0.0001
O
PCa-Stem
r=0.66
P<0.0001
P
PCa-Stem (12)
12
58
0
MYC-sig (58)

**Fig. S13. Integrated multi-omic derivation and cross-validation of the castration-reprogrammed AR activity (cr_AR-A) signature; related to Fig. 10.**

**(A)** Schematic illustrating the multi-layer analytic workflow used to define the cr_AR-A signature. AR cistrome reprogramming data from Pomerantz *et al.*, *Nat Genet* 2020 (AR ChIP-seq, GSE130408; ref. 16) were integrated with H3K27ac ChIP-seq (same dataset) and differential expression profiling from Taylor *et al.*, GSE21032 (ref. 47). Genes showing concordant AR binding, enhancer activation, and transcriptional up-regulation in CRPC relative to Pri-PCa were intersected to generate a 63-gene set representing AR–regulated transcriptional outputs preferentially active in mCRPC**,** termed the cr_AR-A signature (see Supplementary Table S2). Candidate gene sets from each layer—AR-bound loci, enhancer-marked regions, and transcriptionally up-regulated genes in mCRPC—are listed in Supplementary Table S6.

**(B)** Gain of cr_AR-A and loss of canonical c_AR-A during the transition from androgen-dependent (AD) to androgen-independent (AI) LNCaP xenografts. Bars represent mean ± SD and *P* values were calculated by two-tailed Student's *t*-test.

**(C–D)** Genome-browser views of representative loci illustrating enhancer-level reprogramming of AR occupancy. *MELK*, *CDK1*, *ANLN*, *RRM2*, *AR*, *BIRC5*, *UBE2C*, and *UBE2T* displayed stronger AR- and H3K27ac-ChIP-seq peaks in CRPC (red) than in Pri-PCa (blue) (C), whereas *CNN1* and *TNS1* showed the opposite pattern (D). These loci were highlighted in Pomerantz *et al.*, *Nat Genet* 2020 (Fig. 1c and Extended Data Fig. 2) as examples of genes co-enriched for metastasis-specific or primary tumor-specific H3K27ac marks with concordant expression changes (ref. 16). Tracks show averaged signal intensity across all Pri-PCa (n=18) and CRPC (n=15) specimens from GSE130408.

**(E)** Comparison schematic for the cr_AR-Growth_Wang 8-gene signature curated from the data of Wang *et al.*, *Cell* 2009 (ref. 14). This gene set was generated by intersecting (i) AR cistrome data from androgen-independent LNCaP-abl versus parental LNCaP cells, (ii) enhancer histone-mark enrichment, and (iii) differential expression profiling, further enriched for genes shown to promote androgen-independent PCa (AIPC) growth. The resulting eight genes, including *UBE2C*, *CDK1*, and *CDC20*, represent a functional AR program active in androgen-independent states.

**(F–H)** Cross-validation of the 63-gene cr_AR-A and the 8-gene cr_AR-Growth_Wang signatures along the PCa progression continuum using the Bolis integrated cohort (total n=1,223, including normal (n=174), Pri-PCa (n=714), and mCRPC (n=335)). (F) A scatter plot showing strong correlation between the two signatures. (G) Details of symbol colors and shapes. (H) Venn diagram indicating one shared gene (*CDK1*).

**(I–L)** Performance of the cr_AR-Growth_Wang signature in the Bolis integrated cohort (n=1,223). (I) cr_AR-Growth_Wang scores increase progressively from normal to mCRPC (datasets as in Fig. 8: normal prostates (GTEx, n=116), tumor-adjacent benign tissues (TCGA-PRAD, n=52), treatment-naïve Pri-PCa (TCGA, n=422), aggressive Pri-PCa treated with adjuvant ADT (Pri-PCa/ADT)(TCGA, n=70), and two mCRPC cohorts including GSE126078 mCRPC (n=59) and SU2C 2015 (n=135). (J–K) Scatter plots showing strong positive correlations between the cr_AR-Growth_Wang signature and Stemness, and between cr_AR-Growth_Wang and PCa-Stem. (L) Venn diagram showing *UBE2C* as the overlapping gene between the cr_AR-Growth_Wang and PCa-Stem signatures.

**(M–P)** Integration with MYC signaling (related to Fig. 7 and 8). (M) Increasing MYC-sig scores along the disease continuum. (N–O) Scatter plots showing strong positive correlations between MYC-sig and Stemness and between MYC-sig and PCa-Stem. (P) Venn diagram showing no direct gene overlap between MYC-sig and PCa-Stem signatures.

All quantitative data are presented as mean ± SD. One-way ANOVA followed by Tukey's multiple-comparison test was used to compare groups, and Jonckheere–Terpstra (J-T) trend tests assessed monotonic trends across disease states. Pearson correlation coefficients (r) were calculated for pairwise associations. *ns*, not significant; *, $P < 0.05$; **, $P < 0.01$; ***, $P < 0.001$; ****, $P < 0.0001$. See Supplementary Table S4 for full statistical summaries and dataset information.

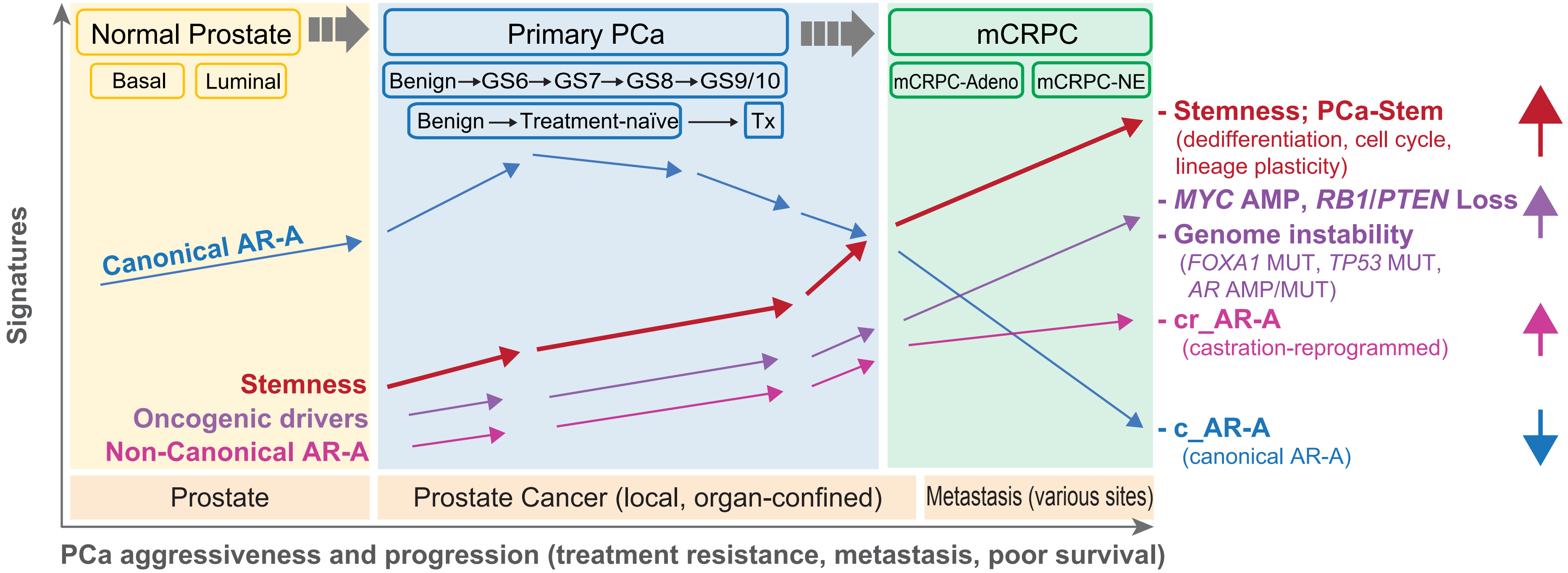


**Fig. S14**. **Schematic summary of PCa aggressiveness and progression illustrating increasing oncogenic Stemness, divergence of canonical and castration-reprogrammed AR signaling, and accumulation of genomic driver events; related to Fig. 10.**

In the current study, we quantified the oncogenic dedifferentiation state (Stemness; indicated by **red arrows**) and canonical AR signaling activity (c_AR-A; **blue arrows**) using transcriptomic data (3,153 bulk RNA-seq and 84,081 microarray samples, together with two independent scRNA-seq resources, and one spatial transcriptomic dataset) collected from 33 datasets encompassing the evolutionary spectrum of PCa development and progression – normal prostate, early-stage and advanced primary tumors, PCa subjected to long-term or neo-adjuvant androgen-deprivation therapy (ADT), and metastatic castration-resistant PCa (mCRPC). Our analysis reveals a continual increase in Stemness along the PCa progression continuum. The c_AR-A, on the other hand, displays a bell-shaped trajectory with an increase in early-stage tumors followed by steady decreases in advanced tumors and the lowest levels in mCRPC. In parallel, the PCa progression is accompanied by the progressive rise of castration-reprogrammed AR activity (cr_AR-A; indicated by **magenta arrows**) together with increasing genome instability and driver events—including MYC activation, *AR* alterations, and tumor suppressor losses (notably *PTEN* and *RB1* loss; indicated by **purple arrows**)—which converge to reinforce the high-Stemness, proliferative, therapy-resistant state in advanced disease/mCRPC. The c_AR-A shows an inverse correlation with the Stemness in mCRPC, and high Stemness correlates with poor patient survival establishing Stemness as a defining feature of PCa progression and aggressiveness.
