## Supplementary material for "Increasing Cancer Stemness Drives Prostate Cancer Progression, Plasticity, Therapy Resistance and Poor Patient Survival": Table S1

**Liu, Jamroze & Cortes et al.: Supplementary Table 1: Datasets Information**

| # | Histology (Sample Type) | Sample Source | Datasets | Reference | PMID | DOI | Sample information | Data type | Data Access Portal |
|---|---|---|---|---|---|---|---|---|---|
| 1 | Normal Prostate | Tissue | GTEx 2013 | GTEx Consortium, Na | 23715323 | 10.1038/ | Normal human prostate from healthy donors (245 from GTEx portal (GTEx v8)) | Bulk RNA-seq; Clinicopathologic data | GTEx portal (gtexportal.org); Xena portal (xenabrowser.net) |
| 2 | Benign Prostate | Primary culture | Smith 2015 | Smith BA et al., PNAS | 26460041 | 10.1073 | 5 human prostate basal (Trop2+ CD49f High); 5 luminal (Trop2+ CD49f Low) from either benign or cancer sample from patients | Bulk RNA-seq | GEO: GSE82071 |
| 3 | Benign Prostate | Primary culture | Liu 2016 | Liu X et al., Cell repor | 27926864 | 10.1016/ | 3 human prostate basal (EpCAM+, CD45-, CD49f Hi, CD38 low); 3 luminal (EpCAM+, CD45-, CD49f Low, CD38 High); 3 luminal progenitor (EpCAM+, CD45-, CD49f Low, CD38 Low) from benign samples collected from patients | Microarray; Bulk RNA-seq | GEO: GSE89050 |
| 4 | Benign Prostate | Primary culture | Zhang 2016 | Zhang D et al., Nat Co | 26924072 | 10.1038/ | 3 human prostate basal (Trop2+ CD49f High); 3 luminal (Trop2+ CD49f Low) from benign samples collected from prostate cancer patients | Bulk RNA-seq | GEO: GSE67070 |
| 5 | PCa, Benign | Tissue | TCGA PRAD 2015; Pan-Cancer Atlas 2018 | Abeshouse A et al., Cell 2015;163(4):1011-1025; Pan-Cancer Atlas 2018; | 26544944<br>24071849<br>29625053<br>29625055 | 10.1016/ | TCGA PRAD 2015: Adjacent benign prostate (52); treatment-naïve Pri-PCa (281); PCa with post-surgery adjuvant ADT (Hormone Tx PCa (n=49));<br>PRAD (Pan-Cancer Atlas 2018): Adjacent benign prostate (52); treatment-naïve pri-PCa (422); PCa with post-surgery adjuvant ADT (Hormone Tx PCa (n=70)); Chemotherapy PCa (2) | Bulk RNA-seq; WES; Clinicopathologic data | Xena portal; cBioportal; firebrowse.org; gdac.broadinstitute.org |
| 6 | PCa (nADT) | Tissue | Rajan 2014 | Rajan P et al., Eur Uro | 24054872 | 10.1016/ | 7 matched pre/post neo-adjuvant ADT PCa samples | Bulk RNA-seq; Clinicopathologic data | GEO: GSE48403 |
| 7 | PCa (nADT) | Tissue | Sharma 2018 | Sharma NV et al., Car | 30314329 | 10.3390/ | 7 matched pre/post neo-adjuvant ADT PCa samples from responder group (defined as "Low Impact Group" in source reference) | Bulk RNA-seq; Clinicopathologic data | GEO: GSE111177 |
| 8 | PCa (nADT), Benign | Tissue | Long 2020 | Long X et al., Cell Dea | 32951005 | 10.1038/ | 6 matched pre/post neo-adjuvant ADT PCa samples; PCa samples (10) and paired adjacent benign prostate tissues (10) obtained from five locally advanced PCa patients | Bulk RNA-seq; Clinicopathologic data | GEO: GSE150368 |
| 9 | PCa (nADT) | Tissue | RPCI Nastiuk 2020 | Jamroze A et al. (Chatta G., Nastiuk KL) medRxiv. [Preprint]. 2026.01.10.26343859. | 41646678 | 10.6489 | Post neo-adjuvant ADT PCa (43); Age-, Stage-, clinically- matched nADT-free PCa (43) | Bulk RNA-seq; Clinicopathologic data | Unpublished RNA-seq and clinicopathologic data from the Nastiuk 2020 cohort were provided by Dr. K. Nastiuk (Roswell Park Comprehensive Cancer Center) and are available upon reasonable request. |
| 10 | PCa, Benign | Tissue | Gerhauser 2018 | Gerhauser C et al., Ca | 30537516 | 10.1016/ | Adjacent benign prostate (9); Early onset PCa (118) | Bulk RNA-seq; Clinicopathologic data | EGA: EGAS00001002923. |
| 11 | PCa, Benign | Tissue | Wyatt 2014 | Wyatt AW et al., Geno | 25155515 | 10.1186/ | Adjacent benign prostate (12); treatment-naïve PCa (12) | Bulk RNA-seq; Clinicopathologic data | The European Nucleotide Archive (ENA): PRJEB6530 |
| 12 | PCa | Primary FFPE tumor tissue | Spratt 2017 | Spratt DE et al., J Clin | 28358655 | 10.1200 | Radical prostatectomy specimens from a multi-instutional study of intermediate and high risk localized disease patients (855) | Microarray; Clinicopathologic data | Veracyte GRID (https://decipherbio.com/grid/) |
| 13 | PCa | Primary FFPE tumor tissue | Tosoian 2020 | Tosoian JJ et al., Pros | 32231245 | 10.1038/ | Radical prostatectomy and biopsy specimens from high risk localized disease patients (405) | Microarray; Clinicopathologic data | Veracyte GRID (https://decipherbio.com/grid/) |
| 14 | PCa | Primary FFPE tumor tissue | CHAARTED correlatives 2021 | Hamid AA et al., Ann Oncol. 2021 Sep;32(9):1157-1166. | 34129855 | 10.1016/ | Biopsy specimens from a Phase 3 trial of metastatic hormone-sensitive patients (160) | Microarray; Clinicopathologic data | NCTN Data Archive: NCT00309985-D14 |
| 15 | PCa | Primary FFPE tumor tissue | Decipher GRID Biopsy | Weiner AB et al. Cancer vol. 129,14 (2023): 2169-2178. doi:10.1002/cncr.34790 | 37060201 | 10.1002/ | Biopsy specimens from a prospectively collected from clinical use of the Decipher test, retrieved from the GRID database (82,470). Transcriptomic profiles from 82,470 prospectively collected prostate biopsy samples were analyzed from the Decipher GRID database (ClinicalTrials.gov identifier: NCT02609269). The data were de-identified according to the Safe Harbor method outlined in the HIPAA Privacy Rule 45 CFR 164.514(b) and (c) (Veracyte, San Diego, CA). These samples, which comprise a large cohort of localized prostate cancer (PCa), were utilized to compare the distribution of the Stemness Index values, focusing on differences across NCCN risk groups.<br>Clinicopathologic Breakdown and Risk Group Analysis: The detailed clinicopathologic data reveal that a majority of the cohort presents with low-grade localized PCa. The breakdown of TNM staging includes:<br>• T1 stage: Approximately 50% of the total samples, indicating early-stage cancer localized within the prostate.<br>• T2 stage: About 8%, representing cancer confined within the prostate but more extensive than T1.<br>• T3 and T4 stages: Less than 1%, suggesting advanced local spread.<br>• N0: Roughly 98%, indicating no regional lymph node involvement.<br>• N1: Approximately 2%, suggesting regional lymph node involvement.<br>The Decipher GRID database categorizes patients into NCCN risk groups as follows, which guide treatment suggestions based on TNM stage, PSA level, and Gleason score:<br>• Very Low and Low Risk: T1-T2a, N0, M0, PSA <10 ng/mL, and Gleason score ≤6. Often managed with active surveillance or less aggressive treatment.<br>• Intermediate Risk - Favorable: Typically T2b-T2c, N0, M0, PSA 10-20 ng/mL, and Gleason score 7. Managed with less aggressive treatment but with more careful monitoring.<br>• Intermediate Risk - Unfavorable: Similar to favorable but might include a higher volume of disease or other adverse features that make the prognosis less certain. Also managed with less aggressive treatment but requires more careful monitoring.<br>• High Risk: Usually involves T3a, N0, M0, PSA >20 ng/mL, or Gleason score 8-10. Often requires aggressive treatment, including a combination of surgery, radiation, and hormonal therapy.<br>• Very High Risk: Typically includes T3b-T4, any N, M0, and high Gleason scores or PSA levels (PSA >20 ng/mL). Patients are at significant risk for metastatic disease and are treated aggressively, often with multi-modal therapies.<br>Observations on Disease Severity and Treatment Naivety: Analysis of the cohort indicates that only 13% of cases fall into the High and Very High risk categories (8,376 + 2,399 = 10,775), suggesting that the majority (approximately 87%) are below Intermediate Risk (~≤ GS7). This distribution implies that most of these lower-grade patients are likely ADT-free due to their less aggressive disease status. | Microarray; Clinicopathologic data | Veracyte GRID (https://decipherbio.com/grid/) |

| | | | | | | | | | |
|---|---|---|---|---|---|---|---|---|---|
| **16** | PCa, CRPC, Benign, Normal Prostate | Tissue | Bolis 2021 | Bolis M et al., Nat Con | 34857732 | 10.1038/ | Bolis 2021 dataset provides normalized RNAseq counts of 1223 clinical samples from an integrated cohort consisting of normal prostate specimens (n = 174), Pri-PCa (n = 714) and mCRPC (n = 335). RNAseq data were downloaded from Zenodo repository (ID: 5546618). As decribed in the reference (PMID: 34857732), this resource of Prostate Cancer Transcriptome Atlas (https://prostatecanceratlas.org) was build and integrated from the following studies/datasets:<br>(1) Genotype-Tissue Expression Database (GTEx; PMID: 23715323);<br>(2) The Cancer Genome Atlas (TCGA, TCGA-PRAD; PMID: 26544944);<br>(3) Atlas of RNA-sequencing profiles of normal human tissues (GSE120795; PMID: 31015567);<br>(4) Integrative epigenetic taxonomy of PNPCa (GSE120741; PMID: 30464211);<br>(5) Prognostic markers in locally advanced lymph node-negative prostate cancer (PRJNA477449);<br>(6) The long noncoding RNA landscape of NEPC and its clinical implications (PRJEB21092; PMID: 29757368);<br>(7) Integrative clinical sequencing analysis of metastatic CRPC reveals a high frequency of clinical actionability (PRJNA283922; dbGaP: phs000915; PMID: 26000489);<br>(8) CSER—exploring precision cancer medicine for sarcoma and rare cancers (PRJNA223419; dbGaP: phs000673; PMID: 28783718);<br>(9) Molecular basis of NEPC (Beltran 2016; PRJNA282856; dbGaP: phs000909; PMID: 26855148);<br>(10) Heterogeneity of androgen receptor splice variant-7 (AR-V7) protein expression and response to therapy in CRPC (GSE118435; PMID: 30334814);<br>(11) Molecular profiling stratifies diverse phenotypes of treatment-refractory metastatic CRPC (PRJNA520923; GEO: GSE126078; PMID: 31361600);<br>(12) Combined TP53 and RB1 Loss Promotes Prostate Cancer Resistance to a Broad Spectrum of Cancer Therapeutics and Confers Vulnerability to Replication Stress (GSE147250; PMID: 32460015);<br>(13) Inter- and intra-tumor heterogeneity of metastatic prostate cancer determined by digital spatial gene expression profiling (GSE147250; PMID: 33658518); and<br>(14) Multiplexed functional genomic analysis of 5' untranslated region mutations across the spectrum of prostate cancer (GSE171729; PMID: 34244513). | Bulk RNA-seq; Clinicopathologic data | Zenodo repository, https://zenodo.org/records/5546618 |
| **17** | PCa, mPCa, Benign | Tissue | Taylor 2010 | Taylor BS et al., Canc | 20579941 | 10.1016/ | Adjacent benign prostate (29); Pri-PCa (131); mPCa (19) | Microarray; Clinicopathologic data | GEO: GSE21034. cBioPortal |
| **18** | mCRPC | Tissue | SU2C 2015 | Robinson D et al., Ce | 26000489 | 10.1016/ | 98 mCRPC samples | Bulk RNA-seq | cBioportal (www.cbioportal.org). |
| **19** | mCRPC | Tissue | SU2C 2019 | Abida W et al., PNAS | 31061129 | 10.1073/ | 266 mCRPC samples | Bulk RNA-seq; WES | cBioportal (www.cbioportal.org). |
| **20** | mCRPC | Tissue, PDX | Labrecque 2019 | Labrecque MP et al., . | 31361600 | 10.1172/ | 98 mCRPC samples; 39 PDX-LuCaP | Bulk RNA-seq | GEO: GSE126078 |
| **21** | mCRPC | Tissue | Alumkal 2020 | Alumkal JJ et al., PNA | 32424106 | 10.1073/ | 25 mCRPC samples before ENZA Tx: 7 nonresponders ( PSA decline <50%) and 18 responders ( PSA decline ≥50%). | Bulk RNA-seq | PMID: 32424106 (supplementary) |
| **22** | mCRPC | Tissue | Westbrook 2022 | Westbrook TC et al., N | 36109521 | 10.1038/ | 42 mCRPC samples: 21 matched pre/post Enzalutamide treatment mCRPC | Bulk RNA-seq | PMID: 36109521 (supplementary) |
| **23** | mCRPC (NEPC, Adeno-mCRPC) | Tissue | Beltran 2016 | Beltran H et al., Natur | 26855148 | 10.1038/ | 15 mCRPC-NE; 34 mCRPC-Adeno | Bulk RNA-seq | cBioportal |
| **24** | NEPC | GEMM tissue | Goodrich 2017 | Ku SY et al., Science | 28059767 | 10.1126/ | 4 SKO (PBCre4:Pten$^{f/f}$); 13 DKO (PBCre4:Pten$^{f/f}$:Rb1$^{f/f}$); 6 TKO (PBCre4:Pten$^{f/f}$:Rb1$^{f/f}$:Trp53$^{f/f}$) | Bulk RNA-seq | GEO: GSE90891 |
| **25** | Various cell lines | Cell line panel | CCLE 2018 | Barretina J et al., Nature 2012;483(7391):603-607;<br>Ghandi M et al. Nature 2019;569(7757):503–508. | 22460905 | 10.1038/ | 1019 human cell lines, including 8 prostate cell lines: DU145, LNCaPcloneFGC, MDAPCA2B, NCIH660, PC3, VCaP, 22RV1; PRECLH | Bulk RNA-seq | depmap portal (depmap.org); Xena portal (xenabrowser.net) |
| **26** | PCa cell line | Xenografts | XG-LNCaP | Li Q et al., Nat Commu | 30190514 | 10.1038/ | 12 LNCaP Xenografts: 4 androgen-dependent PCa; 4 Primary CRPC.; 4 Secondary CRPC (Enzalutamide-resistant) | Bulk RNA-seq | GEO: GSE88752 |
| **26** | PCa cell line | Xenografts | XG-LAPC9 | Li Q et al., Nat Commu | 30190514 | 10.1038/ | 10 LAPC9 Xenografts: 5 androgen-dependent; 5 CRPC (Enzalutamide-resistant) | Bulk RNA-seq | GEO: GSE88752 |
| **27** | Human PCa cell line (LNCaP; AR+ and androgen-sensitive) | Human cell line | GSE135942 | Whitlock NC et al., On | 32681068 | 10.1038/ | LNCaP cells expressing non-targeting shRNA (n = 3) or MYC-targeting shRNAs (512 and 637; n = 6 total). | Bulk RNA-seq | GEO: GSE135942 |
| **28** | Human PCa cell line (22Rv1; AR+ CRPC) | Cell line | GSE178869 | Holmes AG et al., Sci | 35476451 | 10.1126/ | 22Rv1 CRPC cells treated with MYCi975 (10 μM, 48 h) or DMSO control (n=8). | Bulk RNA-seq | GEO: GSE178869 |
| **29** | Human PCa cell line (PC3; AR− CRPC) | Cell line | GSE135800; GSE135876 | Han H et al., Cancer Cell. 2019;36(5):483-497.e15 | 31679823 | 10.1016/ | PC3 cells treated with MYCi975 (8 μM) or MYCi361 (6 μM) for 24 h, each with matched DMSO controls (total n=12). | Bulk RNA-seq | GEO: GSE135800; GSE135876 |
| **30** | Human PCa cell lines | Cell line panel | GSE98069 | Coleman DJ et al., Sc | 30846826 | 10.1038/ | 11 human PCa cell lines treated with JQ1 or vehicle (22 samples): LNCaP (androgen-dependent), abl/C4-2B/LN95/22Rv1/V16D (AR-positive CRPC), MR42D/MR49F (enzalutamide-resistant CRPC), M12/PC3 (AR-negative CRPC), and NCI-H660 (NEPC). | Bulk RNA-seq | GEO: GSE98069 |
| **31** | PCa (ARPC and de novo NEPC) | Primary FFPE tumor tissue | GSE230282 | Watanabe R et al., Int J Mol Sci. 2023;24(10):8955. | 37240308 | 10.3390/ | One FFPE prostate specimen containing spatially adjacent ARPC and de novo NEPC regions (4,397 spatial spots). | Spatial transcriptomics (10x Visium CytAssist) | GEO: GSE230282 |

| | | | | | | | | | |
|---|---|---|---|---|---|---|---|---|---|
| **32** | PCa, CRPC/mCRPC, Benign, Normal Prostate | Tissue | Zhao 2024 | Zhao F et al., EBioMe | 39418984 | 10.1016/ | Integrated human PCa scRNA-seq meta-atlas comprising 93 clinical specimens (222,529 high-quality cells) assembled from 10 previously published human PCa scRNA-seq datasets, including:<br>(1) Henry 2018 (GSE117403; PMID: 30566875; n = 6);<br>(2) Crowley 2020 (GSE150692; PMID: 32915138; n = 3);<br>(3) Dong 2020 (GSE137829; PMID: 33328604; n = 6);<br>(4) Ma 2020 (GSE157703; PMID: 33032611; n = 2);<br>(5) He 2021 (SCP1244/phs001988.v1.p1; PMID: 33664492; n = 15);<br>(6) Kfoury 2021 (GSE143791; PMID: 34719426; n = 16);<br>(7) Tuong 2021 (EGAS00001005787; PMID: 34936871; n = 20); and<br>(8–10) three Chen 2021 datasets (PMID: 33420488), including GSE141445 (n = 13) and two additional datasets from the PCa Cell Atlas resource (www.pradcellatlas.com; n = 3 and n = 9).<br>Following epithelial-cell re-clustering, 89,055 epithelial cells were retained for the present study. | Single-cell RNA-seq | Zenodo repository, https://zenodo.org/records/12605534 |
| **33** | Human prostate epithelial cells (FACS-enriched; Pri-PCa/CRPC/mCRPC) | Human prostate tissues | Cheng 2022 | Cheng Q et al., Eur U | 35058087 | 10.1016/ | Six patients (3 treatment-naïve Pri-PCa including matched adjacent benign prostate tissues from two patients, 2 recurrent CRPC including one SCNC, and 1 mCRPC); 26,142 epithelial cells from 11 FACS-derived epithelial sample-level datasets were analyzed. Epithelial populations were enriched by FACS using Trop2, CD45, CD49f, and CXCR2 markers, including luminal ($Trop2^{+}CD45^{-}CXCR2^{-}$), basal ($Trop2^{+}CD45^{-}CD49f^{+}$), and NE-enriched ($Trop2^{+}CD45^{-}CXCR2^{+}$) populations. | Single-cell RNA-seq; Clinicopathologic data | SRA: PRJNA699369 |
