## Supplementary material for "Increasing Cancer Stemness Drives Prostate Cancer Progression, Plasticity, Therapy Resistance and Poor Patient Survival": Table S2

### Liu, Jamroze & Cortes et al.: Supplementary Table 2: Gene signature Information

#### Index of Gene Signatures Included in Supplementary Table S2

| | Signature Name | Number of Genes | Starting Line No. | Ending Line No. | Biological Description |
|---|---|---|---|---|---|
| 1 | **c_AR-A** | 10 | Line 18 | Line 27 | Canonical AR activity |
| 2 | **MYC-sig** | 58 | Line 32 | Line 89 | MYC transcriptional activity program |
| 3 | **PCa-Stem** | 12 | Line 94 | Line 105 | Prostate Cancer-specific stemness program |
| 4 | **cr_AR-A** | 63 | Line 111 | Line 173 | Castration-reprogrammed AR activity |
| 5 | **cr_AR-Growth_Wang** | 8 | Line 179 | Line 186 | AR-driven androgen-independent proliferation |
| 6 | ***RB1*-loss signature** | 158 | Line 191 | Line 349 | E2F-driven deregulation after RB1 inactivation |
| 7 | ***PTEN*-loss signature** | 45 | Line 354 | Line 398 | Transcriptomic footprint of PTEN deletion |
| 8 | **PCS1/2/3 subtype-enriched genes** | 86/123/219 | Line 403 | Line 621 | PCS1/2/3 subtype-enriched genes |

**1. Canonical AR activity genes (c_AR-A) (10-gene signature)**

Reference: Bluemn, E. G., ... & Nelson, P. S. (2017). Androgen receptor pathway-independent prostate cancer is sustained through FGF signaling. Cancer cell, 32(4), 474-489.

| | AR-A (Bluemn 2017) | Ensembl ID | HGNC Symbol | HGNC gene name | Previous name, Aliases | Mouse Ortholog | AR-A List for NEPC GEMMs |
|---|---|---|---|---|---|---|---|
| 1 | *ALDH1A3* | ENSG00000184254 | ALDH1A3 | Aldehyde Dehydrogenase 1 Family Member A3 | ALDH6 | Aldh1a3 | Aldh1a3 |
| 2 | *FKBP5* | ENSG00000096060 | FKBP5 | FKBP Prolyl Isomerase 5 | FKBP51; FKBP54; PPIase; P54; Ptg-10 | Fkbp5 | Fkbp5 |
| 3 | *KLK2* | ENSG00000167751 | KLK2 | Kallikrein Related Peptidase 2 | | *Egfbp2 | pbsn |
| 4 | *KLK3* | ENSG00000142515 | KLK3 | Kallikrein Related Peptidase 3 | PSA; APS | *Klk1 | |
| 5 | *NKX3-1* | ENSG00000167034 | NKX3-1 | NK3 homeobox 1 | NKX3A; NKX3.1; BAPX2 | Nkx3-1 | Nkx3-1 |
| 6 | *PART1* | ENSG00000152931 | PART1 | Prostate androgen-regulated transcript 1 | | | |
| 7 | *PLPP1* | ENSG00000067113 | PLPP1 | Phospholipid Phosphatase 1 | PPAP2A; PAP-2a; LPP1 | Ppap2a | Ppap2a |
| 8 | *PMEPA1* | ENSG00000124225 | PMEPA1 | Prostate Transmembrane Protein, Androgen Induced 1 | TMEPAI; STAG1 | Pmepa1 | Pmepa1 |
| 9 | *STEAP4* | ENSG00000127954 | STEAP4 | STEAP4 Metalloreductase | TNFAIP9; FLJ23153; TIARP; STAMP2; SchLAH | Steap4 | Steap4 |
| 10 | *TMPRSS2* | ENSG00000184012 | TMPRSS2 | Transmembrane Serine Protease 2 | PRSS10 | Tmprss2 | Tmprss2 |

**2. MYC signaling activity genes (58-gene signature)**

Reference: Liberzon, A., ... & Tamayo, P. (2015). The molecular signatures database hallmark gene set collection. Cell systems, 1(6), 417-425.

| | MYC-sig (Liberzon 2015) | Ensembl ID | HGNC Symbol | HGNC gene name |
|---|---|---|---|---|
| 1 | *AIMP2* | ENSG00000106305 | AIMP2 | aminoacyl tRNA synthetase complex interacting multifunctional protein 2 |
| 2 | *BYSL* | ENSG00000112578 | BYSL | bystin like |
| 3 | *CBX3* | ENSG00000122565 | CBX3 | chromobox 3 |
| 4 | *CDK4* | ENSG00000135446 | CDK4 | cyclin dependent kinase 4 |
| 5 | *DCTPP1* | ENSG00000179958 | DCTPP1 | dCTP pyrophosphatase 1 |
| 6 | *DDX18* | ENSG00000088205 | DDX18 | DEAD-box helicase 18 |
| 7 | *DUSP2* | ENSG00000158050 | DUSP2 | dual specificity phosphatase 2 |
| 8 | *EXOSC5* | ENSG00000077348 | EXOSC5 | exosome component 5 |
| 9 | *FARSA* | ENSG00000179115 | FARSA | phenylalanyl-tRNA synthetase subunit alpha |
| 10 | *GNL3* | ENSG00000163938 | GNL3 | G protein nucleolar 3 |
| 11 | *GRWD1* | ENSG00000105447 | GRWD1 | glutamate rich WD repeat containing 1 |
| 12 | *HK2* | ENSG00000159399 | HK2 | hexokinase 2 |
| 13 | *HSPD1* | ENSG00000144381 | HSPD1 | heat shock protein family D (Hsp60) member 1 |
| 14 | *HSPE1* | ENSG00000115541 | HSPE1 | heat shock protein family E (Hsp10) member 1 |
| 15 | *IMP4* | ENSG00000136718 | IMP4 | IMP U3 small nucleolar ribonucleoprotein 4 |
| 16 | *IPO4* | ENSG00000196497 | IPO4 | importin 4 |
| 17 | *LAS1L* | ENSG00000001497 | LAS1L | LAS1 like ribosome biogenesis factor |
| 18 | *MAP3K6* | ENSG00000142733 | MAP3K6 | mitogen-activated protein kinase kinase kinase 6 |
| 19 | *MCM4* | ENSG00000104738 | MCM4 | minichromosome maintenance complex component 4 |
| 20 | *MCM5* | ENSG00000100297 | MCM5 | minichromosome maintenance complex component 5 |
| 21 | *MPHOSPH10* | ENSG00000124383 | MPHOSPH10 | M-phase phosphoprotein 10 |
| 22 | *MRTO4* | ENSG00000053372 | MRTO4 | MRT4 homolog, ribosome maturation factor |
| 23 | *MYBBP1A* | ENSG00000132382 | MYBBP1A | MYB binding protein 1a |
| 24 | *MYC* | ENSG00000136997 | MYC | MYC proto-oncogene, bHLH transcription factor |
| 25 | *NDUFAF4* | ENSG00000123545 | NDUFAF4 | NADH:ubiquinone oxidoreductase complex assembly factor 4 |
| 26 | *NIP7* | ENSG00000132603 | NIP7 | nucleolar pre-rRNA processing protein NIP7 |
| 27 | *NOC4L* | ENSG00000184967 | NOC4L | nucleolar complex associated 4 homolog |
| 28 | *NOLC1* | ENSG00000166197 | NOLC1 | nucleolar and coiled-body phosphoprotein 1 |
| 29 | *NOP16* | ENSG00000048162 | NOP16 | NOP16 nucleolar protein |
| 30 | *NOP2* | ENSG00000111641 | NOP2 | NOP2 nucleolar protein |
| 31 | *NOP56* | ENSG00000101361 | NOP56 | NOP56 ribonucleoprotein |
| 32 | *NPM1* | ENSG00000181163 | NPM1 | nucleophosmin 1 |
| 33 | *PA2G4* | ENSG00000170515 | PA2G4 | proliferation-associated 2G4 |
| 34 | *PES1* | ENSG00000100029 | PES1 | pescadillo ribosomal biogenesis factor 1 |
| 35 | *PHB1* | ENSG00000167085 | PHB1 | prohibitin 1 |
| 36 | *PLK1* | ENSG00000166851 | PLK1 | polo like kinase 1 |
| 37 | *PLK4* | ENSG00000142731 | PLK4 | polo like kinase 4 |
| 38 | *PPAN* | ENSG00000130810 | PPAN | peter pan homolog |
| 39 | *PPRC1* | ENSG00000148840 | PPRC1 | PPARG related coactivator 1 |
| 40 | *PRMT3* | ENSG00000185238 | PRMT3 | protein arginine methyltransferase 3 |
| 41 | *PUS1* | ENSG00000177192 | PUS1 | pseudouridine synthase 1 |
| 42 | *RABEPK* | ENSG00000136933 | RABEPK | Rab9 effector protein with kelch motifs |
| 43 | *RCL1* | ENSG00000120158 | RCL1 | RNA terminal phosphate cyclase like 1 |

| | | | | |
|---|---|---|---|---|
| 44 | *RRP12* | ENSG00000052749 | RRP12 | ribosomal RNA processing 12 homolog |
| 45 | *RRP9* | ENSG00000114767 | RRP9 | ribosomal RNA processing 9, U3 small nucleolar RNA binding protein |
| 46 | *SLC19A1* | ENSG00000173638 | SLC19A1 | solute carrier family 19 member 1 |
| 47 | *SLC29A2* | ENSG00000174669 | SLC29A2 | solute carrier family 29 member 2 |
| 48 | *SORD* | ENSG00000140263 | SORD | sorbitol dehydrogenase |
| 49 | *SRM* | ENSG00000116649 | SRM | spermidine synthase |
| 50 | *SUPV3L1* | ENSG00000156502 | SUPV3L1 | Suv3 like RNA helicase |
| 51 | *TBRG4* | ENSG00000136270 | TBRG4 | transforming growth factor beta regulator 4 |
| 52 | *TCOF1* | ENSG00000070814 | TCOF1 | treacle ribosome biogenesis factor 1 |
| 53 | *TFB2M* | ENSG00000162851 | TFB2M | transcription factor B2, mitochondrial |
| 54 | *TMEM97* | ENSG00000109084 | TMEM97 | transmembrane protein 97 |
| 55 | *UNG* | ENSG00000076248 | UNG | uracil DNA glycosylase |
| 56 | *UTP20* | ENSG00000120800 | UTP20 | UTP20 small subunit processome component |
| 57 | *WDR43* | ENSG00000163811 | WDR43 | WD repeat domain 43 |
| 58 | *WDR74* | ENSG00000133316 | WDR74 | WD repeat domain 74 |

**3. PCa-Stem signature (12-gene prostate-specific stemness program)**

Reference: This study. Differential gene expression analyses revealed significant differences between the top 33% (Stemness-high) and the bottom 33% (Stemness-low) PCa samples in both treatment-naïve Pri-PCa (Fig. 5B) and treatment-failed mCRPC (Fig. 5C), resulting in a 12-gene 'PCa-Stem signature'.

| | PCa-Stem signature | Ensembl ID | HGNC Symbol | HGNC gene name |
|---|---|---|---|---|
| 1 | *HMMR* | ENSG00000072571 | HMMR | Hyaluronan Mediated Motility Receptor |
| 2 | *AURKB* | ENSG00000178999 | AURKB | Aurora Kinase B |
| 3 | *CENPA* | ENSG00000115163 | CENPA | Centromere Protein A |
| 4 | *DEPDC1B* | ENSG00000035499 | DEPDC1B | DEP Domain Containing 1B |
| 5 | *HJURP* | ENSG00000123485 | HJURP | Holliday Junction Recognition Protein |
| 6 | *PBK* | ENSG00000168078 | PBK | PDZ Binding Kinase |
| 7 | *MELK* | ENSG00000165304 | MELK | Maternal Embryonic Leucine Zipper Kinase |
| 8 | *UBE2C* | ENSG00000175063 | UBE2C | Ubiquitin Conjugating Enzyme E2 C |
| 9 | *DLGAP5* | ENSG00000126787 | DLGAP5 | DLG Associated Protein 5 |
| 10 | *NEK2* | ENSG00000117650 | NEK2 | NIMA Related Kinase 2 |
| 11 | *BIRC5* | ENSG00000089685 | BIRC5 | Baculoviral IAP Repeat Containing 5 |
| 12 | *KLK12* | ENSG00000186474 | KLK12 | Kallikrein Related Peptidase 12 |

**4. Castration-reprogrammed AR-activity signature (cr_AR-A) (63-gene signature)**

Description: Derived by integrating AR and H3K27ac ChIP-seq data from Pomerantz Nat Genet, 2020 (GSE130408) with differential expression from Taylor et al. (GSE21032) to identify genes with CRPC-enriched AR binding, enhancer activation, and transcriptional up-regulation.

Reference: Pomerantz et al. Prostate cancer reactivates developmental epigenomic programs during metastatic progression. Nature genetics 52.8 (2020): 790-799.

| | cr_AR-A signature | Ensembl ID | HGNC Symbol |
|---|---|---|---|
| 1 | *ANLN* | ENSG00000011426 | ANLN |
| 2 | *RRM2* | ENSG00000171848 | RRM2 |
| 3 | *MCM4* | ENSG00000104738 | MCM4 |
| 4 | *MELK* | ENSG00000165304 | MELK |
| 5 | *TYMS* | ENSG00000176890 | TYMS |
| 6 | *CDK1* | ENSG00000170312 | CDK1 |
| 7 | *PLK1* | ENSG00000166851 | PLK1 |
| 8 | *CCNE2* | ENSG00000175305 | CCNE2 |
| 9 | *CCNB1* | ENSG00000134057 | CCNB1 |
| 10 | *AR* | ENSG00000169083 | AR |
| 11 | *SHCBP1* | ENSG00000171241 | SHCBP1 |
| 12 | *MFSD3* | ENSG00000167700 | MFSD3 |
| 13 | *LRRC14* | ENSG00000160959 | LRRC14 |
| 14 | *LAD1* | ENSG00000159166 | LAD1 |
| 15 | *RGS3* | ENSG00000138835 | RGS3 |
| 16 | *DNAAF5* | ENSG00000164818 | DNAAF5 |
| 17 | *LRRC45* | ENSG00000169683 | LRRC45 |
| 18 | *PTK6* | ENSG00000101213 | PTK6 |
| 19 | *STC2* | ENSG00000113739 | STC2 |
| 20 | *GCN1* | ENSG00000089154 | GCN1 |
| 21 | *PKMYT1* | ENSG00000127564 | PKMYT1 |
| 22 | *ZNF695* | ENSG00000197472 | ZNF695 |
| 23 | *CHMP4C* | ENSG00000164695 | CHMP4C |
| 24 | *LACTB2* | ENSG00000147592 | LACTB2 |
| 25 | *PRKDC* | ENSG00000253729 | PRKDC |
| 26 | *BRD9* | ENSG00000028310 | BRD9 |
| 27 | *PPP1R16A* | ENSG00000160972 | PPP1R16A |
| 28 | *DPH7* | ENSG00000148399 | DPH7 |
| 29 | *EVPL* | ENSG00000167880 | EVPL |
| 30 | *MCRIP2* | ENSG00000172366 | MCRIP2 |
| 31 | *CYC1* | ENSG00000179091 | CYC1 |
| 32 | *ARHGEF39* | ENSG00000137135 | ARHGEF39 |
| 33 | *TELO2* | ENSG00000100726 | TELO2 |
| 34 | *SYBU* | ENSG00000147642 | SYBU |
| 35 | *SLC10A5* | ENSG00000253598 | SLC10A5 |
| 36 | *MRM2* | ENSG00000122687 | MRM2 |
| 37 | *OPLAH* | ENSG00000178814 | OPLAH |

| | | | |
|---|---|---|---|
| 38 | *PRR7* | ENSG00000131188 | PRR7 |
| 39 | *INTS8* | ENSG00000164941 | INTS8 |
| 40 | *PRKAR1B* | ENSG00000188191 | PRKAR1B |
| 41 | *ADRM1* | ENSG00000130706 | ADRM1 |
| 42 | *GPAA1* | ENSG00000197858 | GPAA1 |
| 43 | *ESRP1* | ENSG00000104413 | ESRP1 |
| 44 | *ALDOA* | ENSG00000149925 | ALDOA |
| 45 | *UBR5* | ENSG00000104517 | UBR5 |
| 46 | *SAC3D1* | ENSG00000168061 | SAC3D1 |
| 47 | *TESMIN* | ENSG00000132749 | TESMIN |
| 48 | *SLC12A8* | ENSG00000221955 | SLC12A8 |
| 49 | *MXD3* | ENSG00000213347 | MXD3 |
| 50 | *RECQL4* | ENSG00000160957 | RECQL4 |
| 51 | *RBBP8NL* | ENSG00000130701 | RBBP8NL |
| 52 | *ADPRHL1* | ENSG00000153531 | ADPRHL1 |
| 53 | *ZFAND1* | ENSG00000104231 | ZFAND1 |
| 54 | *IARS2* | ENSG00000067704 | IARS2 |
| 55 | *RIDA* | ENSG00000132541 | RIDA |
| 56 | *STAU2* | ENSG00000040341 | STAU2 |
| 57 | *DROSHA* | ENSG00000113360 | DROSHA |
| 58 | *CCDC22* | ENSG00000101997 | CCDC22 |
| 59 | *CPSF1* | ENSG00000071894 | CPSF1 |
| 60 | *JMJD8* | ENSG00000161999 | JMJD8 |
| 61 | *INAVA* | ENSG00000163362 | INAVA |
| 62 | *DPY19L4* | ENSG00000156162 | DPY19L4 |
| 63 | *PHF8* | ENSG00000172943 | PHF8 |

**5. Castration-reprogrammed AR-activity signature enriched in Cell growth (cr_AR-Growth_Wang) (8-gene signature):**

Description: Adopted from Wang et al., Cell 2009, representing AR-bound, enhancer-validated growth drivers active in androgen-independent LNCaP-abl cells (e.g., UBE2C, CDK1, CDC20).

Reference: Wang et al. Androgen receptor regulates a distinct transcription program in androgen-independent prostate cancer. Cell 138.2 (2009): 245-256.

| | **cr_AR-Growth_Wang** | **Ensembl ID** | **HGNC Symbol** |
|---|---|---|---|
| 1 | *UBE2C* | ENSG00000175063 | UBE2C |
| 2 | *CDC20* | ENSG00000117399 | CDC20 |
| 3 | *CDK1* | ENSG00000170312 | CDK1 |
| 4 | *ANAPC10* | ENSG00000164162 | ANAPC10 |
| 5 | *GNL3* | ENSG00000163938 | GNL3 |
| 6 | *BUB3* | ENSG00000154473 | BUB3 |
| 7 | *BTG3* | ENSG00000154640 | BTG3 |
| 8 | *BCCIP* | ENSG00000107949 | BCCIP |

**6. *RB1*-loss Signature (158-gene signature):**

Reference: Ertel et al. (2010). RB-pathway disruption in breast cancer: differential association with disease subtypes, disease-specific prognosis and therapeutic response. Cell cycle, 9(20), 4153-4163.

| | **RB1-loss Signature** | **Ensembl ID** | **HGNC Symbol** |
|---|---|---|---|
| 1 | *AEBP2* | ENSG00000139154 | AEBP2 |
| 2 | *ANAPC11* | ENSG00000141552 | ANAPC11 |
| 3 | *ANAPC5* | ENSG00000089053 | ANAPC5 |
| 4 | *ANGPTL2* | ENSG00000136859 | ANGPTL2 |
| 5 | *ANLN* | ENSG00000011426 | ANLN |
| 6 | *ANP32B* | ENSG00000136938 | ANP32B |
| 7 | *ARHGAP21* | ENSG00000107863 | ARHGAP21 |
| 8 | *ASF1B* | ENSG00000105011 | ASF1B |
| 9 | *ATM* | ENSG00000149311 | ATM |
| 10 | *BIRC5* | ENSG00000089685 | BIRC5 |
| 11 | *BRCA1* | ENSG00000012048 | BRCA1 |
| 12 | *BRCA2* | ENSG00000139618 | BRCA2 |
| 13 | *NCAPH* | ENSG00000121152 | NCAPH |
| 14 | *BUB1* | ENSG00000169679 | BUB1 |
| 15 | *CASP8AP2* | ENSG00000118412 | CASP8AP2 |
| 16 | *CBX2* | ENSG00000173894 | CBX2 |
| 17 | *CBX5* | ENSG00000094916 | CBX5 |
| 18 | *CCNA2* | ENSG00000145386 | CCNA2 |
| 19 | *CCNB1* | ENSG00000134057 | CCNB1 |
| 20 | *CCNB2* | ENSG00000157456 | CCNB2 |
| 21 | *CCNF* | ENSG00000162063 | CCNF |
| 22 | *CD34* | ENSG00000174059 | CD34 |
| 23 | *CDC20* | ENSG00000117399 | CDC20 |
| 24 | *CDC25C* | ENSG00000158402 | CDC25C |
| 25 | *CDC45* | ENSG00000093009 | CDC45 |
| 26 | *CDC6* | ENSG00000094804 | CDC6 |
| 27 | *CDCA3* | ENSG00000111665 | CDCA3 |
| 28 | *CDCA5* | ENSG00000146670 | CDCA5 |
| 29 | *CDCA7* | ENSG00000144354 | CDCA7 |
| 30 | *CDCA8* | ENSG00000134690 | CDCA8 |
| 31 | *CDK2* | ENSG00000123374 | CDK2 |
| 32 | *CDKN1C* | ENSG00000129757 | CDKN1C |

| | | | |
|---|---|---|---|
| 33 | *CDKN3* | ENSG00000100526 | CDKN3 |
| 34 | *CENPA* | ENSG00000115163 | CENPA |
| 35 | *CHAF1B* | ENSG00000159259 | CHAF1B |
| 36 | *CHEK1* | ENSG00000149554 | CHEK1 |
| 37 | *CKAP2* | ENSG00000136108 | CKAP2 |
| 38 | *CKS2* | ENSG00000123975 | CKS2 |
| 39 | *DCK* | ENSG00000156136 | DCK |
| 40 | *DDIT4* | ENSG00000168209 | DDIT4 |
| 41 | *DEK* | ENSG00000124795 | DEK |
| 42 | *DLGAP5* | ENSG00000126787 | DLGAP5 |
| 43 | *DNAJC9* | ENSG00000213551 | DNAJC9 |
| 44 | *DNMT1* | ENSG00000130816 | DNMT1 |
| 45 | *DOK1* | ENSG00000115325 | DOK1 |
| 46 | *DTYMK* | ENSG00000168393 | DTYMK |
| 47 | *E2F1* | ENSG00000101412 | E2F1 |
| 48 | *ECT2* | ENSG00000114346 | ECT2 |
| 49 | *EGR1* | ENSG00000120738 | EGR1 |
| 50 | *EI24* | ENSG00000149547 | EI24 |
| 51 | *EIF4A2* | ENSG00000156976 | EIF4A2 |
| 52 | *ERCC5* | ENSG00000134899 | ERCC5 |
| 53 | *ETV4* | ENSG00000175832 | ETV4 |
| 54 | *EZH2* | ENSG00000106462 | EZH2 |
| 55 | *FBLN1* | ENSG00000077942 | FBLN1 |
| 56 | *FEN1* | ENSG00000168496 | FEN1 |
| 57 | *FEZ1* | ENSG00000149557 | FEZ1 |
| 58 | *FOXM1* | ENSG00000111206 | FOXM1 |
| 59 | *GMNN* | ENSG00000112312 | GMNN |
| 60 | *GTSE1* | ENSG00000075218 | GTSE1 |
| 61 | *H2AZ1* | ENSG00000164032 | H2AZ1 |
| 62 | *HAT1* | ENSG00000128708 | HAT1 |
| 63 | *HELB* | ENSG00000127311 | HELB |
| 64 | *HIP1R* | ENSG00000130787 | HIP1R |
| 65 | *HMGA2* | ENSG00000149948 | HMGA2 |
| 66 | *HMGB2* | ENSG00000164104 | HMGB2 |
| 67 | *HMGB3* | ENSG00000029993 | HMGB3 |
| 68 | *HMGN1* | ENSG00000205581 | HMGN1 |
| 69 | *HMGN2* | ENSG00000198830 | HMGN2 |
| 70 | *HNRNPC* | ENSG00000092199 | HNRNPC |
| 71 | *HNRNPD* | ENSG00000138668 | HNRNPD |
| 72 | *HNRNPR* | ENSG00000125944 | HNRNPR |
| 73 | *HNRNPU* | ENSG00000153187 | HNRNPU |
| 74 | *INCENP* | ENSG00000149503 | INCENP |
| 75 | *KIF11* | ENSG00000138160 | KIF11 |
| 76 | *KIF1C* | ENSG00000129250 | KIF1C |
| 77 | *KIF20A* | ENSG00000112984 | KIF20A |
| 78 | *KIF22* | ENSG00000079616 | KIF22 |
| 79 | *KIF23* | ENSG00000137807 | KIF23 |
| 80 | *KIF2C* | ENSG00000142945 | KIF2C |
| 81 | *KLF4* | ENSG00000136826 | KLF4 |
| 82 | *KPNA2* | ENSG00000182481 | KPNA2 |
| 83 | *LBR* | ENSG00000143815 | LBR |
| 84 | *LIG1* | ENSG00000105486 | LIG1 |
| 85 | *MAD2L1* | ENSG00000164109 | MAD2L1 |
| 86 | *MCM2* | ENSG00000073111 | MCM2 |
| 87 | *MCM3* | ENSG00000112118 | MCM3 |
| 88 | *MCM4* | ENSG00000104738 | MCM4 |
| 89 | *MCM5* | ENSG00000100297 | MCM5 |
| 90 | *MCM7* | ENSG00000166508 | MCM7 |
| 91 | *MDM2* | ENSG00000135679 | MDM2 |
| 92 | *MKI67* | ENSG00000148773 | MKI67 |
| 93 | *MRE11A* | ENSG00000020922 | MRE11A |
| 94 | *MSH2* | ENSG00000095002 | MSH2 |
| 95 | *MTCP1* | ENSG00000214827 | MTCP1 |
| 96 | *NAP1L1* | ENSG00000187109 | NAP1L1 |
| 97 | *NASP* | ENSG00000132780 | NASP |
| 98 | *NEK2* | ENSG00000117650 | NEK2 |
| 99 | *NUSAP1* | ENSG00000137804 | NUSAP1 |
| 100 | *ORC6* | ENSG00000091651 | ORC6 |
| 101 | *PCNA* | ENSG00000132646 | PCNA |
| 102 | *PDCD6IP* | ENSG00000170248 | PDCD6IP |
| 103 | *PDGFRA* | ENSG00000134853 | PDGFRA |
| 104 | *PERP* | ENSG00000112378 | PERP |
| 105 | *PHC1* | ENSG00000111752 | PHC1 |
| 106 | *PHF13* | ENSG00000116273 | PHF13 |
| 107 | *PLK1* | ENSG00000166851 | PLK1 |
| 108 | *PLK4* | ENSG00000142731 | PLK4 |

| | | | |
|---|---|---|---|
| 109 | *PLTP* | ENSG00000100979 | PLTP |
| 110 | *PML* | ENSG00000140464 | PML |
| 111 | *POLD1* | ENSG00000062822 | POLD1 |
| 112 | *PRC1* | ENSG00000198901 | PRC1 |
| 113 | *PRDX4* | ENSG00000123131 | PRDX4 |
| 114 | *PRIM1* | ENSG00000198056 | PRIM1 |
| 115 | *PSIP1* | ENSG00000164985 | PSIP1 |
| 116 | *RACGAP1* | ENSG00000161800 | RACGAP1 |
| 117 | *RAD21* | ENSG00000164754 | RAD21 |
| 118 | *RAD51* | ENSG00000051180 | RAD51 |
| 119 | *RAD51AP1* | ENSG00000111247 | RAD51AP1 |
| 120 | *RBBP4* | ENSG00000162521 | RBBP4 |
| 121 | *RBL1* | ENSG00000080839 | RBL1 |
| 122 | *REV3L* | ENSG00000009413 | REV3L |
| 123 | *RFC2* | ENSG00000049541 | RFC2 |
| 124 | *RFC5* | ENSG00000111445 | RFC5 |
| 125 | *ARHGEF28* | ENSG00000214944 | ARHGEF28 |
| 126 | *RPA1* | ENSG00000132383 | RPA1 |
| 127 | *RRM1* | ENSG00000167325 | RRM1 |
| 128 | *RRM2* | ENSG00000171848 | RRM2 |
| 129 | *SIVA1* | ENSG00000184990 | SIVA1 |
| 130 | *SLBP* | ENSG00000163950 | SLBP |
| 131 | *SLK* | ENSG00000065613 | SLK |
| 132 | *SMC2* | ENSG00000136824 | SMC2 |
| 133 | *SMC4* | ENSG00000113810 | SMC4 |
| 134 | *STMN1* | ENSG00000117632 | STMN1 |
| 135 | *TACC3* | ENSG00000013810 | TACC3 |
| 136 | *TARDBP* | ENSG00000120948 | TARDBP |
| 137 | *TCF19* | ENSG00000137310 | TCF19 |
| 138 | *TFDP1* | ENSG00000198176 | TFDP1 |
| 139 | *TK1* | ENSG00000167900 | TK1 |
| 140 | *TMPO* | ENSG00000120802 | TMPO |
| 141 | *TOP2A* | ENSG00000131747 | TOP2A |
| 142 | *TOPBP1* | ENSG00000163781 | TOPBP1 |
| 143 | *TRIP13* | ENSG00000071539 | TRIP13 |
| 144 | *TTK* | ENSG00000112742 | TTK |
| 145 | *TUBGCP3* | ENSG00000126216 | TUBGCP3 |
| 146 | *TYMS* | ENSG00000176890 | TYMS |
| 147 | *UHRF1* | ENSG00000276043 | UHRF1 |
| 148 | *UPF3B* | ENSG00000125351 | UPF3B |
| 149 | *USP1* | ENSG00000162607 | USP1 |
| 150 | *WAC* | ENSG00000095787 | WAC |
| 151 | *WDHD1* | ENSG00000198554 | WDHD1 |
| 152 | *ZMAT3* | ENSG00000172667 | ZMAT3 |
| 153 | *XRCC1* | ENSG00000073050 | XRCC1 |
| 154 | *AURKA* | ENSG00000087586 | AURKA |
| 155 | *AURKB* | ENSG00000178999 | AURKB |
| 156 | *CCNE1* | ENSG00000105173 | CCNE1 |
| 157 | *CCNE2* | ENSG00000175305 | CCNE2 |
| 158 | *CDT1* | ENSG00000167513 | CDT1 |
| 159 | *DBF4* | ENSG00000006634 | DBF4 |

**7.*PTEN*-loss Signature (45-gene signature):**

Reference: Liu et al. (2021). Tumor subtype defines distinct pathways of molecular and clinical progression in primary prostate cancer. The Journal of clinical investigation, 131(10).

| | PTEN-loss Signature | Ensembl ID | HGNC Symbol |
|---|---|---|---|
| 1 | *ABHD12* | ENSG00000100997 | ABHD12 |
| 2 | *AGRN* | ENSG00000188157 | AGRN |
| 3 | *AMMECR1* | ENSG00000101935 | AMMECR1 |
| 4 | *ANO10* | ENSG00000160746 | ANO10 |
| 5 | *ARF3* | ENSG00000134287 | ARF3 |
| 6 | *ASNA1* | ENSG00000198356 | ASNA1 |
| 7 | *ATAD1* | ENSG00000138138 | ATAD1 |
| 8 | *ATP12A* | ENSG00000075673 | ATP12A |
| 9 | *C1orf115* | ENSG00000162817 | C1orf115 |
| 10 | *CCDC6* | ENSG00000108091 | CCDC6 |
| 11 | *CCNO* | ENSG00000152669 | CCNO |
| 12 | *CD276* | ENSG00000103855 | CD276 |
| 13 | *CDR2L* | ENSG00000109089 | CDR2L |
| 14 | *CLSTN3* | ENSG00000139182 | CLSTN3 |
| 15 | *CUTC* | ENSG00000119929 | CUTC |
| 16 | *ECE1* | ENSG00000117298 | ECE1 |
| 17 | *ENC1* | ENSG00000171617 | ENC1 |
| 18 | *EPB41L4A-AS1* | ENSG00000224032 | EPB41L4A-AS1 |
| 19 | *FAM19A5* | ENSG00000219438 | FAM19A5 |
| 20 | *FAM35A* | ENSG00000122376 | FAM35A |
| 21 | *FKBP1A* | ENSG00000088832 | FKBP1A |

| | | | |
|---|---|---|---|
| 22 | *GJC3* | ENSG00000176402 | GJC3 |
| 23 | *GLB1* | ENSG00000170266 | GLB1 |
| 24 | *GLUD1* | ENSG00000148672 | GLUD1 |
| 25 | *GP2* | ENSG00000169347 | GP2 |
| 26 | *KHDRBS3* | ENSG00000131773 | KHDRBS3 |
| 27 | *KPNA2* | ENSG00000182481 | KPNA2 |
| 28 | *LACTB* | ENSG00000103642 | LACTB |
| 29 | *LAT2* | ENSG00000086730 | LAT2 |
| 30 | *LRRN1* | ENSG00000175928 | LRRN1 |
| 31 | *MLX* | ENSG00000108788 | MLX |
| 32 | *PCCB* | ENSG00000114054 | PCCB |
| 33 | *PEX10* | ENSG00000157911 | PEX10 |
| 34 | *PTEN* | ENSG00000171862 | PTEN |
| 35 | *RAB6B* | ENSG00000154917 | RAB6B |
| 36 | *RAP2B* | ENSG00000181467 | RAP2B |
| 37 | *RNLS* | ENSG00000184719 | RNLS |
| 38 | *RSPH1* | ENSG00000160188 | RSPH1 |
| 39 | *SLC37A1* | ENSG00000160190 | SLC37A1 |
| 40 | *SUB1* | ENSG00000113387 | SUB1 |
| 41 | *TBC1D22B* | ENSG00000065491 | TBC1D22B |
| 42 | *TMEM106C* | ENSG00000134291 | TMEM106C |
| 43 | *TMEM217* | ENSG00000172738 | TMEM217 |
| 44 | *TPM3* | ENSG00000143549 | TPM3 |
| 45 | *ZNF467* | ENSG00000181444 | ZNF467 |

**8. Prostate Cancer Classification System (PCS) PCS1-3 subtype-enriched genes**

Reference: You, S., ... & Freeman, M. R. (2016). Integrated classification of prostate cancer reveals a novel luminal subtype with poor outcome. Cancer research, 76(17), 4948-4958.

| | **PCS1 (86 genes)** | **PCS2 (123 genes)** | **PCS3 (219 genes)** |
|---|---|---|---|
| 1 | *BUB1* | *PPAP2A* | *ACTC1* |
| 2 | *KIF4A* | *AGR2* | *ACTG2* |
| 3 | *CCNA2* | *NKX3-1* | *ATP1A2* |
| 4 | *CENPE* | *EPCAM* | *RARRES2* |
| 5 | *CKS2* | *RAB3B* | *CXCL1* |
| 6 | *DLGAP5* | *ST6GAL1* | *PGM5* |
| 7 | *KIF2C* | *TMEFF2* | *TAGLN* |
| 8 | *BUB1B* | *AMD1* | *KCNAB1* |
| 9 | *CDK1* | *TOM1L1* | *SPON1* |
| 10 | *CDC6* | *ARG2* | *ANGPT1* |
| 11 | *CENPA* | *CACNA1D* | *DES* |
| 12 | *KIF23* | *GUCY1A3* | *DPT* |
| 13 | *CCNB1* | *UAP1* | *FOSB* |
| 14 | *CDC20* | *KLK3* | *GABRP* |
| 15 | *CENPF* | *F5* | *MYLK* |
| 16 | *HMMR* | *HLA-DMB* | *PPP1R12B* |
| 17 | *RRM2* | *KCNN2* | *NBL1* |
| 18 | *AURKA* | *ABCC4* | *DDR2* |
| 19 | *CCNB2* | *STK39* | *RARRES1* |
| 20 | *PTTG1* | *SH3RF1* | *RBP1* |
| 21 | *KNTC1* | *TARP* | *TGM4* |
| 22 | *KIF14* | *COL9A2* | *TPM2* |
| 23 | *ZWINT* | *DPP4* | *ADAMTS1* |
| 24 | *NUSAP1* | *GHR* | *CXCL13* |
| 25 | *CDCA5* | *MYO6* | *TNC* |
| 26 | *ASPM* | *COLEC12* | *NCAM1* |
| 27 | *MCM4* | *ACPP* | *ACTA2* |
| 28 | *KIAA0101* | *ALDH1A3* | *ANPEP* |
| 29 | *TPX2* | *ANK3* | *ATF3* |
| 30 | *UHRF1* | *DHCR24* | *COL4A6* |
| 31 | *DTL* | *DSC2* | *CRYAB* |
| 32 | *BIRC5* | *ERG* | *FOXF1* |
| 33 | *ECT2* | *F3* | *FLNA* |
| 34 | *FABP5* | *ACSL3* | *CXCL2* |
| 35 | *LMNB1* | *FMOD* | *ITGA5* |
| 36 | *TOP2A* | *GCNT1* | *ITGA7* |
| 37 | *TTK* | *GJB1* | *KRT17* |
| 38 | *TYMS* | *HPN* | *MAL* |
| 39 | *CDC45* | *KLK2* | *MYH11* |
| 40 | *PKMYT1* | *LCP1* | *PENK* |
| 41 | *MELK* | *TACSTD2* | *SERPINB5* |
| 42 | *KIF20A* | *MSMB* | *PMP22* |
| 43 | *CIT* | *MYBPC1* | *PTGDS* |
| 44 | *ESRP1* | *PGM3* | *S100A6* |

| | | | |
|---|---|---|---|
| 45 | *HJURP* | *PPP3CA* | *CXCL12* |
| 46 | *NCAPG* | *SORD* | *SMTN* |
| 47 | *CENPM* | *SPOCK1* | *SRD5A2* |
| 48 | *NETO2* | *TMPRSS2* | *TPM1* |
| 49 | *HES6* | *PLA2G7* | *ZFP36* |
| 50 | *EZH2* | *SLC4A4* | *NR4A3* |
| 51 | *UGT2B15* | *CLDN8* | *SRPX* |
| 52 | *ANLN* | *ZMPSTE24* | *ALDH1A2* |
| 53 | *NCAPG2* | *CRISP3* | *IER3* |
| 54 | *KIF15* | *PDLIM5* | *SOCS3* |
| 55 | *NUF2* | *IQGAP2* | *PDLIM7* |
| 56 | *ALB* | *PDIA5* | *PAGE4* |
| 57 | *AR* | *CUX2* | *MYL9* |
| 58 | *FOXM1* | *NEDD4L* | *SORBS1* |
| 59 | *HMGB2* | *SLC39A6* | *DKK1* |
| 60 | *KIF11* | *RDH11* | *LMOD1* |
| 61 | *STMN1* | *FAM198B* | *EFEMP2* |
| 62 | *MKI67* | *PLA1A* | *SPOCK3* |
| 63 | *TK1* | *FNIP2* | *FXYD6* |
| 64 | *PRC1* | *DNASE2B* | *SMOC1* |
| 65 | *CCNE2* | *ENPP5* | *PTRF* |
| 66 | *EXO1* | *ELOVL5* | *AOX1* |
| 67 | *TROAP* | *TRPM8* | *RND3* |
| 68 | *TACC3* | *STEAP4* | *ASPA* |
| 69 | *UBE2C* | *ACSM1* | *ATP2B4* |
| 70 | *UBE2T* | *TMEM178A* | *CFB* |
| 71 | *RACGAP1* | *OR51E1* | *BMP5* |
| 72 | *CDCA8* | *CREB3L4* | *SERPING1* |
| 73 | *CEP55* | *SLC38A11* | *C1S* |
| 74 | *PBK* | *REPS2* | *CAV1* |
| 75 | *MLF1IP* | *FBP1* | *CAV2* |
| 76 | *FAM111B* | *SLC30A4* | *CCND2* |
| 77 | *HOXC6* | *HOXB13* | *CES1* |
| 78 | *CDKN3* | *SLC27A2* | *CLU* |
| 79 | *CELSR3* | *KIAA1324* | *CNN1* |
| 80 | *SHMT2* | *TMEM45B* | *COL6A1* |
| 81 | *SPP1* | *ANXA3* | *COL6A2* |
| 82 | *RAD54L* | *ENTPD5* | *COL16A1* |
| 83 | *SPAG5* | *FOLH1* | *COX7A1* |
| 84 | *POLQ* | *HGD* | *CSRP1* |
| 85 | *HILPDA* | *NEFH* | *CYP3A5* |
| 86 | *HN1* | *NPY* | *CYP4B1* |
| 87 | | *PLA2G2A* | *CYP27A1* |
| 88 | | *RAB27B* | *DEFB1* |
| 89 | | *ACSM3* | *CFD* |
| 90 | | *SLC12A2* | *DPYSL3* |
| 91 | | *SOAT1* | *FBLN1* |
| 92 | | *TSPAN8* | *EFEMP1* |
| 93 | | *TM9SF2* | *FGFR2* |
| 94 | | *FAM189A2* | *FHL1* |
| 95 | | *TSPAN1* | *FHL2* |
| 96 | | *TMSB15A* | *FLNC* |
| 97 | | *AMACR* | *GABRE* |
| 98 | | *MPC2* | *GAS1* |
| 99 | | *SLC17A5* | *GSN* |
| 100 | | *STEAP1* | *GSTM1* |
| 101 | | *GPR160* | *GSTM2* |
| 102 | | *MMADHC* | *GSTM5* |
| 103 | | *CRYL1* | *GSTP1* |
| 104 | | *HSD17B11* | *ID1* |
| 105 | | *GOLM1* | *ID3* |
| 106 | | *CYP39A1* | *IGFBP6* |
| 107 | | *DHRS7* | *CYR61* |
| 108 | | *GALNT7* | *IL6* |
| 109 | | *DNAJC10* | *KCNJ8* |
| 110 | | *RBM47* | *KCNMB1* |
| 111 | | *CSGALNACT1* | *KRT5* |
| 112 | | *TDRD1* | *KRT13* |
| 113 | | *SMOC2* | *KRT15* |
| 114 | | *ZNF614* | *LAMA4* |

| | | | |
|---|---|---|---|
| 115 | | CWH43 | LAMB3 |
| 116 | | OR51E2 | LCN2 |
| 117 | | C15orf48 | LGALS1 |
| 118 | | TMTC4 | LTF |
| 119 | | SEC11C | MAOB |
| 120 | | GLYATL1 | MATN2 |
| 121 | | CPNE4 | MEIS1 |
| 122 | | FOLH1B | MEIS2 |
| 123 | | ZNF615 | MFAP4 |
| 124 | | | ROR2 |
| 125 | | | OGN |
| 126 | | | PCDH7 |
| 127 | | | PCP4 |
| 128 | | | SERPINF1 |
| 129 | | | FXYD1 |
| 130 | | | PLN |
| 131 | | | PRKCB |
| 132 | | | MASP1 |
| 133 | | | PTN |
| 134 | | | PYGM |
| 135 | | | S100A2 |
| 136 | | | S100A4 |
| 137 | | | CCL2 |
| 138 | | | CX3CL1 |
| 139 | | | SELE |
| 140 | | | SGCA |
| 141 | | | SLC2A5 |
| 142 | | | SLC14A1 |
| 143 | | | SMARCD3 |
| 144 | | | STAC |
| 145 | | | SVIL |
| 146 | | | TGFB1I1 |
| 147 | | | TGFB3 |
| 148 | | | TIMP2 |
| 149 | | | CLEC3B |
| 150 | | | TNS1 |
| 151 | | | TRIP6 |
| 152 | | | SCGB1A1 |
| 153 | | | VCL |
| 154 | | | RNF112 |
| 155 | | | ACOX2 |
| 156 | | | SPARCL1 |
| 157 | | | LTBP4 |
| 158 | | | PPAP2B |
| 159 | | | TP63 |
| 160 | | | AOC3 |
| 161 | | | PDE5A |
| 162 | | | HEPH |
| 163 | | | RCAN2 |
| 164 | | | EFS |
| 165 | | | SPEG |
| 166 | | | MRVI1 |
| 167 | | | WFDC2 |
| 168 | | | OLFM4 |
| 169 | | | FAXDC2 |
| 170 | | | ADIRF |
| 171 | | | RBPMS |
| 172 | | | EMILIN1 |
| 173 | | | LDB3 |
| 174 | | | FAM107A |
| 175 | | | FILIP1L |
| 176 | | | SCRG1 |
| 177 | | | PALLD |
| 178 | | | SYNM |
| 179 | | | VSIG2 |
| 180 | | | TRIM29 |
| 181 | | | KANK2 |
| 182 | | | KRT23 |
| 183 | | | CLIP3 |
| 184 | | | HSPB8 |

| | | | |
|---|---|---|---|
| 185 | | | *PCOLCE2* |
| 186 | | | *DKK3* |
| 187 | | | *HSPB7* |
| 188 | | | *PDZRN4* |
| 189 | | | *RASL12* |
| 190 | | | *ASB2* |
| 191 | | | *LIMS2* |
| 192 | | | *WFDC1* |
| 193 | | | *TMEM35* |
| 194 | | | *POPDC2* |
| 195 | | | *NDNF* |
| 196 | | | *C1orf54* |
| 197 | | | *FHOD3* |
| 198 | | | *CCDC3* |
| 199 | | | *CRISPLD2* |
| 200 | | | *C2orf40* |
| 201 | | | *TUBB6* |
| 202 | | | *C16orf45* |
| 203 | | | *NEXN* |
| 204 | | | *CHRDL1* |
| 205 | | | *MYOCD* |
| 206 | | | *PPP1R14A* |
| 207 | | | *PRKCDBP* |
| 208 | | | *AHNAK2* |
| 209 | | | *MRGPRF* |
| 210 | | | *PCAT4* |
| 211 | | | *HSPB6* |
| 212 | | | *TCEAL2* |
| 213 | | | *RBFOX3* |
| 214 | | | *DACT3* |
| 215 | | | *ANKRD35* |
| 216 | | | *SYNPO2* |
| 217 | | | *MSRB3* |
| 218 | | | *C11orf96* |
| 219 | | | *MIR143HG* |
