## Supplementary material for "Increasing Cancer Stemness Drives Prostate Cancer Progression, Plasticity, Therapy Resistance and Poor Patient Survival": Table S5

**Liu, Jamroze & Cortes et al.: Supplementary Table 5: Gene signature and Gene sets Information used in Gene Set Enrichment Analysis (GSEA)**

| No. | Gene Signature or Gene Set | Figures | Gene Set Enrichment Category | Gene Set Source: MSigDB, Literature (Pubmed ID) | No. of Genes in the Gene Set | Brief Description |
|---|---|---|---|---|---|---|
| 1 | Benporath_ES_2 | Fig. 5E | stemness | GSEA MSigDB, C2 (curated gene sets); Pubmed 18443585 | 40 | Genes overexpressed in human embryonic stem cells according to a meta-analysis of 8 profiling studies.<br>MSigDB standard name: Benporath_ES_2 |
| 2 | Bhattacharya_ESC | Fig. 5E | stemness | GSEA MSigDB, C2 (curated gene sets); Pubmed 15070671 | 84 | The 'stemnes' signature: genes up-regulated and common to 6 human embryonic stem cell lines tested.<br>MSigDB standard name: BHATTACHARYA_EMBRYONIC_STEM_CELL |
| 3 | Chandran_Metastasis_Up;<br>Chandran_Metastasis_Dn | Fig. 5D, 5E | Cancer aggressiveness | GSEA MSigDB, C2 (curated gene sets); Pubmed 17430594 | 209;<br>313 | Genes up-regulated in metastatic tumors from the whole panel of patients with prostate cancer;<br>Genes down-regulated in metastatic tumors from the whole panel of patients with prostate cancer.<br>MSigDB standard name: CHANDRAN_METASTASIS_UP; CHANDRAN_METASTASIS_DN |
| 4 | Lee_Metastasis_Up | Fig. 5E | Cancer aggressiveness | GSEA MSigDB, C2 (curated gene sets); Pubmed 18245461 | 16 | Components of RNA post-transcriptional modification machinery up-regulated in MDA-MB-435 cells (breast cancer) whose metastatic potential has been reduced by expression of NME1 [GeneID=4830]<br>MSigDB standard name: LEE_METASTASIS_AND_RNA_PROCESSING_UP |
| 5 | Wang_Metastasis_Up | Fig. 5E | Cancer aggressiveness | GSEA MSigDB, C2 (curated gene sets); Pubmed 15721472 | 22 | Genes whose expression in primary ER(+) [GeneID=2099] breast cancer tumors positively correlates with developing distant metastases.<br>MSigDB standard name: WANG_METASTASIS_OF_BREAST_CANCER_ESR1_UP |
| 6 | Wallace_PCa_Cancer_Up | Fig. 5E | Cancer aggressiveness | GSEA MSigDB, C2 (curated gene sets); Pubmed 18245496 | 20 | Genes up-regulated in prostate tumor vs normal tissue samples. MSigDB standard name: WALLACE_PROSTATE_CANCER_UP |
| 7 | Tomlins_PCa_Cancer_Up;<br>Tomlins_PCa_Cancer_Dn | Fig. 5D, 5E | Cancer aggressiveness | GSEA MSigDB, C2 (curated gene sets); Pubmed 17173048 | 40;<br>40 | Genes up-regulated in prostate cancer vs benign prostate tissue, based on a meta-analysis of five gene expression profiling studies.<br>MSigDB standard name: TOMLINS_PROSTATE_CANCER_UP; TOMLINS_PROSTATE_CANCER_DN |
| 8 | Rhodes_undifferentiated_Cancer | Fig. 5E | Cancer aggressiveness | GSEA MSigDB, C2 (curated gene sets); Pubmed 15184677 | 69 | Genes commonly up-regulated in undifferentiated cancer (more aggressiveness) relative to well-differentiated (less aggressiveness) cancer, based on the meta-analysis of the OncoMine gene expression database.<br>MSigDB standard name: RHODES_UNDIFFERENTIATED_CANCER |
| 9 | Liu_Prostate_Cancer_Dn | Fig. 5D | Cancer aggressiveness | GSEA MSigDB, C2 (curated gene sets); Pubmed 16618720 | 493 | Genes down-regulated in prostate cancer samples vs. benign tissue.<br>MSigDB standard name: LIU_PROSTATE_CANCER_DN |
| 10 | Smid_Breast_Cancer_Dn | Fig. 5D | Cancer aggressiveness | GSEA MSigDB, C2 (curated gene sets); Pubmed 18451135 | 485 | Genes down-regulated in aggressive subtypes of breast cancer vs. normal-like subtype of breast cancer.<br>MSigDB standard name: SMID_BREAST_CANCER_NORMAL_LIKE_UP |
| 11 | DNA_REPAIR | Fig. 5E | GSEA Hallmark Pathway | GSEA MSigDB, Hallmark; Pubmed 26771021 | 150 | Genes involved in DNA repair.<br>MSigDB standard name: HALLMARK_DNA_REPAIR |
| 12 | E2F_TARGETS | Fig. 5E | GSEA Hallmark Pathway | GSEA MSigDB, Hallmark; Pubmed 26771021 | 200 | Genes encoding cell cycle related targets of E2F transcription factors.<br>MSigDB standard name: HALLMARK_E2F_TARGETS |
| 13 | IFN_GAMMA_RESPONSE | Fig. 5D | GSEA Hallmark Pathway | GSEA MSigDB, Hallmark; Pubmed 26771021 | 200 | Genes up-regulated in response to IFNG.<br>MSigDB standard name: HALLMARK_INTERFERON_GAMMA_RESPONSE |
| 14 | INFLAMMATORY_RESPONSE | Fig. 5D | GSEA Hallmark Pathway | GSEA MSigDB, Hallmark; Pubmed 26771021 | 200 | Genes defining inflammatory response.<br>MSigDB standard name: HALLMARK_INFLAMMATORY_RESPONSE |
| 15 | MTORC1_SIGNALING | Fig. 5E | GSEA Hallmark Pathway | GSEA MSigDB, Hallmark; Pubmed 26771021 | 200 | Genes up-regulated through activation of mTORC1 complex.<br>MSigDB standard name: HALLMARK_MTORC1_SIGNALING |
| 16 | MYC_TARGETS_V1 | Fig. 5E | GSEA Hallmark Pathway | GSEA MSigDB, Hallmark; Pubmed 26771021 | 200 | A subgroup of genes regulated by MYC - version 1 (v1). MSigDB standard name: HALLMARK_MYC_TARGETS_V1 |
| 17 | MYC_TARGETS_V2 | Fig. 5E | GSEA Hallmark Pathway | GSEA MSigDB, Hallmark; Pubmed 26771021 | 58 | A subgroup of genes regulated by MYC - version 2 (v2). MSigDB standard name: HALLMARK_MYC_TARGETS_V2 |
| 18 | TGF_BETA_SIGNALING | Fig. 5D | GSEA Hallmark Pathway | GSEA MSigDB, Hallmark; Pubmed 26771021 | 54 | Genes up-regulated in response to TGFB1.<br>MSigDB standard name: HALLMARK_TGF_BETA_SIGNALING |
| 19 | TNFA_SIGNALING_VIA_NFKB | Fig. 5D | GSEA Hallmark Pathway | GSEA MSigDB, Hallmark; Pubmed 26771021 | 200 | Genes regulated by NF-kB in response to TNF.<br>MSigDB standard name: HALLMARK_TNFA_SIGNALING_VIA_NFKB |
| 20 | You_PCA_subtype: PCS1;<br>You_PCA_subtype: PCS3 | Fig. 5H, 5I, 5J, 5K | PCa subtypes | Pubmed 27302169 | 86;<br>219 | Integrated classification of prostate cancer based on human PCa transcriptome profiles from a large virutal cohort (n=1,321) developed a novel classification system consisting of 3 distinct subtypes (named PCS1, PCS2, PCS3). PCS1 and PCS2 tumors reflect luminal subtypes, while PCS3 represents a basal subtype. PCS1 tumors progress more rapidly to metastatic disease in comparison to PCS2 or PCS3.<br>Wilcoxon rank-sum test and subsequent false discovery rate (FDR) correction with Storey's method (25) were employed to identify differentially expressed genes between the subtypes. Genes were selected with FDR<0.001 and fold change ≥1.5, resulting in 428 SEGs. Among 428 subtype enriched genes, 86 for PCS1, 123 for PCS2, and 219 for PCS3. |
| 21 | Tang_mCRPC-AR-dependent;<br>Tang_mCRPC-NE-subtype | Fig. 5L, 5M | mCRPC subtypes | Pubmed 35617398 | 93;<br>93 | Integrated classfication of castration-resistant prostate cancer based on Chromatin (accessbility) profiles (ATAC-seq), transcriptomic profiles (RNA-seq), genomic alterations (DNA sequncing) in 40 metastatic prostate cancer models, including 22 organoids, 6 PDXs, 7 cell lines, and 5 derived CRPC cell lines from Park et al Science 2018, the study revealed four subtypes using ATAC-seq clusering, AR-dependent, neuroendocrine, and two AR-negative/low groups (Wnt-dependent, stem cell-like). Gene signatures of the four epigenetically defined subgroups were generated using RNA-seq data. |
| 22 | Coleman_Cell cycle progression | Fig. 5N, 5O | Cell Cycle Progression | Pubmed 35552660; 21310658 | 31 | This gene signature is curated in a published study Pubmed 21310658, where the authors initially selected 126 cell cycle progression (CCP) genes from the Gene Expression Omnibus database and tested their performance with RNA extracted from 96 commercially available FFPE prostate tumour sections (Asterand, Detroit, MI, USA) obtained from anonymous patients and subsequently the authors selected genes for inclusion in the signature on the basis of their correlation with the mean expression of the entire set of CCP genes. The final signature consisted of 31 CCP genes (FOXM1, CDC20, CDKN3, CDC2, KIF11, KIAA0101, NUSAP1, CENPF, ASPM, BUB1B, RRM2, DLGAP5, BIRC5, KIF20A, PLK1, TOP2A, TK1, PBK, ASF1B, C18orf24, RAD54L, PTTG1, CDCA3, MCM10, PRC1, DTL, CEP55, RAD51, CENPM, CDCA8, and ORC6L). These highly correlated genes were used to provide a robust and highly reproducible measurement of cell proliferation and were not intended to capture information related to other factors (eg, invasive potential). The signature was assessed retrospectively in a cohort of patients from the USA who had undergone radical prostatectomy, and in a cohort of randomly selected men with clinically localised prostate cancer diagnosed by use of a transurethral resection of the prostate (TURP) in the UK who were managed conservatively. The CCP score was predictive of outcome in both cohorts. The authors also suggested that Expression of CCP genes is higher in actively growing cells, and presumably by measuring the expression of CCP genes, we are able to indirectly measure the growth rate and inherent aggressiveness of the tumour. |
| 23 | Alumkal_Lineage plasticity risk | Fig. 5P, 5Q | Lineage Plasticity Risk in CRPC | Pubmed 36109521 | 14 | This gene signature is curated in a published study using the transcriptomes of matched biopsies from men with metastatic CRPC obtained prior to the androgen receptor (AR) signaling inhibitor Enzalutamide (enza) treatment and at progression (n = 21). To determine if any of the progression tumors in the cohort underwent lineage plasticity after enza, the authors determined the Aggarwal cluster and Labrecque classifier designation. Twelve of 21 matched pairs did not change their Aggarwal cluster designation. However, three of the 21 progression tumors (hereafter referred to as converters) had gene expression profiles consistent with cluster 2, suggesting enza-induced conversion to an alternate lineage. By identifying genes significantly upregulated in the baseline tumors from converters vs. non-converters, the authors identified a 14-gene signature highly activated in the three baseline tumors from converters. This signature indicates the susceptibility to transcriptional conversion and lineage plasticity. |
